# Hi-cGAN: Prediction of Hi-C interaction matrices with conditional generative adversarial networks

**DOI:** 10.64898/2026.08.30.748103

**Authors:** Ralf Krauth, Anup Kumar, Joachim Wolff

## Abstract

**Background:** The three-dimensional organization of the genome is a fundamental aspect of its function and regulation. High-throughput chromosome conformation capture techniques, such as Hi-C, have revolutionized our understanding of spatial genome organization. However, 3C-based methods are resource-intensive and technically demanding. This has driven the development of computational approaches for predicting Hi-C interaction matrices. Hi-cGAN, a novel approach based on conditional generative adversarial networks, offers a computational alternative to extensive wet-lab work by predicting Hi-C interaction matrices. This computational approach contributes to a broader exploration and understanding of genome architecture.

**Findings:** The network pairs a convolutional generator with a convolutional discriminator, evaluated across bin sizes, inputs and cell types. It predicts a whole genome as a cool file at bin sizes from 2 to 25 kb, where Akita, C.Origami and Epiphany emit fixed windows of 1 Mb, 2 Mb and 990 kb. With the input chosen on a validation chromosome, agreement approaches Epiphany’s and stays below the sequence-based C.Origami and Akita: over Akita’s 411 held-out windows the mean correlation is 0.238 against 0.506. Boundary and loop calls agree less closely, placing the maps at the domain scale.

**Conclusions:** Chromatin factor occupancy determines a substantial part of contact structure, and two tracks capture most of it. The most informative track depends on the resolution: CTCF and the cohesin subunits at 5 to 10 kb, active histone marks at 25 kb. Transfer to an unseen cell type costs about 0.12 SCC, and which method leads depends on the measure.

**Key points:**

- Hi-cGAN predicts Hi-C interaction matrices with a conditional generative adversarial network from chromatin factor occupancy alone, without DNA sequence.
- The same architecture learns raw contact counts, genomic-distance z-scores and observed/ expected normalized data.
- On held-out data Hi-cGAN reaches matrix-level agreement slightly below Epiphany’s, when the input configuration is chosen on a validation chromosome rather than the chromosomes scored, and well below C.Origami’s and Akita’s, which read DNA sequence. Which method is better depends on the measure used, and boundary and loop recovery does not follow the matrix-level scores.
- A ridge regression on the same single track reaches 0.691 against Hi-cGAN’s 0.715 on HiCRep, while its predicted matrix shows stripes but no domains, so that measure alone does not establish what the network contributes.
- Bin sizes of 2 to 25 kb are evaluated and freely configurable, whole chromosomes are predicted, and the result is written as a cool file, whereas the sequence-based methods emit a single fixed window.

## 1 Introduction

High-throughput chromosome conformation capture, in particular Hi-C, is the standard technique for studying the three-dimensional architecture of the genome Lieberman-Aiden et al. (2009), and has shaped current understanding of chromatin organization and its role in transcription, replication and repair Bonev and Cavalli (2016). It established the features that structure the genome: A/B compartments Lieberman-Aiden et al. (2009); Schwarzer et al. (2017), topologically associating domains (TADs) Dixon et al. (2012); Nora et al. (2012) and DNA loops Rao et al. (2014). Because the assay is costly, computational prediction of contact matrices has developed quickly, and the approaches differ mainly in their input data.

Polymer simulations model DNA as beads on a string and take spatial constraints from ChIP-seq of chromatin factors Brackley et al. (2016); MacPherson et al. (2018) or from chromatin states derived from them Di Pierro et al. (2017); Qi and Zhang (2019). The matrices they produce bear little resemblance to measured Hi-C. Farré and Blanchette estimate contact probabilities from chromatin states called on DamID signal Farré and Emberly (2018); Greil et al. (2006), which identifies highly interacting regions without reproducing a Hi-C map.

Random forests predict loops as a binary call, in 3DEpiLoop Al Bkhetan and Plewczynski (2018) and Lollipop Kai et al. (2018). HiC-Reg Zhang et al. (2019) extends the same family to real-valued matrices, using chromatin features and genomic distance as decision criteria. Decision trees, logistic regression and neural networks have been compared for predicting TAD boundaries from histone modifications and CTCF Martens et al. (2020), and a one-dimensional convolutional filter over ChIP-seq tracks predicts complete Hi-C submatrices Farré et al. (2018).

A second family reads DNA sequence. SPEID predicts promoter-enhancer interactions with convolutional and recurrent layers Singh et al. (2019), as do Rambutan Schreiber et al. (2017) and Akita Fudenberg et al. (2020); DeepC transfers a sequence model trained on ChIP-seq tracks Schwessinger et al. (2019), and oligomer distance histograms with a support vector machine predict 5C interactions Nikumbh and Pfeifer (2017). Epiphany predicts cell-type-specific contact maps from five functional tracks with convolutional and bidirectional LSTM layers and an optional adversarial term Yang et al. (2023). C.Origami combines sequence with CTCF occupancy and chromatin accessibility in one encoder, for in silico screening in which the sequence input is perturbed and the predicted map inspected Tan et al. (2023).

We present a conditional generative adversarial network that predicts Hi-C matrices from chromatin factor occupancy. Adversarial training is not new to this problem: Epiphany reads functional tracks and offers an adversarial term of its own Yang et al. (2023). The difference is what the discriminator is shown. In Epiphany’s released implementation the discriminator takes a single-channel contact map and returns one scalar for it, so it classifies whether a map is a Hi-C map. Here the discriminator receives the chromatin tracks alongside the matrix and scores overlapping patches, so it classifies whether a map is plausible *for this input*. The same architecture learns raw contact counts, genomic-distance z-scores or observed/expected normalized data, is not restricted to a fixed window, and writes whole chromosomes to a cool file. The intended use is domain-scale description of a cell type for which chromatin tracks exist and Hi-C does not; the evaluation below shows that loop and boundary calling are not supported uses.

## 2 Methods

The implementation of a conditional generative adversarial network (cGAN), named Hi-cGAN, for Hi-C prediction is motivated by the performance of GANs in image processing tasks, such as image synthesis. In this scenario, chromatin features are considered as descriptions from which the target Hi-C sub-matrices are to be synthesized.

The input data must first be transformed to conform to the input shape required by the underlying image synthesis network. To facilitate this, a convolutional neural network (CNN) is employed as an embedding network for the chromatin features. The advantage of using CNNs is their ability to preserve localization information contained in the chromatin feature data, which is crucial for accurately predicting spatial relationships in Hi-C matrices.

### 2.1 Hi-cGAN architecture

Hi-cGAN follows the conditional GAN formulation: a generator *G*(*x, z*) and a discriminator *D*(*x, y*) play the minimax game of Equations 1 and 2, in which **x** are the chromatin features, **y** the measured Hi-C matrix, **z** random noise and E the expected value Isola et al. (2017). The generator learns to synthesise a matrix from chromatin features and noise; the discriminator learns to distinguish it from a measured matrix.

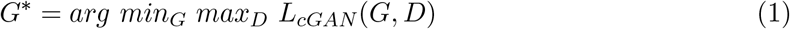

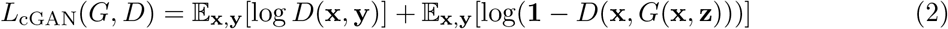

Both networks follow pix2pix Isola et al. (2017), each extended by a CNN that turns the chromatin input into the grayscale image the network expects. The generator is a U-Net Ronneberger et al. (2015), whose encoder-decoder structure with skip connections retains features at several levels of abstraction; it receives the ChIP-seq data and produces the predicted submatrix through convolution and upsampling. The discriminator is a PatchGAN Isola et al. (2017), and it is conditional: it receives the chromatin input alongside the matrix it classifies, so its output is a function of the pair rather than of the matrix alone.

The stochastic input **z** of Figure 1 and of the loss equations below is realised as it is in pix2pix, which supplies no noise vector and instead introduces the randomness via dropout Isola et al. (2017). Here that is dropout at rate 0.5 in the first decoder blocks, three of them at a window of 256 and 512 and two at the smaller windows, where the decoder is shortened. In pix2pix, dropout is left on at test time, so the generator is stochastic; here it is disabled at inference, so a trained model returns the same matrix for the same input on every run and every prediction reported below is deterministic.

**Figure 1:**
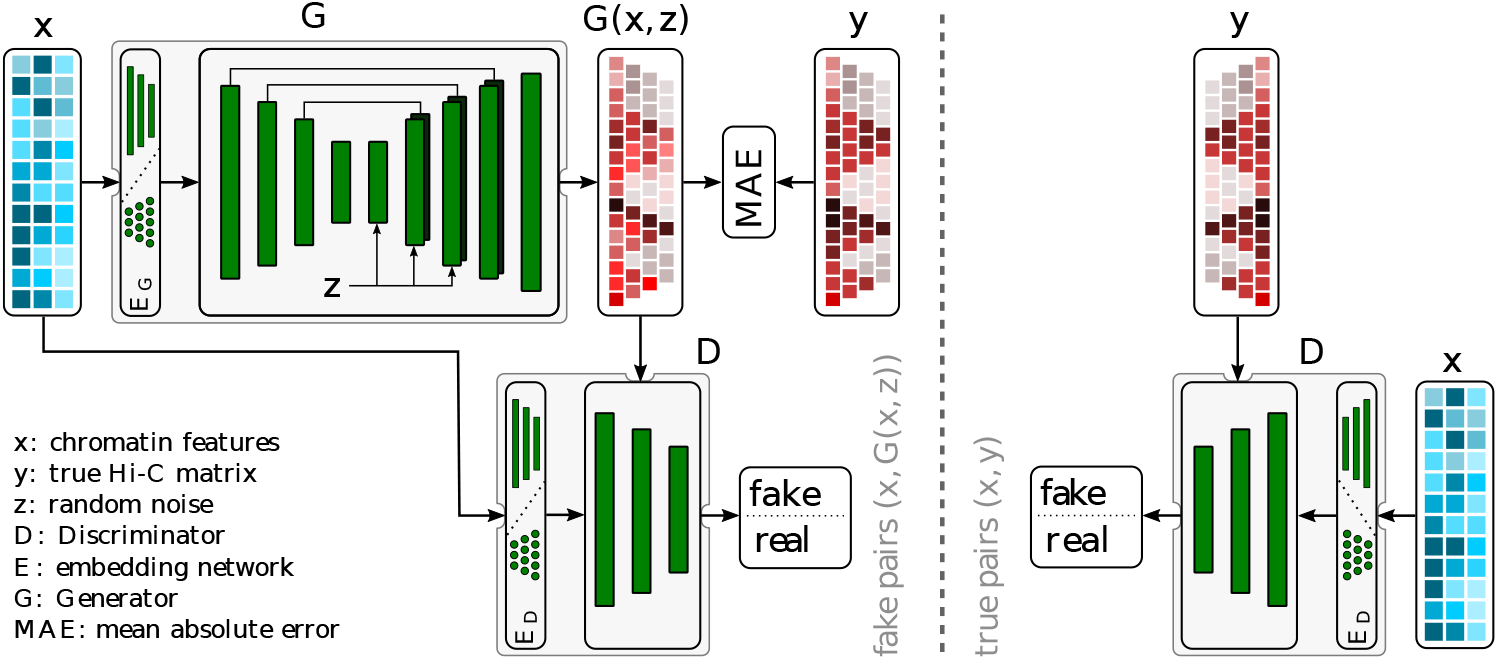
Architecture of the Hi-cGAN approach: The model employs a conditional GAN, where the input data *x* is derived from one-dimensional chromatin features. These are based on ChIP-seq data primarily of histone variants. Real Hi-C matrices *y* are used for training the discriminator to distinguish between generated and true Hi-C matrices.

The generator loss function minimized while training the generator is similar to the pix2pix approach, but with an additional total variation (TV) loss function to reduce noise, see equation 3. In this context, *L_MAE_* refers to the mean absolute error (L1 loss) between the synthetic output *G*(**x, z**) and the actual Hi-C submatrices *y* (represented as grayscale images). *L_d_*, described in equation 4, is the binary cross-entropy loss for the discriminator between its inputs **s** and **r**, which classifies input matrices as real or generated. *L_T V_* is a total variation loss, and *σ* the sigmoid function. In this context, the discriminator processes batches containing image patches with a shape of (*n, n*), i.e., tensors of shape (*bs, n, n,* 1), and **1** also represents a tensor of shape (*bs, n, n,* 1), filled entirely with ones. The architecture of the generator is fully detailed in Supplementary Section S3.

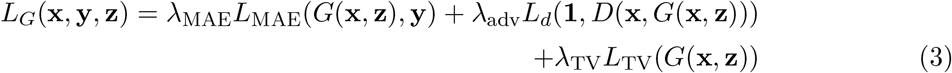

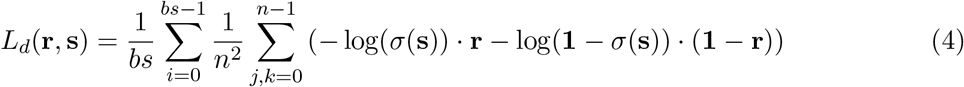

The PatchGAN loss of the discriminator is defined as Isola et al. (2017):

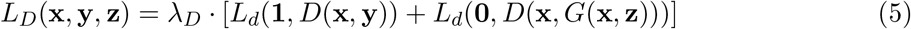

Here *λ_D_*is the scalar of Isola et al. Isola et al. (2017) that moderates the discriminator’s learning rate relative to the generator’s, and **1** and **0** are tensors filled with ones and zeros. Both architectures are given in full in Supplementary Section S3.

The weights are those of pix2pix where pix2pix defines them: *λ*_MAE_ = 100, *λ*_adv_ = 1 and *λ_D_* = 0.5. The total variation term is the one addition, and it is set to *λ*_TV_ = 10*^−^*^10^, which is small enough that it does not contribute; it is retained in the equation because it is present in the implementation and can be raised; the results below do not depend on it. Both networks use Adam with *β*_1_ = 0.5, at a learning rate of 2*×*10*^−^*^5^ for the generator and 10*^−^*^6^ for the discriminator, the latter lower so that the discriminator does not outpace the generator. No weight was tuned per configuration.

### 2.2 Input sample generation

The input is one-dimensional ChIP-seq signal, mostly of histone variants (Table S1). CTCF marks TAD and loop anchors Rao et al. (2014), and histone modifications report activating or repressing states and thereby chromatin accessibility Henikoff and Smith (2015); Cheung and Lau (2005). Multiple ChIP-seq, respectively bigwig, files can be combined as input, and the signal is not restricted to ChIP-seq: any technique serves as long as its output is a bigwig file.

To be processable by Hi-cGAN’s generator and discriminator networks, the input data needs to be transformed. Each of the *n* one-dimensional data sets is binned into bins of size *b_feature_*, which is equal to the resolution of the Hi-C matrix used for training (e.g. 5 kb). This is done by taking the mean signal value within each bin. For a chromosome size of *cs* and *n* chromatin features, this yields a *l_feature_ × n* matrix named chromatin feature array. We then normalize each column of the chromatin feature array to value range [0, 1].

From the chromatin feature array, subarrays of dimension 3*w × n* are cut in a sliding-window fashion, where *w* is the window size. The window size defines the size of the cutout of the Hi-C matrix. Each submatrix is associated with a left and right flanking region, similar to the approach of Farré and Blanchette Farré and Emberly (2018), see Figure 2. The sub-matrix *G* gets assigned with the left flanking region *A*, the main region *B*, and the right flanking region *C* from the chromatin feature array. In the provided example in Figure 2, a total of five training samples can be generated.

**Figure 2:**
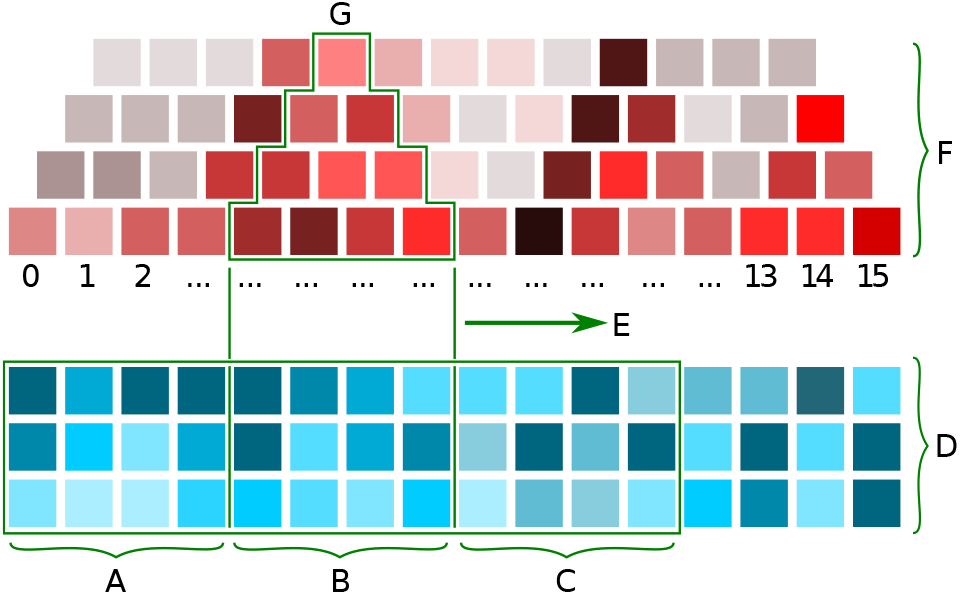
Input sample generation: Each sub-matrix of window size *w*, here *w* = 4 (letter G), gets associated with a left flanking region (A), main region (B), and a right flanking region (C) associated. The chromatin feature array has 3 ChIP-seq samples. In total, five training samples can be extracted. E is the sliding direction to generate the samples, and F is the Hi-C matrix at the diagonal rotated by 45 degrees.

Prediction generates samples the same way. The network emits a *w × w × b* tensor, where *b* is the number of generated feature windows; the *b* layers are summed and each value divided by the number of windows contributing to it, which away from the two corners of the matrix is *w − d* for a bin at distance *d* from the diagonal. The overlapping submatrices are then added and averaged in the same way to give the chromosome matrix. The first and last *w* bins on the main diagonal cannot be predicted, because of the flanking regions. The predicted values can be scaled back to the regular Hi-C value range and are stored as a *cooler* file Abdennur and Mirny (2020).

### 2.3 Input data embedding

The generator of the pix2pix network requires grayscale images of dimension (*w, w,* 1) as input, where *w* is the window size and 1 is the color channel. To transform the (3*w, n*) chromatin feature array, where *w* is the window size and *n* is the number of chromatin features, into the required shape, we used a CNN.

The embedding network has eight blocks of a 1D convolution, batch normalization and a leaky ReLU Maas et al. (2013), which keeps the gradient non-zero for negative inputs and so avoids dead units, and a ninth convolution that fixes the output shape. The full setup is given in Supplementary Section S3.

## 3 Results

To investigate the performance of Hi-cGAN, multiple approaches are pursued. First, the number of training epochs is examined; next, the window size; then the predictive value of the individual histone marks and structure-forming proteins; then the accuracy of the predicted matrices and the comparison against HiC-Reg Zhang et al. (2019), Epiphany Yang et al. (2023), C.Origami Tan et al. (2023) and Akita Fudenberg et al. (2020); last, the ability of trained models to predict other cell types. The methods are compared with the Pearson correlation stratified by genomic distance, the stratum-adjusted correlation coefficient of HiCRep Yang et al. (2017), GenomeDISCO Ursu et al. (2018), HiC-Spector Yan et al. (2017) and the agreement of insulation scores and the boundaries called from them Crane et al. (2015).

The experiments use the Hi-C matrices of Rao et al. Rao et al. (2014) of different cell types(GM12878, HMEC, HUVEC, IMR-90 and K562) and, as input, ChIP-seq data of histone modifications (H3K4me1, H3K4me2, H3K4me3, H3K9ac, H3K9me3, H3K27ac, H3K27me3, H3K36me3, H3K79me2 and H4K20me1) and of CTCF and the cohesin subunits RAD21 and SMC3, together with DNase accessibility. CTCF, RAD21 and SMC3 were chosen because they define chromatin structure, creating loops and domain boundaries Rao et al. (2014, 2017), and the histone marks because they report active and repressed chromatin Henikoff and Smith (2015); Cheung and Lau (2005). A nearly identical set underlies HiC-Reg Zhang et al. (2019), Epiphany Yang et al. (2023) reads five of these tracks, and C.Origami Tan et al. (2023) reads CTCF with chromatin accessibility.

### Training and prediction split

Training uses the odd chromosomes 1, 3, 5, 7, 9, 11, 13, 15, 17 and 21, chromosome 19 is the validation chromosome, and the twelve remaining chromosomes, the even ones and X, are test data. The split is by parity rather than by any property of the chromosomes, so the assignment cannot be influenced by how well a chromosome turns out to be predicted, and the held-out set spans a range of lengths and gene densities.

### Training set size

The Hi-cGAN sampler slides by one bin, so the number of training positions is large. At 25 kb with a window of 64 the ten training chromosomes offer 54,508 sliding positions, but only 857 non-overlapping windows fit into the same 55,148 target bins. At 10 kb with a window of 512 the figures are 125,702 positions and 251 non-overlapping windows, and at 5 kb 266,760 and 526.

### 3.1 Number of training epochs

The training length was tested by training the fourteen single-factor models at 25 kb with a window of 64 for 1000 epochs, with a checkpoint every ten (Figure 3). Averaged over the fourteen factors, held-out SCC is 0.699 at epoch 40, 0.704 at epoch 100 and 0.714 at epoch 1000, so accuracy is still rising, but by 0.010 over nine hundred further epochs. Choosing the best epoch per factor on the test set would raise SCC by 0.015 on average and 0.028 at most, a gain obtainable only through test-set leakage. Every model in this study is read at epoch 100.

**Figure 3:**
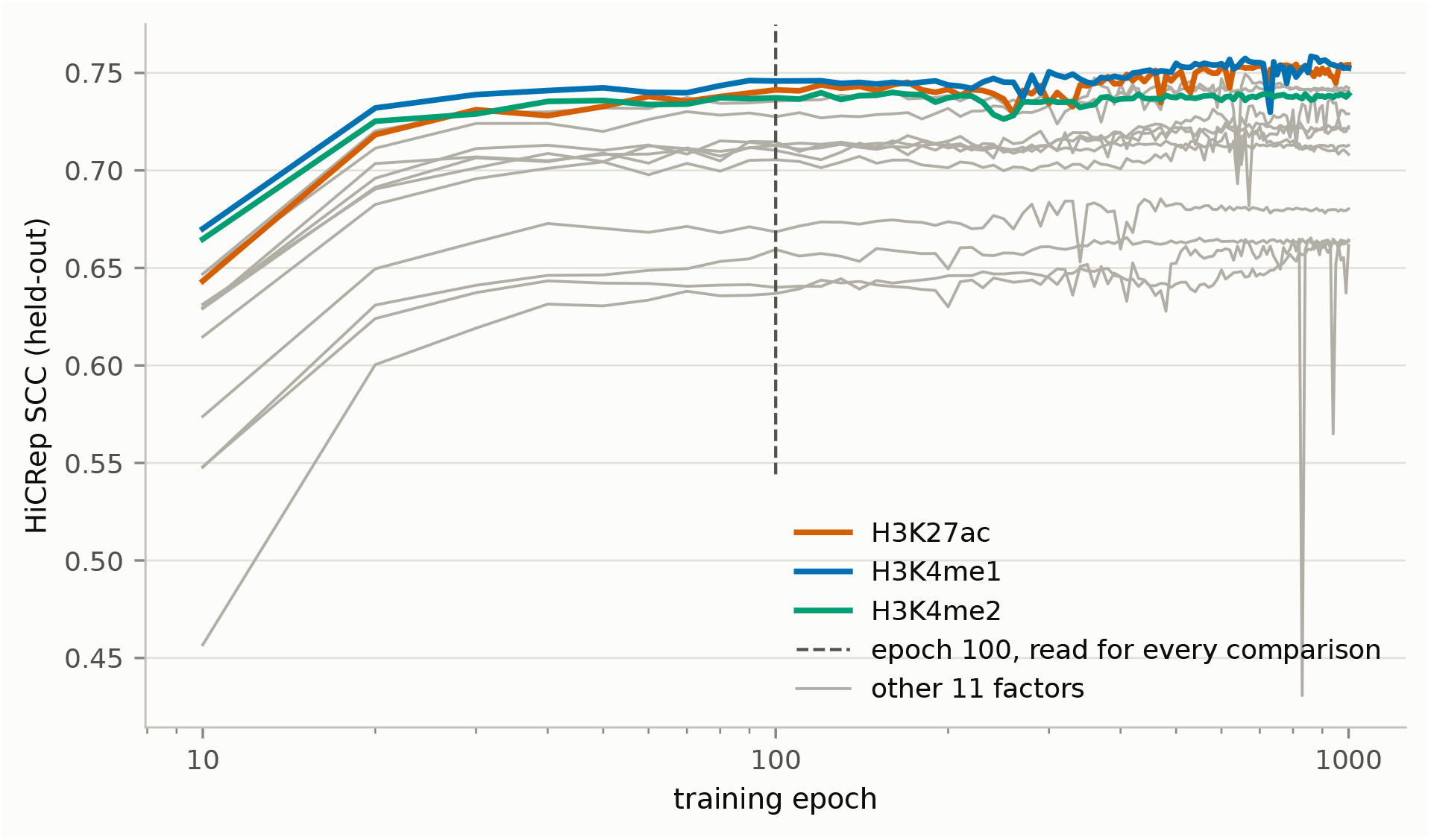
Accuracy over 1000 training epochs. Held-out SCC against training epoch on a logarithmic axis for the fourteen single-factor models at 25 kb with a window of 64, 100 checkpoints each. The three factors that rank highest at epoch 100 are coloured, the remaining eleven are grey, and epoch 100, the checkpoint used throughout this study, is marked.

### 3.2 Window sizes

Window size *w* sets both the regional context available to the model and the maximum genomic distance it can predict, *w* bins. To test the impact of the windows size, models at 25kb, 10kb and 5kb with a single protein input were trained. All read at epoch 100 and scored to 1 Mb. At 25 kb the smallest window is the most accurate for every one of the fourteen factors, with held-out SCC averaging 0.703 at *w* = 64, 0.645 at 128 and 0.526 at 256. At 10 kb the ordering is the same, again for every factor: 0.642 at *w* = 128, 0.586 at 256 and 0.502 at 512. At 5 kb the smallest window loses instead, 0.446 at *w* = 128 against 0.572 at 256 and 0.538 at 512, with 256 the best window for thirteen of the fourteen factors. The pattern follows the reach: the best window at each bin size is the smallest one that still covers the 1 Mb over which the maps are scored, 1.6 Mb at 25 kb with *w* = 64 and 1.28 Mb at 10 and 5 kb with *w* = 128 and 256, whereas *w* = 128 at 5 kb reaches only 0.64 Mb (Figure 5).

While the SCC favours the smaller windows, the later predictions are made with the larger ones, whose greater reach covers features at distances the small windows cannot represent. Figure 4 shows the same locus predicted at the three 10 kb windows: interactions beyond 1.28 Mb are absent at *w* = 128 and beyond 2.56 Mb at 256, and only the window of 512 carries the structure up to 5.12 Mb. The same view at 25 and 5 kb is Supplementary Figures S23 and S24.

**Figure 4:**
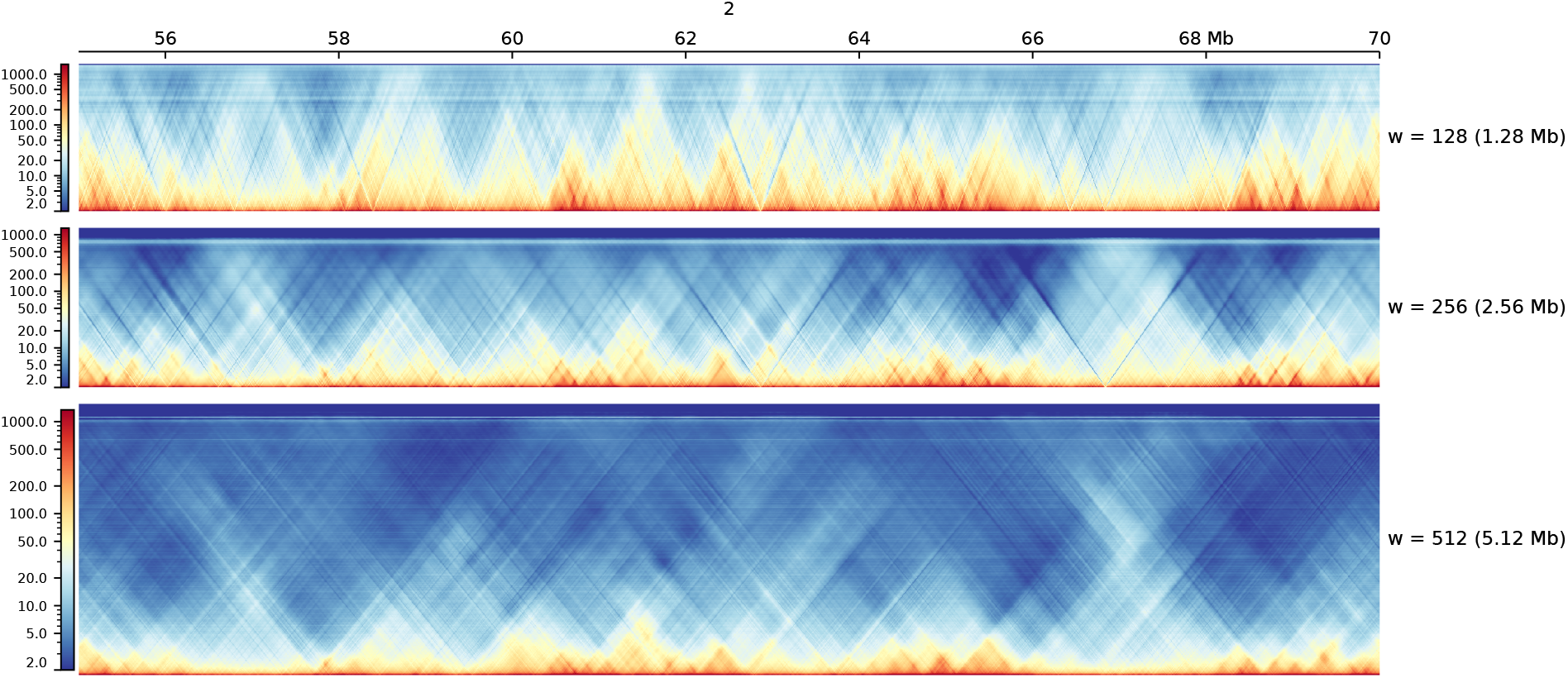
Prediction depth of the three windows at 10 kb. Chromosome 2, 55–70 Mb, predicted from CTCF at windows of 128, 256 and 512 bins and drawn by pyGenomeTracks at a shared scale per megabase. The height of each track is the maximum genomic distance that window can predict, 1.28, 2.56 and 5.12 Mb.

### 3.3 Chromatin factor selection

We next ask which chromatin factor, alone or in combination with others, gives the most accurate prediction at each bin size. Testing every combination is not feasible: fourteen factors give 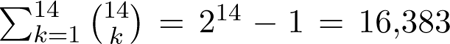 possible input sets. Training a single set for 100 epochs on an Nvidia A100 card takes 7.0 h at 25 kb with a window of 256, so the full search would take about 13 years on one card at this bin size alone, and far longer at 10 and 5 kb.

We therefore use forward selection. The fourteen factors are trained one at a time and ranked by their held-out SCC; the best factor is paired with each of the next-ranking factors, the best pair is extended by a third in the same way, and the search stops when adding a factor no longer improves the mean.

At 25 kb, with a window of 256, the fourteen single factors reach between 0.412 and 0.581 held-out SCC, with H3K27ac the best. Paired with a second factor, H3K27ac with CTCF wins at 0.586, and every third factor lowers it again, the best of the triples reaching 0.575, so the selection stops at two factors. At 10 kb, with a window of 512, the single factors span 0.412 to 0.574, led by CTCF. CTCF with H3K4me2 reaches 0.623, the largest gain from a second factor at any bin size, and the best triple falls to 0.563. At 5 kb, with the same window, the single factors span 0.439 to 0.615, led by RAD21. RAD21 with H3K4me3 reaches 0.643, adding SMC3 holds it at 0.643, and a fourth factor lowers it to 0.619. Accuracy therefore peaks at two or three well-chosen tracks: the tracks are strongly correlated with one another, so a further one adds parameters faster than information, while the number of training samples is fixed by the genome (Supplementary Table S14).

Which factor leads depends on the resolution: at 5 and 10 kb CTCF and the cohesin subunits RAD21 and SMC3 lead in every configuration tested, at 25 kb the active marks H3K4me1, H3K4me3 and H3K27ac, with CTCF at rank five to eight. This follows what a map contains at each scale: at 5 to 10 kb the resolvable features are the loop anchors and TAD boundaries that CTCF and cohesin create through loop extrusion Rao et al. (2014, 2017), at 25 kb compartment organization, which tracks transcriptional activity Lieberman-Aiden et al. (2009); Schwarzer et al. (2017). Within each family the members mark the same sites and are nearly interchangeable, cohesin being retained where CTCF halts it. This is also why the winning pairs combine an architectural track with a histone mark rather than two architectural ones: at 10 kb, CTCF with SMC3 reaches 0.600 and CTCF with RAD21 0.565 against the 0.623 of CTCF with H3K4me2, the second architectural track largely repeating what the first already provides, while the histone mark adds the activity signal (Figure 5). The complete results, including the per-chromosome distributions of every factor, are reported in Supplementary Sections S11 to S14.

**Figure 5:**
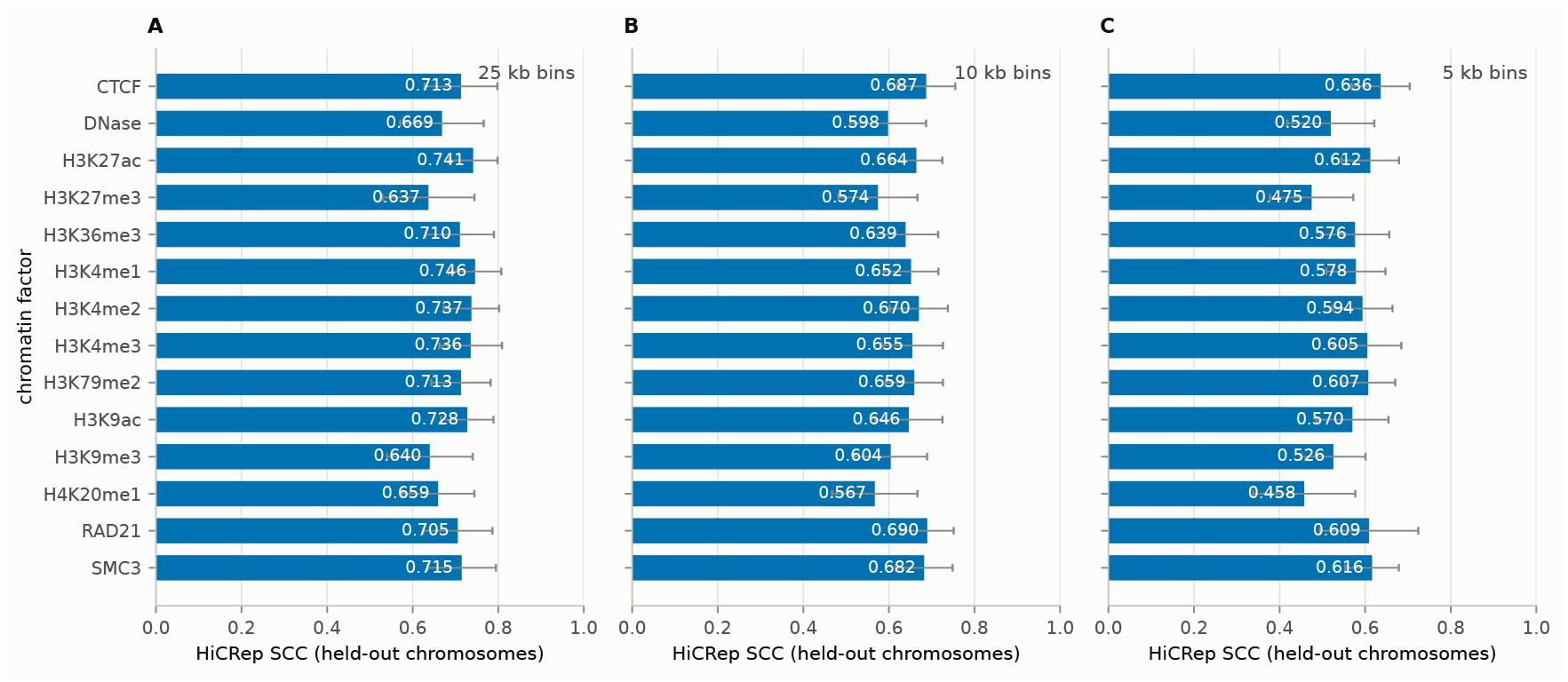
Single-factor accuracy across configurations. Held-out SCC for each of the fourteen chromatin factors used alone, at every bin size and window size in the sweep. The spread between factors is narrower than the spread between configurations of the same factor, and the ranking changes with the bin size: CTCF and the cohesin subunits lead at 5 and 10 kb, the active histone marks at 25 kb.

### 3.4 Prediction of raw Hi-C matrices

To evaluate the accuracy of complete predicted matrices, the best input of each bin size as determined above, H3K27ac with CTCF at 25 kb with a window of 256, CTCF with H3K4me2 at 10 kb and RAD21 with H3K4me3 and SMC3 at 5 kb, both with a window of 512, is trained on the odd chromosomes of GM12878 and its prediction of the held-out chromosomes is scored against the experimentally measured matrix of Rao et al. Rao et al. (2014).

With HiCRep the held-out SCC, averaged over the twelve held-out chromosomes, is 0.586 at 25 kb, between 0.486 on chromosome 20 and 0.751 on chromosome 14; at 10 kb it is 0.623, between 0.539 on chromosome 20 and 0.751 on chromosome 14; at 5 kb it is 0.643, between 0.538 on chromosome X and 0.750 on chromosome 14. GenomeDISCO gives 0.735 at 25 kb, 0.709 at 10 kb and 0.665 at 5 kb, and HiC-Spector 0.568, 0.598 and 0.478, lowest on chromosome 16 at every bin size. The insulation profile correlates at 0.749 at 25 kb, between 0.620 on chromosome X and 0.902 on chromosome 14, at 0.785 at 10 kb and at 0.750 at 5 kb. Chromosome 14 scores highest under nearly every measure, chromosomes 16, 20 and X lowest. On the training chromosomes the same models reach an average SCC of 0.683 at 25 kb, between 0.547 on chromosome 21 and 0.881 on chromosome 13, of 0.770 at 10 kb, between 0.508 on chromosome 17 and 0.933 on chromosome 15, and of 0.773 at 5 kb, between 0.670 on chromosome 11 and 0.912 on chromosome 7. The gap between training and held-out chromosomes is memorisation. The other measures show the same direction: on the training chromosomes HiC-Spector averages 0.613, 0.734 and 0.554, GenomeDISCO 0.758, 0.779 and 0.700, and the insulation correlation 0.896, 0.923 and 0.861 (Figure 6).

**Figure 6:**
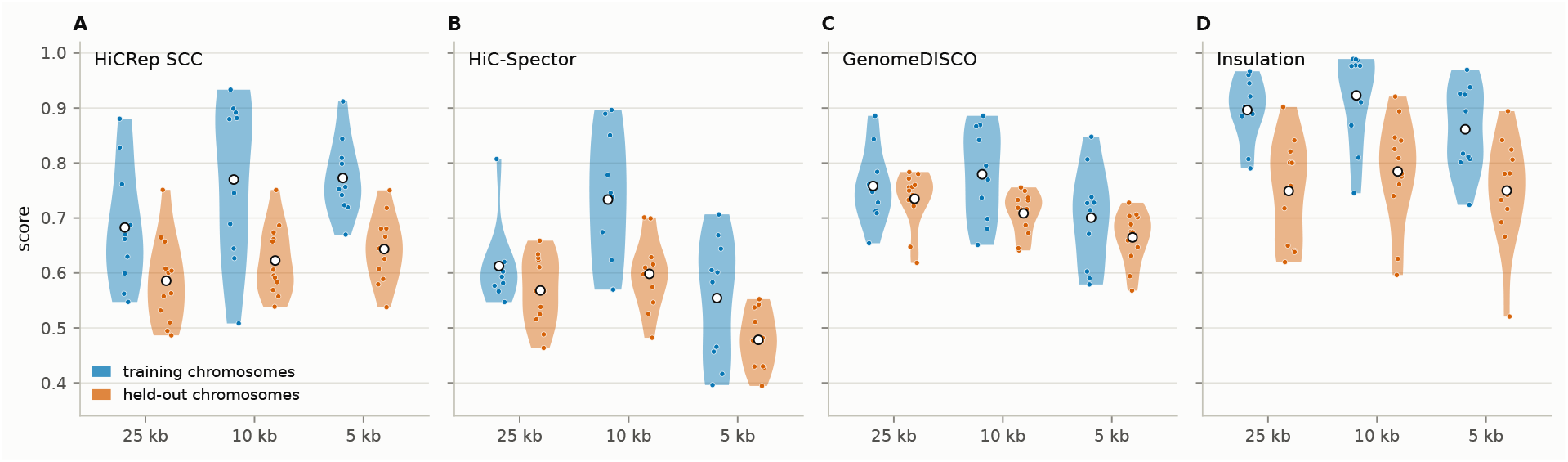
Training against held-out accuracy of the best configurations. HiCRep SCC, HiC-Spector, GenomeDISCO and the insulation correlation for H3K27ac with CTCF at 25 kb, CTCF with H3K4me2 at 10 kb and RAD21 with H3K4me3 and SMC3 at 5 kb, one violin over the ten training and one over the twelve held-out chromosomes, each point one chromosome, the mean marked in white.

### 3.5 Computational cost

Training ran on Nvidia A100 cards with 40 GB of VRAM. On a single card an epoch takes 1 minute at 25 kb with a window size of 64, 40 minutes at 10 kb with a window size of 512 and 90 minutes at 5 kb with the same window; the full set of configurations is costed in Table 1. The 2 kb observed/expected model was trained on eight cards on one node, at 31 minutes per epoch and 51.6 hours for the 100 epochs read here.

**Table 1:** Training cost of the configurations used in this study. Per-epoch times are median values recovered from checkpoint timestamps and therefore include validation and checkpointing. All single-GPU training runs used one Nvidia A100; the 2 kb configuration used eight Nvidia A100 cards with 40 GB of VRAM each on one node. Prediction was run on different hardware and is costed separately in Table 2.

| Bin size | Window | GPUs | Min./epoch | 100 epochs |
| --- | --- | --- | --- | --- |
| 25 kb | 64 | 1 | 1.0 | 1.7 h |
| 25 kb | 128 | 1 | 1.8 | 3.0 h |
| 25 kb | 256 | 1 | 4.2 | 7.0 h |
| 10 kb | 128 | 1 | 4.1 | 6.8 h |
| 10 kb | 512 | 1 | 40.5 | 67.5 h |
| 5 kb | 512 | 1 | 89.5 | 149.1 h |
| 2 kb | 512 | 8 | 31.0 | 51.6 h |

The same times are why the search of the section *Chromatin factor selection* paired combinations with one window per bin size rather than three, and why no stabilisation schemes were explored for the 5 kb configuration.

Predicting a genome needs one consumer card and runs in three steps. *Prepare* reads the bigwigs, bins them and writes the input records. *Predict* runs the generator forward pass over the sliding windows. *Fold* averages the overlapping windows into chromosome matrices and writes the cool file. Only the second uses the GPU, so the three can be queued as separate jobs, each costed for chromosomes 1 to 22 and X on a single Nvidia RTX 4090 (Table 2). Peak device memory is set by the largest chromosome rather than by the genome, because the overlapping windows are summed on the card as they are produced, so every resolution reported here fits on a card with 8 GB.

**Table 2:** Whole-genome prediction cost on a single Nvidia RTX 4090, for chromosomes 1 to 22 and X, using the input tracks each model was trained on. RAM and VRAM are separate pools: RAM is the peak resident set over the whole process tree and peaks in the folding step, VRAM is the peak on the graphics card and peaks in the forward pass.

| Bin size | Window | Wall clock (min) |  |  |  | Peak memory (GB) |  |
| --- | --- | --- | --- | --- | --- | --- | --- |
|  |  | Prepare | Predict | Fold | Total | RAM | VRAM |
| 25 kb | 64 | 0.2 | 1.1 | 0.1 | 1.4 | 1.9 | 2.5 |
| 10 kb | 512 | 0.9 | 11.7 | 1.3 | 13.9 | 2.9 | 4.5 |
| 5 kb | 512 | 1.9 | 23.5 | 2.8 | 28.1 | 3.7 | 4.5 |
| 2 kb | 512 | 1.7 | 37.2 | 6.6 | 45.5 | 7.8 | 6.5 |

### 3.6 Comparison with other approaches

Four published predictors are compared against here, and they differ in what they read. HiC-Reg Zhang et al. (2019) reads fourteen one-dimensional tracks, structural proteins, DNase I and histone marks, together with the genomic distance between the two bins as an explicit feature. Epiphany Yang et al. (2023) reads five: DNase I, CTCF, H3K27ac, H3K27me3 and H3K4me3. C.Origami Tan et al. (2023) reads DNA sequence together with CTCF ChIP-seq and ATAC-seq. Akita Fudenberg et al. (2020) reads DNA sequence alone; its input is therefore the same in every cell type, so it separates them only through the five output heads it was trained with and cannot be directed at a new cell type by supplying that cell type’s data.

Every competing method is used with the models as published by its authors, on the cell type it was published for; the differences between the methods are collected in Table 3.

**Table 3:** Differences between the compared methods. Every property in this table is confounded with accuracy in the comparisons that follow. Akita is multi-task over HFF, H1hESC, GM12878, IMR90 and HCT116, and its GM12878 head is used throughout. Hi-cGAN is evaluated in hg19 except for the comparison against Akita, which is carried out in hg38. Its bin size is configurable and the distance it reaches follows the window size rather than the bin size, and its held-out unit is the chromosome except against Akita, whose split is by window. Akita’s 917,504 bp span is cropped from an input window of 1,048,576 bp.

|  | Hi-cGAN | HiC-Reg | Epiphany | C.Origami | Akita |
| --- | --- | --- | --- | --- | --- |
| Input modality | chromatin tracks | chromatin tracks | chromatin tracks | sequence + tracks | sequence |
| DNA sequence | no | no | no | yes | yes |
| Cell-type-specific input | yes | yes | yes | partly | no |
| Input tracks | 1-2 | 14 + distance | 5 | 2 | none |
| Cell type of weights | GM12878, IMR-90 | GM12878 | GM12878 | IMR-90 | five, jointly |
| Genome build | hg19 | hg19 | hg38 | hg38 | hg38 |
| Native bin size | 2-25 kb, configurable | 25 kb | 10 kb | 8,192 bp | 2,048 bp |
| Prediction span | whole chromosome | 1 Mb | 990 kb | 2 Mb | 917,504 bp, cropped |
| Weights | trained here | released | released | released | released |
| Held-out unit | chromosome, or window | undocumented | chromosome | chromosome | window |
| Output | cool file | text | cool file | array | array |

The Hi-cGAN configuration in each block below was chosen on the held-out chromosomes (section *Chromatin factor selection*), so the two GM12878 blocks were also rescored with the configuration validation chromosome 19 would have chosen. HiCRep falls by 0.031 at 25 kb and 0.021 at 10 kb while the other four measures are unchanged or higher, and every comparison with HiC-Reg keeps its direction; at 10 kb the fall matters, 0.640 against Epiphany’s 0.663 where the held-out-selected 0.662 was level, and 0.640 is the figure we quote (Supplementary Table S17). The IMR-90 block was not rescored, its validation-selected configuration would have to be trained on IMR-90; it is reported under the held-out rule and is optimistic relative to the rescored blocks.

#### 3.6.1 Comparison to Epiphany

Epiphany Yang et al. (2023) is the method closest to Hi-cGAN: it predicts cell-type-specific contact maps from five functional tracks with convolutional and bidirectional LSTM layers. Its released model is for GM12878 on hg38 at 10 kb and emits HiC-DC+ z-scores over the first 990 kb of genomic distance, 100 diagonals. For the match, Hi-cGAN reads CTCF with H3K4me2 at 10 kb, trained on the odd chromosomes of GM12878 in hg19; both outputs are converted to contact counts before scoring, Epiphany’s exponentiated signal through the expected profile of the measured map, and each method is scored against the measured matrix of the build it predicts in.

Epiphany is ahead on HiCRep, 0.663 against the validation-selected 0.640, on the insulation profile, 0.874 against 0.761, and on boundary agreement, 0.106 against 0.076; Hi-cGAN is ahead on HiC-Spector, 0.610 against 0.475, and on GenomeDISCO, 0.716 against 0.499 (Supplementary Table S3). On the distance-stratified correlation only chromosome 2 is available, a training chromosome of Epiphany, where the two remain close, 0.553 against 0.536 Spearman; the two predictions are on different genome builds, so we neither test the difference nor claim parity (Supplementary Table S4). Contact maps of both methods are shown in Figure 8.

#### 3.6.2 Comparison to HiC-Reg

HiC-Reg Zhang et al. (2019) predicts contact counts directly with a random forest, reading fourteen chromatin tracks together with the genomic distance of the bin pair, and reaches at most 1 Mb. The authors’ released predictions for GM12878 are used, in hg19 at 25 kb, the same build and bin size as Hi-cGAN, whose model reads H3K4me1 at a window of 64; HiC-Reg’s log-scaled values are returned to counts and both methods are scored against the measured matrix. The chromosome split behind the released predictions is undocumented, so its values may include training chromosomes and be inflated by the memorisation gap reported above (Supplementary Figure S20).

Hi-cGAN is ahead on HiCRep, 0.746 against 0.671, and on GenomeDISCO, 0.732 against 0.437, level on the insulation profile, 0.851 against 0.853, and behind on HiC-Spector, 0.623 against 0.730, and on boundary agreement, 0.055 against 0.105 (Supplementary Table S3). On the distance-stratified correlation HiC-Reg is ahead by 0.209 Spearman, 0.818 against 0.609, on all eleven held-out chromosomes (Supplementary Tables S4 and S5). Contact maps of both methods are shown in Figure 8.

#### 3.6.3 Comparison to C.Origami

C.Origami Tan et al. (2023) predicts a fixed 2 Mb window from DNA sequence together with CTCF ChIP-seq and chromatin accessibility, at a native bin size of 8,192 bp, and was built for in silico screening of sequence perturbations. Its released weights and both feature tracks are for IMR-90 in hg38, so the comparison is made on IMR-90. C.Origami’s log-scaled output is returned to counts and regridded onto the 8 kb grid of the measured map, which is left untouched, and it is scored on chromosome 2, the chromosome its authors held out. Hi-cGAN predicts IMR-90 at 10 kb in hg19 with CTCF and H3K4me2, using the IMR-90 model of the section *Prediction across cell types*, which is trained on all chromosomes except 19, so chromosome 2 lies in its training data. The two are scored at different bin sizes, each against the measured map of its own build; regridding the C.Origami pair onto the 10 kb grid instead moves its scores by at most 0.033.

C.Origami reaches a HiCRep score of 0.914 against Hi-cGAN’s 0.724, is ahead on HiC-Spector, 0.663 against 0.552, on GenomeDISCO, 0.724 against 0.695, and on boundary agreement, 0.091 against 0.070, and level on the insulation profile, 0.872 against 0.882 (Supplementary Table S3). On the distance-stratified correlation it reaches 0.776 against 0.482 Spearman (Supplementary Table S4). Hi-cGAN’s chromosome 2 is a training-chromosome value, so its side of this block is, if anything, overstated: the gap to C.Origami is not explained by the split. Contact maps of both methods are shown in Figure 8.

#### 3.6.4 Comparison to Akita

The last comparison is against a predictor that reads no chromatin data at all. Akita Fudenberg et al. (2020) is a convolutional network that takes about one megabase of DNA sequence and returns the normalized Hi-C map of that region at 2 kb resolution, trained jointly for five cell types, of which the GM12878 output is used here. Hi-cGAN never sees sequence and Akita never sees a chromatin track, so the comparison measures how far protein occupancy alone determines contact structure. For the match, Hi-cGAN reads a single CTCF track and is trained on Akita’s own normalized target. Akita holds out windows rather than chromosomes, so its held-out windows were excluded from Hi-cGAN’s training, and both methods are scored on exactly those windows, on the same pixels and against the same reference (Supplementary Section S6).

Across the 411 held-out windows Akita reaches a mean per-window Pearson correlation of 0.506 against Hi-cGAN’s 0.238 and is ahead on 371 of them, with the same ordering on Spearman, 0.488 against 0.232, and on the insulation profile, 0.492 against 0.297; Hi-cGAN places 22.4% of insulation boundaries within two bins of a measured boundary, Akita 40.2%. GenomeDISCO and HiC-Spector discriminate nothing below one megabase and are reported only to record it (Supplementary Table S5, Supplementary Figure S8). Contact maps of both methods are shown in Figure 8.

Akita’s lead is widest at the scale of topologically associating domains, 0.303 between 100 and 200 kb, and narrows below 50 kb and beyond 717 kb, where both approach zero (Supplementary Figure S9). The difference is one of calibration: regressing predicted contrast on measured contrast over the windows gives a slope of 0.71 for Akita and 0.22 for Hi-cGAN, so Akita produces structure where structure is present, whereas Hi-cGAN produces it regardless; windows across the range of outcomes are shown in Supplementary Figure S10.

#### 3.6.5 Baselines and the behaviour of the measures

Two baselines bound what the scores mean. A ridge regression per genomic distance on Hi-cGAN’s own input reaches 0.691 held-out SCC against 0.746, yet draws stripes rather than domains (Supplementary Figure S17); a map of the distance decay alone reaches 0.003 on HiCRep but 0.663 on HiC-Spector and 0.709 on GenomeDISCO. A high SCC therefore does not certify a usable map, and the latter two measures discriminate nothing at this scale. The remaining rank flips follow from what each correlation family rewards, fine within-diagonal variation against domain-scale agreement (Supplementary Tables S3 to S5).

Called features are the stricter test: boundary *F*_1_ puts Hi-cGAN last, 0.300 to 0.355 against 0.416 for Epiphany, 0.650 for C.Origami and 0.824 for HiC-Reg, loops are recovered poorly by all methods except C.Origami, and rescaling the predicted contrast changes nothing, Hi-cGAN calling 275 boundaries per chromosome against 259 measured: boundaries are misplaced, not missing (Supplementary Section S8 and Table S10).

A Hi-cGAN prediction is an ordinary cool file Abdennur and Mirny (2020): the matrices here are read by HiCExplorer Wolff et al. (2018, 2020) and hicrep without conversion, and Figure 7 draws a measured and a predicted matrix with pyGenomeTracks Lopez-Delisle et al. (2020) from one track definition.

**Figure 7:**
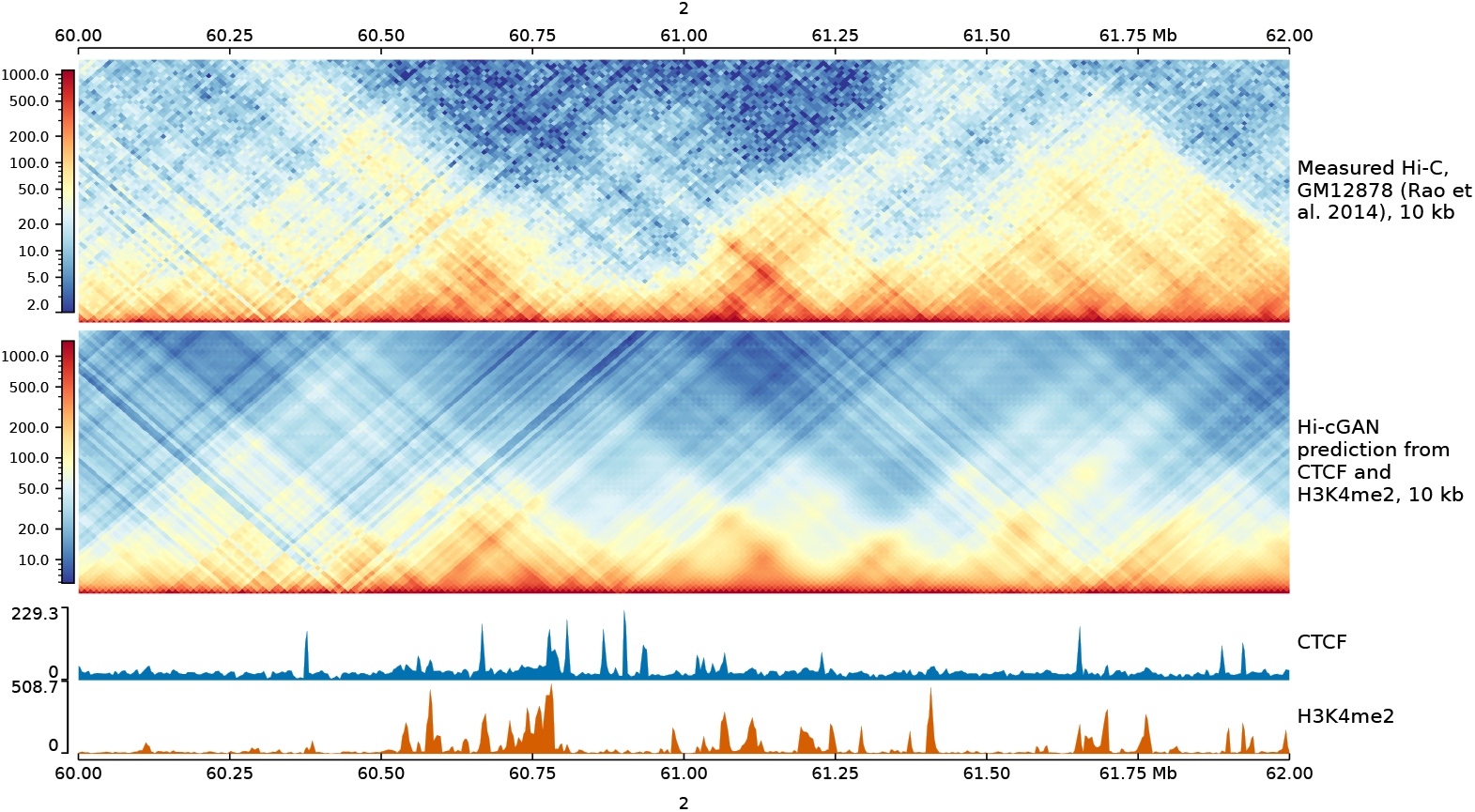
Measured and predicted Hi-C at chr2:60-62 Mb, GM12878, 10 kb. Measured Hi-C (top), the Hi-cGAN prediction from CTCF and H3K4me2 (middle), and the two chromatin tracks the prediction was made from (bottom). Both matrices are drawn by pyGenomeTracks from a single track definition, with the same track type, depth and colour map; only the file name differs between them. The predicted map reproduces the domain boundaries and their positions, but is smoother and lower in contrast than the measurement, which is the behaviour quantified in the section *Comparison to Akita*.

**Figure 8:**
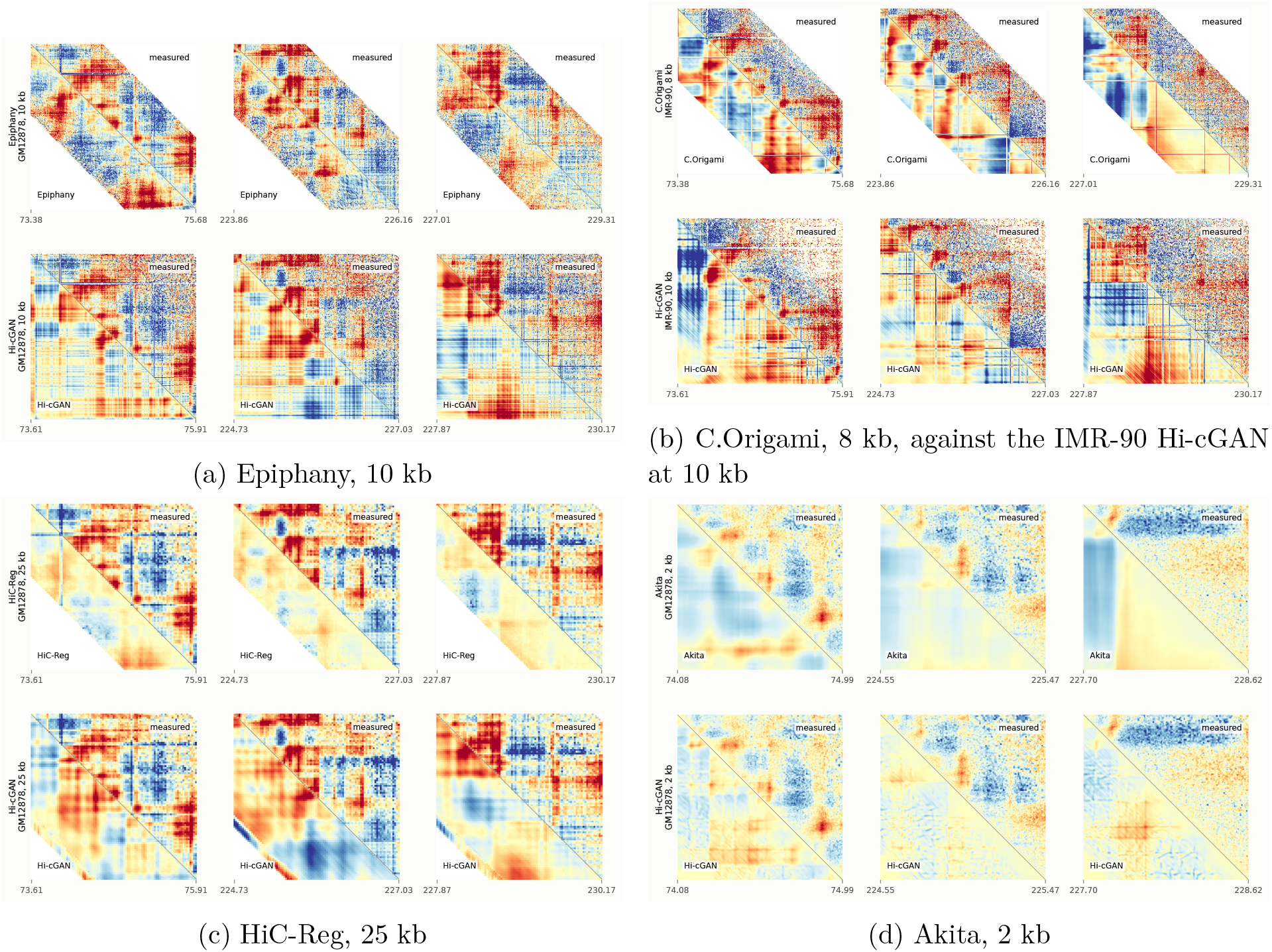
Every comparison on the same three regions. Three regions of chromosome 2, drawn from Akita’s held-out windows because Akita predicts nowhere else, coordinates lifted between genome builds so every panel is centred on the same locus. Each block shows the competing method on top and Hi-cGAN at the same bin size below, the measured map in the upper right of every panel and the prediction in the lower left, all as log2 observed/expected on one colour scale. Panels span 2.3 Mb, the Akita block its fixed 0.92 Mb window. Empty corners mark what a panel cannot contain: Epiphany reaches 990 kb, and the HiC-Reg and C.Origami predictions are banded to the 1 Mb used for scoring. The same regions in the ordinary contact-count view are Supplementary Figure S11.

### 3.7 Prediction across cell types

The network reads chromatin tracks rather than sequence, so a trained model is applied to a different cell type by exchanging its input tracks, with no retraining. Every pairing of five cell types was scored this way at 25 kb, each model predicting each target from that target’s own tracks (Figure 9). Predicting the cell type a model was trained on gives 0.670 to 0.750; predicting a different one gives 0.493 to 0.649. Transfer therefore costs roughly 0.1 to 0.2 SCC, and the loss is asymmetric: HUVEC is the hardest target from every source, between 0.493 and 0.527, while it is no worse than the others as a source. The same matrix at 10 kb has the same shape with a slightly higher diagonal, 0.715 to 0.765.

**Figure 9:**
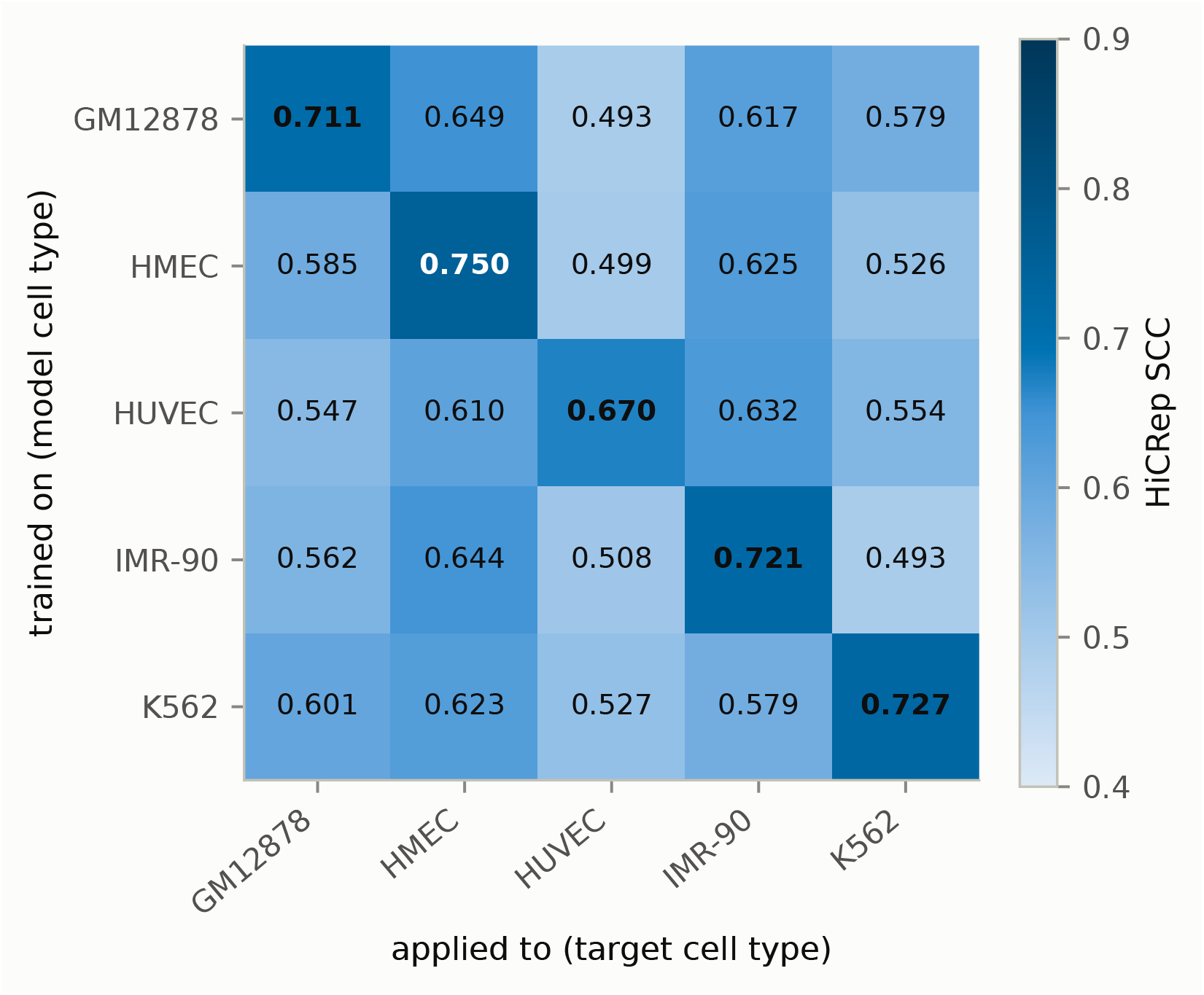
Every pairing of five cell types at 25 kb. A model trained on the cell type of the row, applied to the cell type of the column using that target’s own chromatin tracks. The diagonal is the cell type the model was trained on and is bold. HiCRep SCC, mean over chromosomes.

Training on four cell types and predicting the fifth, which contributed no training data at all, gives 0.545 for HUVEC, 0.578 for K562, 0.590 for GM12878, 0.625 for IMR-90 and 0.659 for HMEC, a mean of 0.600 over 23 chromosomes each (Supplementary Figure S18). That is above the 0.574 obtained by transferring from a single source cell type, so training across cell types helps, and below the 0.716 the same models reach on the cell type they were trained on. The gap between those two figures is what a user should expect to pay for a cell type with no Hi-C of its own.

## 4 Discussion

In this study, Hi-cGAN, which predicts Hi-C matrices from chromatin factor occupancy alone, was evaluated across resolutions, input factors and cell types and compared against four published predictors. From one or two tracks it reaches a held-out HiCRep SCC of 0.586 to 0.746 depending on the resolution, is close to Epiphany at 10 kb, 0.640 against 0.663, ahead of HiC-Reg on HiCRep and GenomeDISCO at 25 kb, and clearly behind the sequence-reading C.Origami and Akita on every measure. Which factor carries the signal follows the biology of each scale: CTCF and cohesin, which create loop anchors and domain boundaries through loop extrusion, lead at 5 and 10 kb Rao et al. (2014, 2017), the marks of active chromatin lead at 25 kb, where compartmentalisation is what remains resolvable Lieberman-Aiden et al. (2009); Schwarzer et al. (2017); Henikoff and Smith (2015). Predictive value is not evidence of a causal role, and transfer to a cell type never seen in training costs about 0.12 SCC, 0.600 against 0.716.

The ranking of the methods depends on the measure used. Distance-stratified correlation rewards fine variation within a diagonal and favours HiC-Reg and the sequence methods; HiCRep rewards agreement at domain scale and favours Hi-cGAN’s sharp-boundary maps. A linear baseline reaches 0.691 SCC while drawing stripes no analysis would accept, a map of the distance decay alone beats several predictions on HiC-Spector and GenomeDISCO, and boundary agreement contradicts the insulation correlation for every method. A single accuracy number for a Hi-C predictor is therefore not well defined, although papers in this area commonly report one.

The overriding observation is that no method is precise. Akita, the best per window, has a median per-window correlation of 0.551, with 18% of its held-out windows below 0.3; Hi-cGAN’s median is 0.238 with 60% below. Boundary *F*_1_ spans 0.300 to 0.824 across the methods and loops are recovered poorly by all but C.Origami. Above all, no method reaches the agreement of biological replicates: the reproducibility measures were calibrated to separate replicate experiments from non-replicates Yardımcı et al. (2019), and against the threshold of 0.8 its authors propose for GenomeDISCO Ursu et al. (2018), every prediction in this study, the sequence-based ones included, is classified as a non-replicate. The single value approaching replicate range is C.Origami’s HiCRep of 0.914, on one chromosome and one measure.

Part of this imprecision sits in the models. No pixel loss rewards abstention: regressing predicted on measured contrast gives slopes of 0.22 to 0.71, so every model, Hi-cGAN most strongly, emits structure whether or not structure is present, and rescaling the output afterwards repairs nothing. The adversarial term itself does no measurable work in the one configuration ablated, the three losses differing by less than the seed-to-seed spread, so Hi-cGAN behaves there as a convolutional encoder-decoder; that null rests on three seeds and one chromosome per role (Supplementary Section S16). Part may sit in the input: population-averaged occupancy of a dozen proteins may underdetermine folding, since chromatin tracing finds domains in single cells at positions that average away in the population map Bintu et al. (2018). And part sits in the target: the windows every method scores worst on are the least sampled, contact count correlating with measured contrast at 0.482 and with Akita’s accuracy at 0.556, and a 918 kb window at 2 kb holds about four reads per pixel even in a map of 2.3 billion contacts. A featureless target is indistinguishable from thin sampling, protocol biases are fixed features of a locus missing from every replicate, and assays of the same regions at greater depth reveal loops that Hi-C does not show Krietenstein et al. (2020); Liu et al. (2024). This is no formal upper bound, but part of every accuracy reported in this field is set by what the assay did not capture.

Deciding between these explanations needs different data, not different scores. Rescoring the low-coverage windows against an assay with another capture bias, Micro-C or capture Hi-C, would separate unstructured loci from unmeasured ones, and translation between the assays is feasible Liu et al. (2024). Sequence could be added as an input rather than treated as a competitor, as C.Origami does, and training on perturbation data such as cohesin depletion Rao et al. (2017) would test whether a model has learned the mechanism or only its correlates.

## 5 Availability of Source Code and Requirements

- Project name: Hi-cGAN
- Project home page: https://github.com/joachimwolff/Hi-cGAN
- Operating system: Linux
- Programming language: Python 3
- Other requirements: TensorFlow 2, cooler, pyBigWig, scikit-learn; a CUDA-capable GPU is needed
- License: GPLv3
- RRID: SCR_026732

Exact package versions, and the contents of the archive, are given in Supplementary Section S21.

## 6 Availability of Supporting Data and Materials

The hg19 Hi-C data is from Rao et al. Rao et al. (2014), GEO accession GSE63525, in the quality-filtered combined_30 form, coarsened to 5, 10 and 25 kb: GM12878 for training and for the accuracies of Supplementary Tables S4 and S5, K562 for the cross-cell-type prediction of Supplementary Section S18, and IMR-90 for the hg19 side of the C.Origami comparison. The chromatin protein and DNase-seq tracks, with their ENCODE and GEO identifiers, are listed in Supplementary Table 1.

The comparisons carried out in hg38 rest on two measured matrices. GM12878 is the matrix of the 4D Nucleome Data Portal Reiff et al. (2022), accession 4DNFIXP4QG5B, read at 10 kb where Epiphany is scored and at 2 kb, KR balanced and observed over expected, as the source of the target for the Akita comparison, to which Akita’s own transform is applied. IMR-90, against which C.Origami is scored, is a measured 8 kb matrix; it is aggregated to 10 kb together with C.Origami’s 8,192 bp output as described in the section *Comparison with other approaches*. A second hg38 GM12878 matrix was tested and rejected as a reference matrix. We fixed the criterion before scoring either candidate: two measured Hi-C maps of different cell types agree at a distance-stratified Spearman correlation of roughly 0.5 to 0.8, so we required at least 0.45 against the measured IMR-90 map. The 4D Nucleome matrix used here reaches 0.571 and the rejected one 0.328. The check is part of the released code and both scores are archived with the data.

Hi-C data was processed with cooler Abdennur and Mirny (2020) and HiCExplorer Wolff et al. (2018, 2020), and plotted with pyGenomeTracks Lopez-Delisle et al. (2020). The comparison against a sequence-based predictor uses the weights, parameter file and test records published with Akita Fudenberg et al. (2020); those records supply both input sequence and target, so the comparison depends on no data preparation of ours beyond the interval file defining the held-out regions.

All processed data, the held-out interval file, the per-window scores for every method under every measure, the per-window implementations of SCC, insulation agreement, GenomeDISCO and HiC-Spector, the script validating SCC against the hicrep reference implementation, and every script producing a figure or table, including those written with the assistance of the tools declared under *Use of Artificial Intelligence Tools*, are on Zenodo at 10.5281/zenodo.11402892.

## 9 Authors’ Contributions

R.K. developed and implemented the presented approach as part of his master thesis at the Bioinformatics Lab, University Freiburg, Germany, in the winter semester 2019/2020. The master thesis is available on the Bioinformatics Lab website: http://www.bioinf.uni-freiburg.de/Lehre/Theses/MA_Ralf_Krauth.pdf. A.K. and J.W. supervised the master thesis. J.W. computed all evaluations and wrote the publication.

## 10 Use of Artificial Intelligence Tools

Generative artificial intelligence was used in the preparation of this work and is declared here in full. The Hi-cGAN source code was written by R.K. Claude Code Anthropic (2026) was subsequently used for computational analysis and code improvement: writing and refactoring the analysis and plotting scripts, implementing the per-window reproducibility measures and their validation against the reference implementations, running the evaluations in the Results section, and improving the text of this manuscript. Claude Opus 4.8 was used during development of the analysis code, Claude Opus 5 for the comparison against Akita, the per-window measures and the associated figures, and Claude Fable 5 for the restructuring and editing of the manuscript, the region and violin figures, and the evaluation of the selected configurations across all measures.

ChatGPT-4 OpenAI (2024) was used to check spelling and grammar in an earlier draft, parts of whose Background and architecture description survive in the present text.

## 11 Acknowledgments

We thank de.NBI-Cloud and the UFR-RZ for GPU resources for R.K., A.K. and J.W. The authors acknowledge the Hannover Medical School for providing MHH-HPC resources and technical support that have contributed to the research results reported within this paper.

## Supplementary material

### S1 Data collection and pre-processing

Every chromatin track used is listed with its accession in Table S1.

**Table S1:** Chromatin features used for the learning and prediction of Hi-C matrices using Hi-cGAN.

| feature name | cell line GM12878 | cell line K562 |
| --- | --- | --- |
| CTCF | GSM733752 | GSM733719 |
| DNaseI | wgEncodeEH000534 | wgEncodeEH000530 |
| H3k27ac | GSM733771 | GSM733656 |
| H3k27me3 | GSM733758 | GSM733658 |
| H3k36me3 | GSM733679 | GSM733714 |
| H3k4me1 | GSM733772 | GSM733692 |
| H3k4me2 | GSM733769 | GSM733651 |
| H3k4me3 | GSM733708 | GSM733680 |
| H3k79me2 | GSM733736 | GSM733653 |
| H3k9ac | GSM733677 | GSM733778 |
| H3k9me3 | GSM733664 | GSM733776 |
| H4k20me1 | GSM733642 | GSM733675 |
| Rad21 | wgEncodeEH000749 | wgEncodeEH000649 |
| Smc3 | wgEncodeEH001833 | wgEncodeEH001845 |

### S2 Data preparation

Hi-C data produced by Rao et al. (2014) was obtained in hic file format from the Gene Expression Omnibus with the accession key GSE63525. We selected the quality-filtered *combined*_30 matrices, which include high-quality reads from both replicates. These matrices were then processed to 5 kbp bin size, converted to cooler format using hic2cool, and further coarsened to resolutions of 10 and 25 kbp with cooler coarsen, with the commands given in the section *Commands*. Unlike Farré and Emberly (2018), who employed ICE-plus distance-based normalization, and others who use ICE-or KR-normalization, these matrices were left unnormalized, no advantage having been observed from normalizing them.

ChIP-seq data covering 13 chromatin features plus DNaseI-seq data were used, as listed in Table S1. Read alignments for the first and second replicates were downloaded in bam format from either the ENCODE project or directly from the University of California’s (UCSC) file server in their latest versions. Identifiers from UCSC (wgEncode…) and GEO (GSM…) are listed in Table S1. The bam files were then converted to bigwig format for easier handling, and the replicates were combined into a single bigwig file by averaging.

The selection of chromatin features aligns closely with the approach of Zhang et al. (2019), incorporating structural proteins like CTCF and Cohesin subcomponents RAD21 and SMC3, along with markers of active/passive chromatin states.

Hi-C data was processed with cooler (Abdennur and Mirny (2020) and HiCExplorer (Wolff et al. (2018, 2020), Hi-C plots were generated with pyGenomeTracks Lopez-Delisle et al. (2020).

### S3 Neural network description

A prediction is assembled from overlapping sliding windows (Figure S1). The three networks are described below.

#### Embedding network

The chromatin input for one sample is an array of 3*w* bins by *n_f_* factors, where *w* is the window in bins and the factor of three is the window with its two flanks. Eight Conv1D blocks of kernel 4, stride 1 and same padding, each followed by group normalization with one group per channel and a LeakyReLU with *α* = 0.2, take the channel count from *n_f_*through 1024, 512, 512, 256, 256, 128, 128 to 64, holding the length at 3*w*. A ninth Conv1D then maps the one-dimensional track to a matrix: kernel 4, *stride 3*, same padding, a sigmoid activation and *w* output filters, which divides the length by three and sets the channel count to the window, giving a *w ×w* tensor. It is averaged with its transpose to enforce symmetry, and the final reshape only appends the channel axis to give (*w, w,* 1).

The window therefore enters the architecture twice, in the input length 3*w* and in the filter count of that ninth convolution, and the output is *w × w* for every *w* rather than for one distinguished value. The filter count is the only part of the embedding that changes with the window, which is why the parameter counts in Table S2 barely move between 64 and 512. Every convolution carries an *L*_2_ kernel penalty of 0.01. The generator and the discriminator each use a separate embedding network of this structure; the embedding weights are not shared.

#### Generator

The generator is a U-Net. The encoder is eight Conv2D blocks of kernel 4 and stride 2 with 64, 128, 256, 512, 512, 512, 512 and 512 filters, group normalization on all but the first and last, and LeakyReLU. The decoder is seven Conv2DTranspose blocks of kernel 4 and stride 2 with 512, 512, 512, 512, 256, 128 and 64 filters, group normalization and ReLU, each concatenated with the encoder block of matching resolution. Dropout at rate 0.5 is applied in the first three decoder blocks. Windows below 256 bins drop encoder and decoder blocks in pairs, which leaves five decoder blocks and two with dropout at a window of 64. A final Conv2DTranspose of kernel 4 and stride 2 gives one channel, which is averaged with its transpose to enforce symmetry and passed through a sigmoid.

#### Discriminator

A conditional PatchGAN. The embedded chromatin input and the matrix to be classified are concatenated into two channels, then passed through Conv2D blocks of kernel 4 and stride 2, one Conv2D of kernel 4 and stride 1 with 512 filters, and a final Conv2D of kernel 4 and stride 1 to a single channel. Windows above 64 use three strided blocks with 64, 128 and 256 filters; at a window of 64 the first two are replaced by a single block of 256, leaving two strided blocks and halving the depth of the downsampling. Every intermediate tensor is averaged with its transpose, so symmetry is imposed throughout the discriminator and not only on the generator output. The output is a grid of logits, one per patch, and the sigmoid of the binary cross-entropy *L_d_*in the main text is applied to them.

**Table S2:** Sizes of the three networks. Parameter counts are for a single input factor; each further factor adds to the first embedding convolution only. The PatchGAN output is the grid of patch verdicts, and the receptive field is the extent in bins of the input that one output unit sees, accumulated backwards through the convolutions by *r →* (*r −* 1)*s* + *k* from a single output pixel, with *k* the kernel and *s* the stride. It is 34 bins at a window of 64, where the discriminator has two strided blocks, and 70 at 256 and above, where it has three. One output unit therefore covers a little over half the window at 64, a quarter of it at 256 and about an eighth at 512: the discriminator is a local critic at every window, least so at the smallest.

| Window | Generator<br>params | Discriminator<br>params | Embedding<br>params | Embedding<br>output | PatchGAN<br>output | Receptive<br>field |
| --- | --- | --- | --- | --- | --- | --- |
| 64 | 33,428,609 | 7,354,369 | 4,190,720 | 64x64x1 | 16x16x1 | 34 |
| 256 | 58,647,873 | 7,004,097 | 4,240,064 | 256x256x1 | 32x32x1 | 70 |
| 512 | 58,713,665 | 7,069,889 | 4,305,856 | 512x512x1 | 64x64x1 | 70 |

#### Assembling a chromosome from overlapping windows

The generator emits one *w × w* upper triangle per sliding-window position, and a chromosome is the average of the triangles that cover each cell. The procedure is the following.

1. Windows slide by one bin. Position *j* reads chromatin bins [*j, j* + 3*w*), the window with a flank of *w* on each side, and predicts the triangle covering matrix bins [*j* + *w, j* + 2*w*).
2. Each predicted triangle is scattered into a band accumulator of shape (*w, N*) indexed by distance from the diagonal and by row, *N* being the number of bins on the chromosome. A window of *w* bins cannot place a value further than *w −* 1 off the diagonal, so the band holds the whole result and the matrix is never formed densely.
3. Every covering window contributes with equal weight. The divisor for a cell on diagonal *d* at row *p* is the number of window starts in [*p −* (*w −* 1 *− d*)*, p*], obtained from a prefix sum over the start positions rather than from a stored count matrix. Cells covered by no window, which occur only within *w* bins of a chromosome end or where prediction was restricted to given regions, stay empty and are not written.
4. The averaged band is converted to a sparse upper-triangular matrix. Symmetry is not imposed here: it is already enforced inside both networks, and the cool file is written in symmetric_upper form, which is how cooler stores a symmetric matrix.
5. Values are returned to contact scale. With a stored target value range the affine map that placed the target in [0, 1] during training is inverted and nothing further is applied. Without one, the matrix is scaled to the unit range and multiplied by --multiplier, which is a display scale for raw-count predictions. Values stay floating point; no rounding to integers takes place.

Prediction draws no randomness: dropout is inactive, the generator takes no noise input, and the assembly is a sum and a division in a fixed order. It is not bitwise reproducible on a GPU. Predicting chromosome 22 twice from the same checkpoint on the same card gives the same 121,184 pixels at the same coordinates, of which 7.3% agree bit for bit and the rest differ by at most 1.7*×*10*^−^*^6^ in relative terms, the two runs correlating to twelve decimal places. The residual is float32 reduction order inside cuDNN, not anything in the model: the same two runs on CPU are bitwise identical. A CPU and a GPU run of the same checkpoint agree to a Pearson correlation of 0.999 999 97, with a median absolute difference of 6.4 *×* 10*^−^*^4^ on values reaching 1,000. Nothing reported in this article is sensitive at that magnitude, and a reader needing bitwise reproduction should predict on CPU.

**Figure S1:**
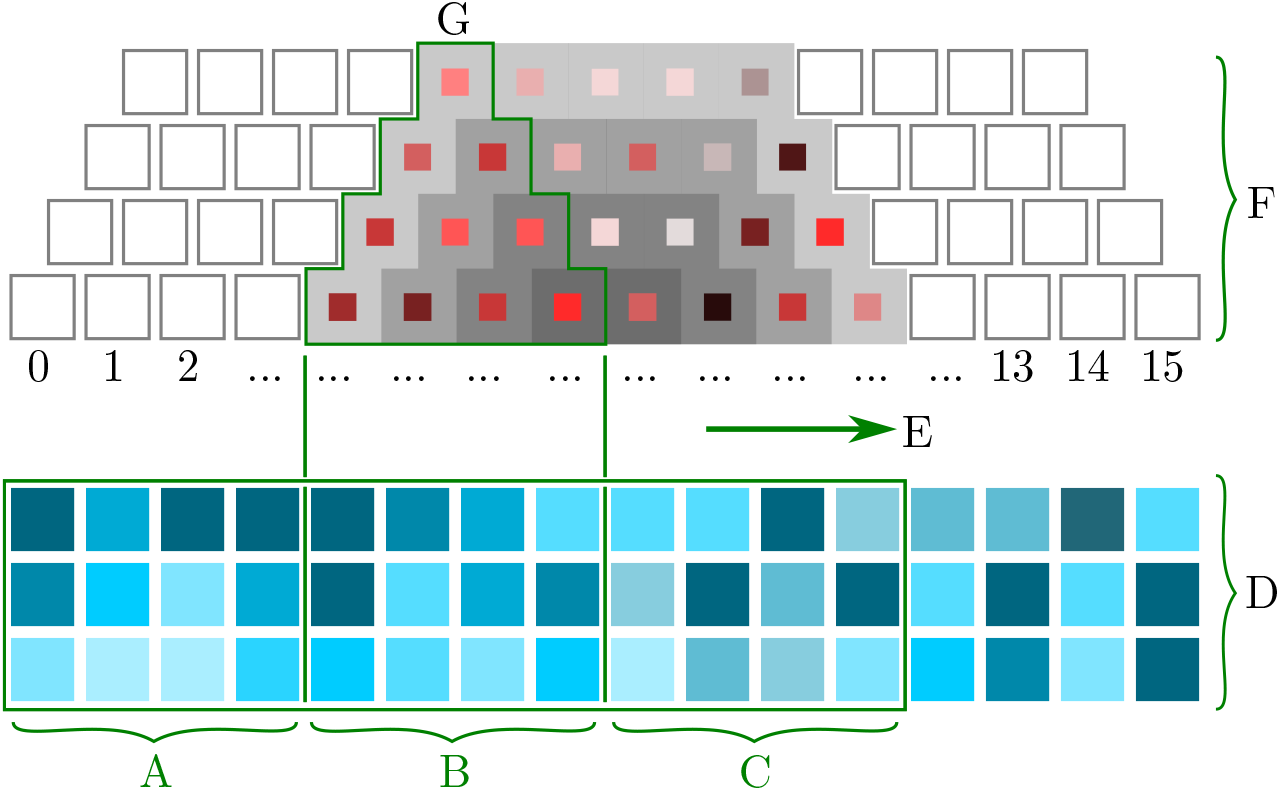
Prediction process

### S4 Training statistics

Generator and discriminator losses over training (Figures S2, S3 and S4) are shown for the resolutions and window sizes used in the main text. They document training behaviour and convergence; they do not compare configurations, and stable losses are not by themselves evidence of an equilibrium.

**Figure S2:**
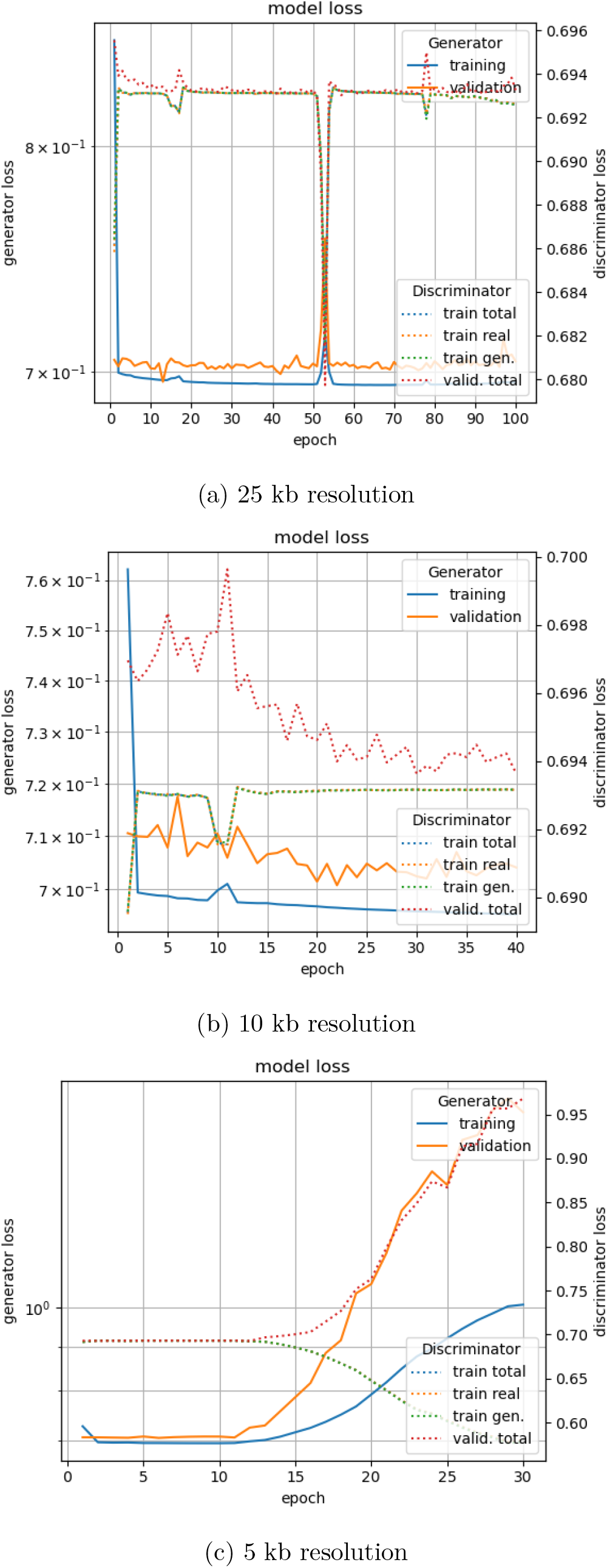
Model loss over training epochs, three resolutions. On GM12878 at 25 kb, 10 kb and 5 kb, all for a window size of 256.

**Figure S3:**
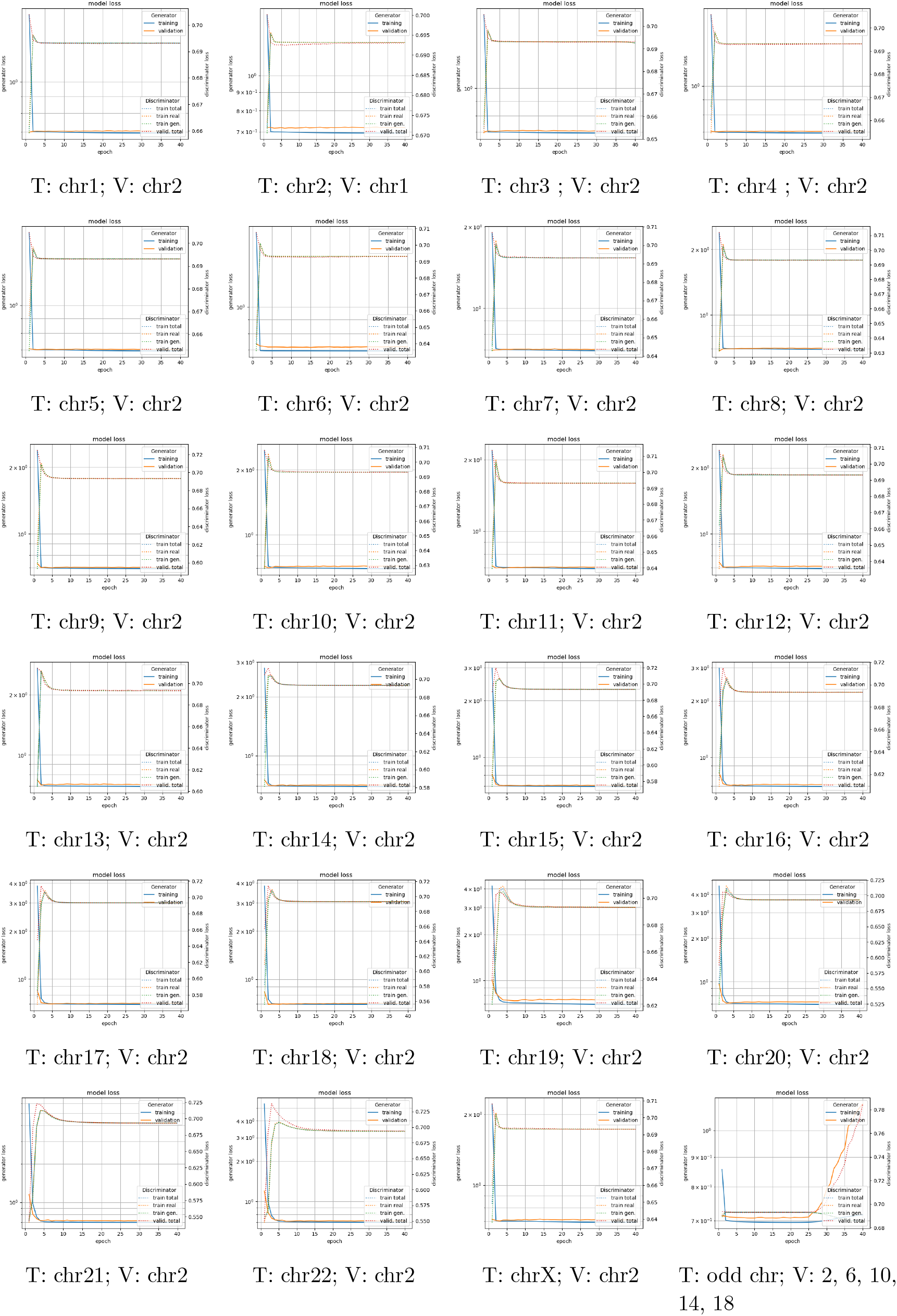
Loss over epochs for 23 individually trained models on a single chromosome. Validation always on chromosome 2, except for training on chromosome 2 with a validation on chromosome 1. Subfigure (f) shows the training on the odd chromosomes 1, 3, 5, 7, 9, 11, 13, 15, 17, 19, 21 and the validation on 2, 6, 10, 14 and 18. Training for a window size of 128.

**Figure S4:**
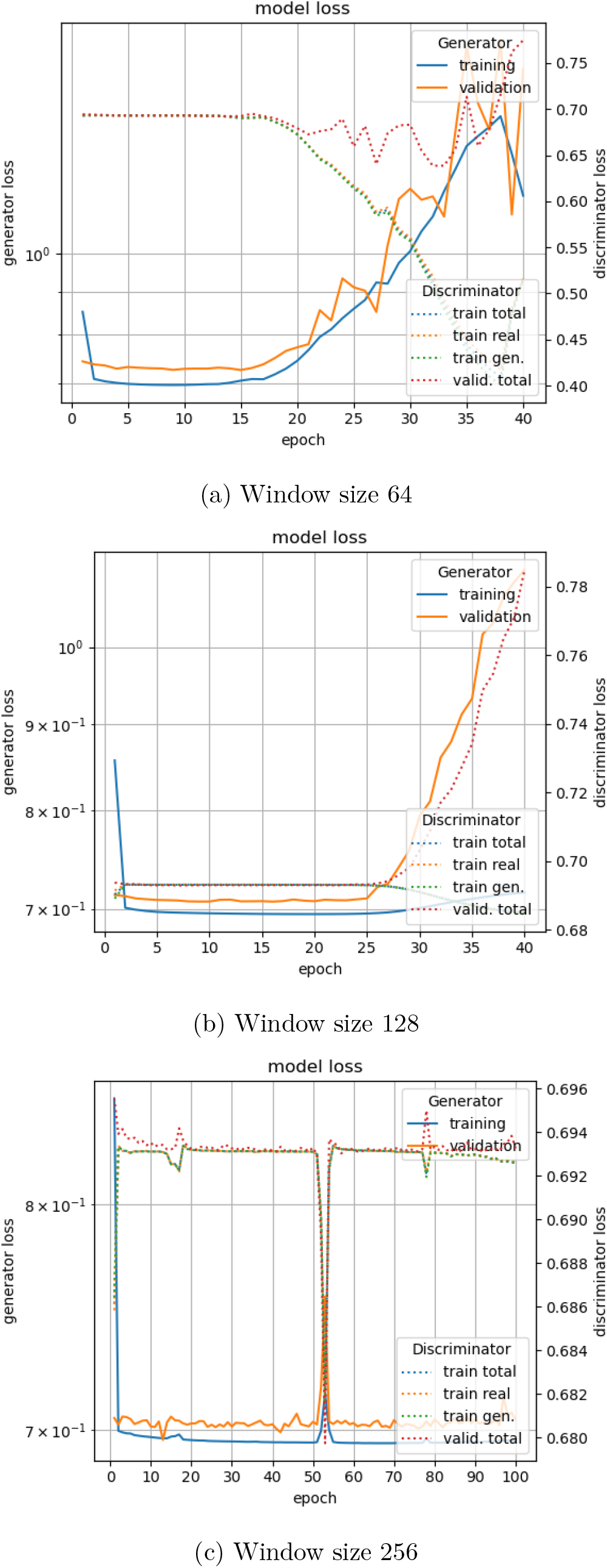
Model loss over training epochs, three window sizes. On GM12878 at 25 kb for window sizes 64, 128 and 256.

### S5 Method comparison tables

The three tables behind the method comparison of the main text.

**Table S3:** Established Hi-C measures across all four methods. Mean over the held-out chromosomes of each block, on count-space matrices banded to 1 Mb: HiCRep SCC Yang et al. (2017), HiC-Spector Yan et al. (2017), GenomeDISCO Ursu et al. (2018), insulation-score Pearson Crane et al. (2015) and boundary Jaccard. Best per block in bold. Blocks are not comparable with one another, and the GM12878 blocks rest on one to twelve chromosomes. The two IMR-90 rows are scored at different bin sizes, each near its own native one, so no best is marked there; each is scored against the measured map of the build it predicts in, and C.Origami on the chromosome its authors held out. The split used for the released HiC-Reg model is not documented. The ridge row is one regression per genomic distance from the same single track Hi-cGAN reads. The distance-only row is not a method but the measured matrix with every pixel replaced by the mean of its diagonal, and its insulation profile is flat, so that correlation is undefined.

| Block | Method | HiCRep | Spector | DISCO | Insul. | Bound. |
| --- | --- | --- | --- | --- | --- | --- |
| 25 kb GM12878 | Hi-cGAN | **0.746** | 0.623 | **0.732** | 0.851 | 0.055 |
|  | HiC-Reg | 0.671 | **0.730** | 0.437 | **0.853** | **0.105** |
|  | ridge | 0.691 | 0.549 | 0.538 | 0.701 | 0.081 |
|  | distance only | 0.003 | 0.663 | 0.709 | – | 0.000 |
| 10 kb GM12878 | Hi-cGAN | 0.662 | **0.610** | **0.716** | 0.761 | 0.076 |
|  | Epiphany | **0.663** | 0.475 | 0.499 | **0.874** | **0.106** |
| IMR-90 | Hi-cGAN, 10 kb | 0.724 | 0.552 | 0.695 | 0.882 | 0.070 |
|  | C.Origami, 8 kb | 0.914 | 0.663 | 0.724 | 0.872 | 0.091 |

## S6 Comparison against a sequence-based predictor

All figures in this section cover the 411 of Akita’s 413 held-out test windows for which both methods produced a prediction; the two omitted windows fall below 50% coverage by Hi-cGAN because they lie within one window length of a chromosome end.

### S6.1 Held-out protocol and verification

Akita separates its data by window rather than by chromosome. Every sliding-window position overlapping an Akita validation or test region was therefore excluded from Hi-cGAN training, and prediction was restricted to windows lying entirely inside those regions, containment being the exact complement of the overlap test used during training. An audit of the resulting matrix reports that no predicted bin falls outside a held-out region and that 227,462 of 230,464 held-out bins were reached, or 98.70%. Of the 3,002 bins not reached, 2,240 lie in regions too short to contain a single 512-bin window and 762 lie within one window of a chromosome start or end. No gap remained that was attributable to neither cause.

### S6.2 Reproduction of the reference Akita evaluation

Akita was run locally from the published weights and parameter file, over the test records distributed with the model, which carry the input sequence and the target together. Akita is multi-task over five cell types, HFF, H1hESC, GM12878, IMR90 and HCT116, and the main text uses the GM12878 head throughout to match the cell type Hi-cGAN predicts. As a check on the local run, the HFF head was also scored and compared against the per-region correlations distributed with the model: the two agree to a maximum absolute difference of 9.8 *×* 10*^−^*^6^. Mean per-window correlations for all five heads were 0.545 (HFF), 0.610 (H1hESC), 0.505 (GM12878), 0.510 (IMR90) and 0.492 (HCT116), so the choice of head changes the reference figure by up to 0.118 and has to be stated.

**Table S4:** Hi-cGAN against every method compared here. Spearman and Pearson on held-out data, stratified by genomic distance for the three chromosome-scale blocks and taken over the whole window for the 2 kb block, where the unit of prediction is the window. Blocks are separated because they differ in resolution, cell type and genome build and are not comparable across rows. In the IMR-90 block each method is scored against the measured IMR-90 map of the build it predicts in, hg38 for C.Origami and hg19 for Hi-cGAN.

| Block | Method | Spearman | Pearson |
| --- | --- | --- | --- |
| 25 kb, GM12878, 11 chr | HiC-Reg | 0.818 | 0.836 |
|  | Hi-cGAN | 0.609 | 0.686 |
| 10 kb, GM12878, chr2 | Epiphany | 0.553 | 0.541 |
|  | Hi-cGAN | 0.536 | 0.426 |
| 10 kb, IMR-90, chr2 | C.Origami | 0.776 | 0.825 |
|  | Hi-cGAN | 0.482 | 0.480 |
| 2 kb, GM12878, 411 windows | Akita | 0.488 | 0.506 |
|  | Hi-cGAN | 0.232 | 0.238 |

**Table S5:** Paired comparisons over the 411 held-out windows. Each comparison is paired on the same window, or on the same chromosome for the last row, and tested with a two-sided Wilcoxon signed-rank test; the interval on the mean difference is a 20,000-sample percentile bootstrap. *Other* is Akita except in the last row, where it is HiC-Reg at 25 kb. *Ahead* counts the units on which the other method scores higher. Insulation is computed on the signed log observed/expected data; GenomeDISCO and HiC-Spector require non-negative input and are given its exponential. The last two rows before the chromosome comparison are reported to document that these two measures cease to discriminate below one megabase.

| Comparison | $n$ | Other | Hi-cGAN | Diff. | 95% CI | $p$ | Ahead |
| --- | --- | --- | --- | --- | --- | --- | --- |
| Pearson (log obs/exp) | 411 | 0.506 | 0.238 | 0.268 | [0.247, 0.289] | < 0.001 | 371/411 |
| Spearman | 411 | 0.488 | 0.232 | 0.257 | – | – | 369/411 |
| Insulation correlation | 411 | 0.492 | 0.297 | 0.195 | [0.173, 0.217] | < 0.001 | 334/411 |
| Boundary agreement | 276 | 0.402 | 0.224 | 0.178 | [0.151, 0.205] | < 0.001 | 204/276 |
| GenomeDISCO | 411 | 0.929 | 0.907 | 0.022 | – | – | 348/411 |
| HiC-Spector | 411 | 0.103 | 0.107 | –0.004 | – | – | 172/411 |
| vs HiC-Reg, per chromosome | 11 | 0.818 | 0.609 | 0.209 | [0.156, 0.255] | < 0.001 | 11/11 |

### S6.3 Invariance to metric and value space

The ordering does not depend on the choice of metric or on the sign convention of the target (Table S6). Moving to a rank-based coefficient costs Akita 0.017 and Hi-cGAN 0.006, so the difference is not carried by a small number of extreme pixels, and moving to the strictly positive ratio space changes the gap by 0.003. Hi-cGAN is negatively correlated with the measured map on 48 windows and Akita on 15; the tenth percentile of the per-window distribution is *−*0.013 for Hi-cGAN and 0.110 for Akita, while the ninetieth percentiles are 0.520 and 0.761 (Figures S5 and S6; five windows spanning the range of joint difficulty are drawn in Figure S7).

**Table S6:** The same predictions and the same target scored three ways over 411 windows. The ratio column is the exponential of the log observed/expected data, which is strictly positive and therefore admissible to methods that reject negative input; the transformation is monotone and bijective. Spearman’s coefficient depends only on rank and is consequently identical in both value spaces by construction, which serves here as a check that the two spaces contain the same data.

|  | Pearson (log obs/exp) | Pearson (ratio) | Spearman |
| --- | --- | --- | --- |
| Akita | 0.506 | 0.503 | 0.488 |
| Hi-cGAN | 0.238 | 0.238 | 0.232 |
| Difference | 0.268 | 0.265 | 0.257 |
| Akita ahead | 371/411 | 375/411 | 369/411 |

### S6.4 Validation of the per-window SCC

The reference hicrep implementation cannot be restricted to a sub-chromosomal region, so the SCC reported in the main text was computed with our own implementation and validated against the reference. Pairs of cool files were constructed at 448 bins and 2048 bp, matching Akita’s cropped window, with a controlled fraction of shared structure superimposed on a distance decay, and scored both by the hicrep binary and by the implementation used here; the two agree to better than 5 *×* 10*^−^*^3^ (Table S7).

Two details of the reference implementation are necessary for agreement and are easy to miss. The stratum weight is *N_k_ · varV stran*(*N_k_*) = (*N_k_* + 1)*/*12, a function of the number of entries in the stratum alone; substituting the observed standard deviations instead produces a different statistic, disagreeing by up to 0.48. The band is also trimmed before the mean filter is applied rather than after, so the filter sees zeros outside the retained band, and the filter’s divisor is purely geometric rather than a count of the present neighbours.

**Figure S5:**
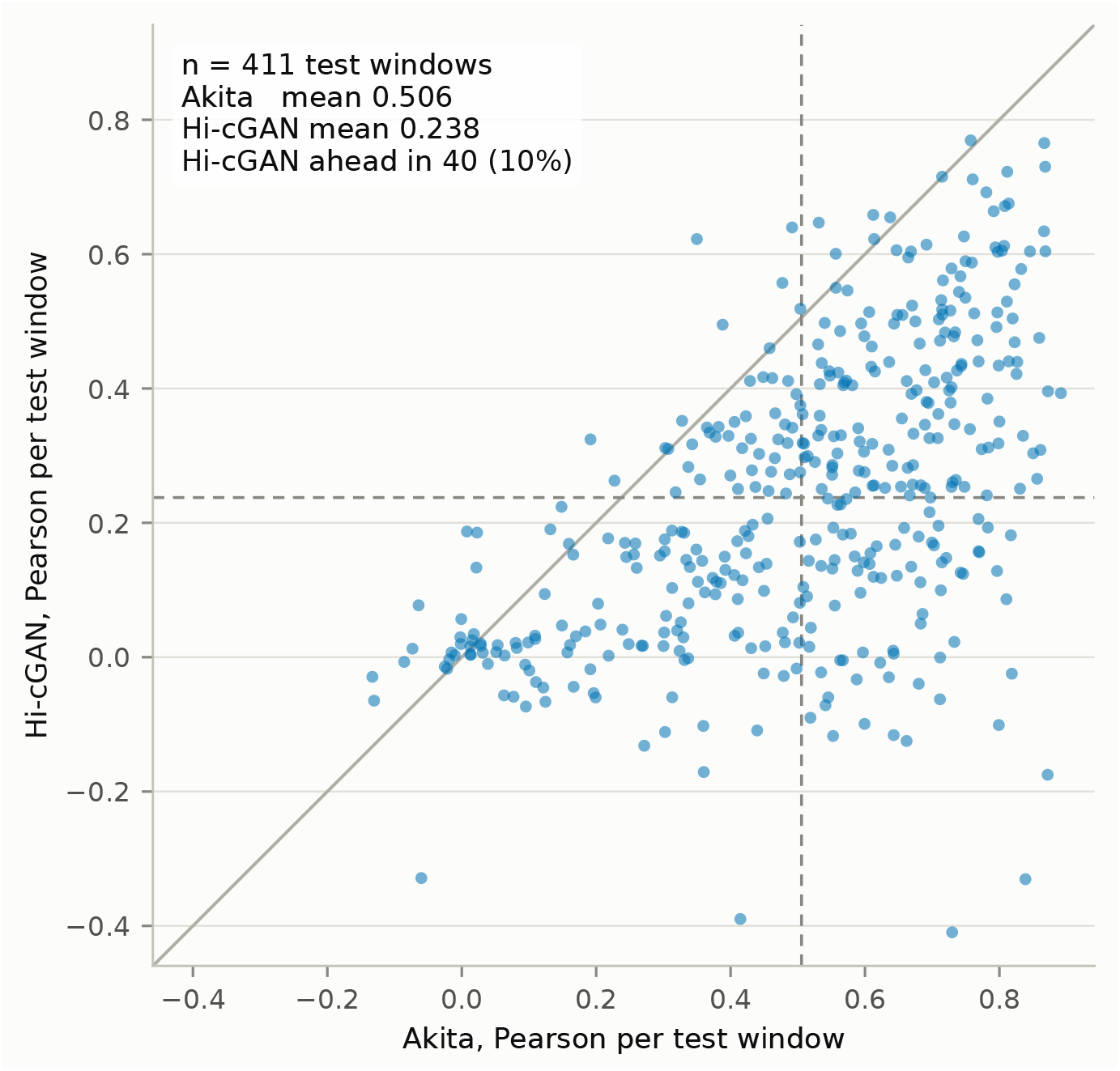
Hi-cGAN against Akita, one point per held-out window. With the line of equality. Dashed lines mark the two means. Points above the diagonal are windows Hi-cGAN predicts more accurately than Akita does.

**Figure S6:**
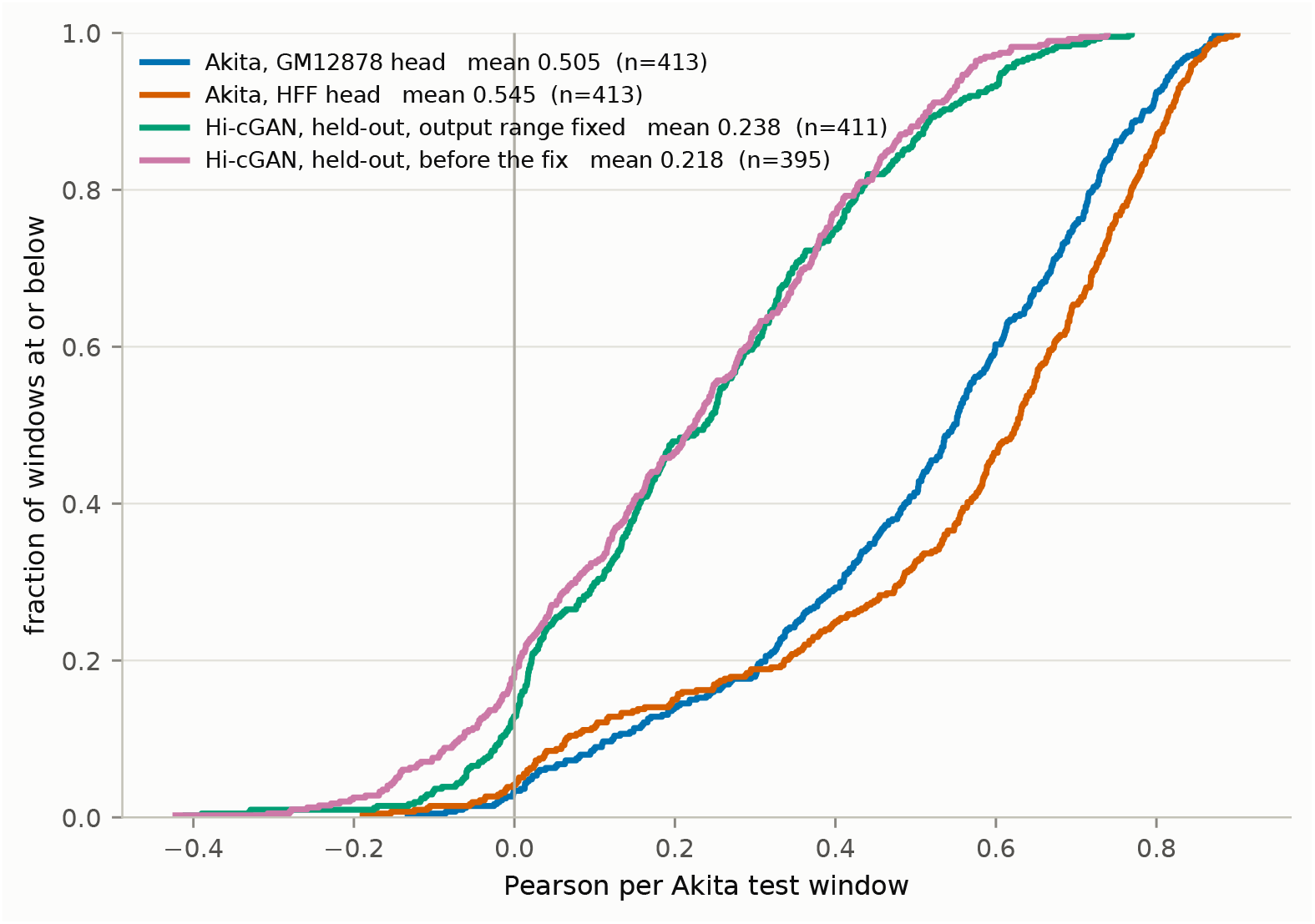
Distribution of per-window accuracy. Empirical cumulative distribution of the per-window Pearson correlation for each model, which compares them at every quantile rather than only at the mean.

**Figure S7:**
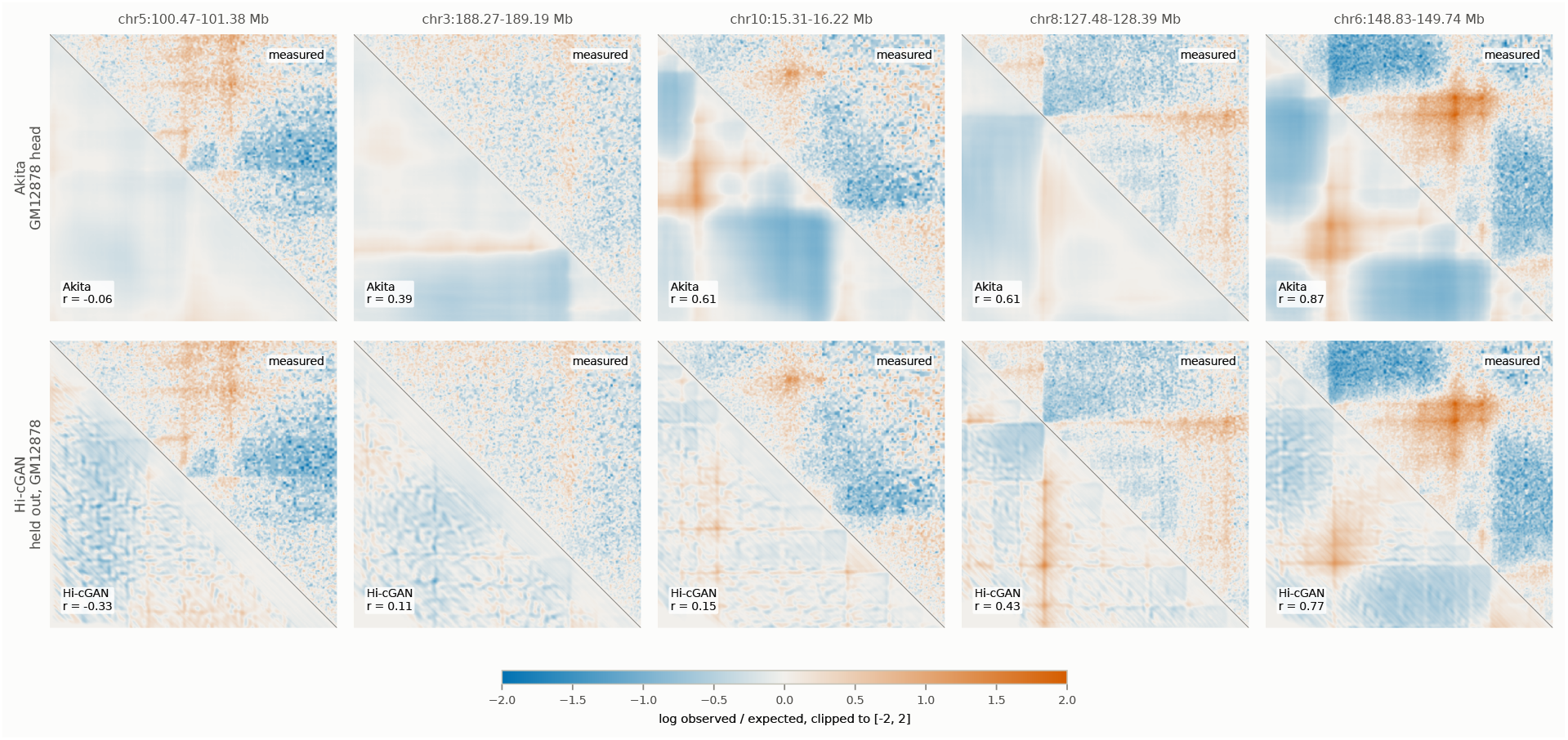
Five held-out windows across the range of joint difficulty. Measured map upper-right, prediction lower-left, shared scale. In contrast to Figure S10, which selects one window per outcome, these five span the range of joint difficulty from the hardest to the easiest.

**Table S7:** Per-window SCC against the reference implementation, on matched pairs of cool files with a controlled fraction of shared structure. Agreement is better than 5 *×* 10*^−^*^3^ throughout and better than 3 *×* 10*^−^*^4^ at *h* = 1.

| Shared | $h$ | hicrep | This work | Difference |
| --- | --- | --- | --- | --- |
| 0.0 | 1 | 0.008501 | 0.008338 | $1.6 \times 10^{-4}$ |
| 0.0 | 5 | 0.111092 | 0.114691 | $3.6 \times 10^{-3}$ |
| 0.3 | 1 | 0.130166 | 0.130264 | $9.8 \times 10^{-5}$ |
| 0.3 | 5 | 0.230889 | 0.235763 | $4.9 \times 10^{-3}$ |
| 0.6 | 1 | 0.576713 | 0.576948 | $2.4 \times 10^{-4}$ |
| 0.6 | 5 | 0.613530 | 0.616157 | $2.6 \times 10^{-3}$ |
| 0.9 | 1 | 0.935100 | 0.935161 | $6.1 \times 10^{-5}$ |
| 0.9 | 5 | 0.937818 | 0.937890 | $7.2 \times 10^{-5}$ |
| 1.0 | 1 | 0.999970 | 1.000000 | $3.0 \times 10^{-5}$ |
| 1.0 | 5 | 1.000000 | 1.000000 | 0 |

**Figure S8:**
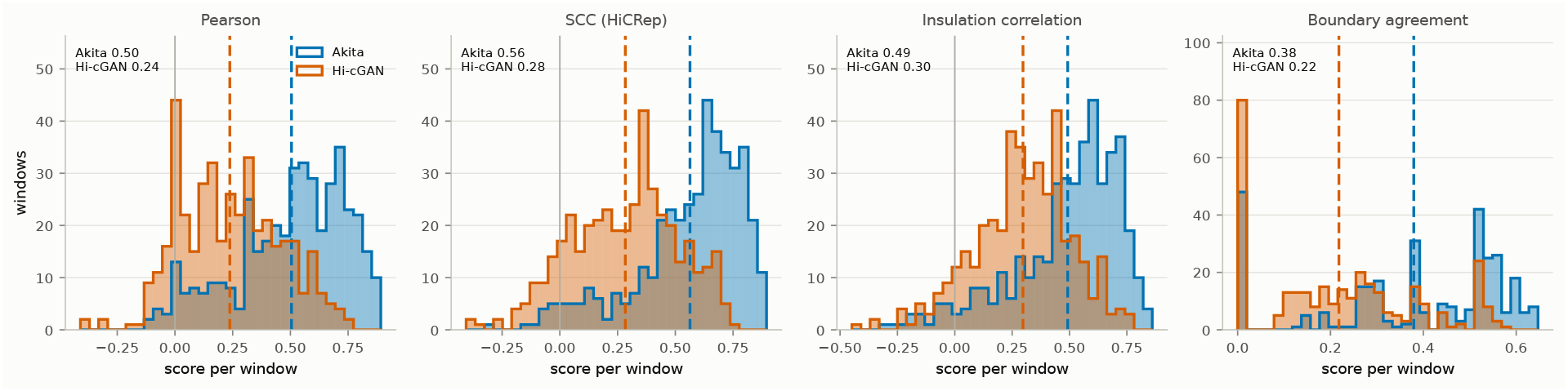
Distribution of each measure over the held-out windows. Akita and Hi-cGAN, one panel per measure, with the means marked by dashed lines. GenomeDISCO and HiC-Spector are omitted: below one megabase they compress every comparison into a narrow band, as reported in the main text.

**Figure S9:**
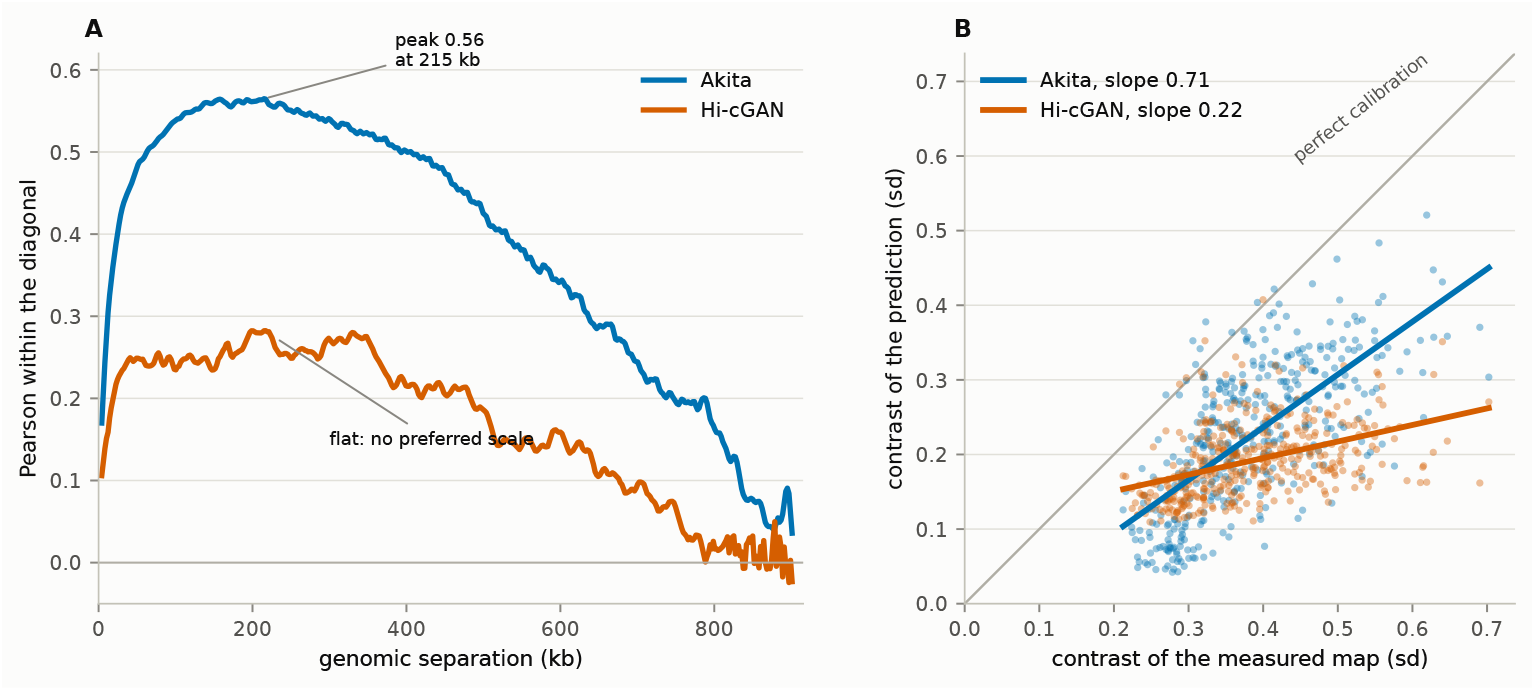
Decomposition of the Akita and Hi-cGAN difference. Over the 411 held-out windows. (A) Pearson correlation within each diagonal against genomic separation, averaged over windows. Akita peaks at 0.56 at 215 kb, the scale of topologically associating domains; Hi-cGAN stays near 0.25 from 50 to 400 kb and has no separation at which it does better. (B) Contrast of the prediction against contrast of the measured map, one point per window, with fitted slopes. A model whose output contrast tracks the target lies along the diagonal. A flat line does not mean the output is the same everywhere, only that its amplitude barely responds to how much structure the window holds.

**Figure S10:**
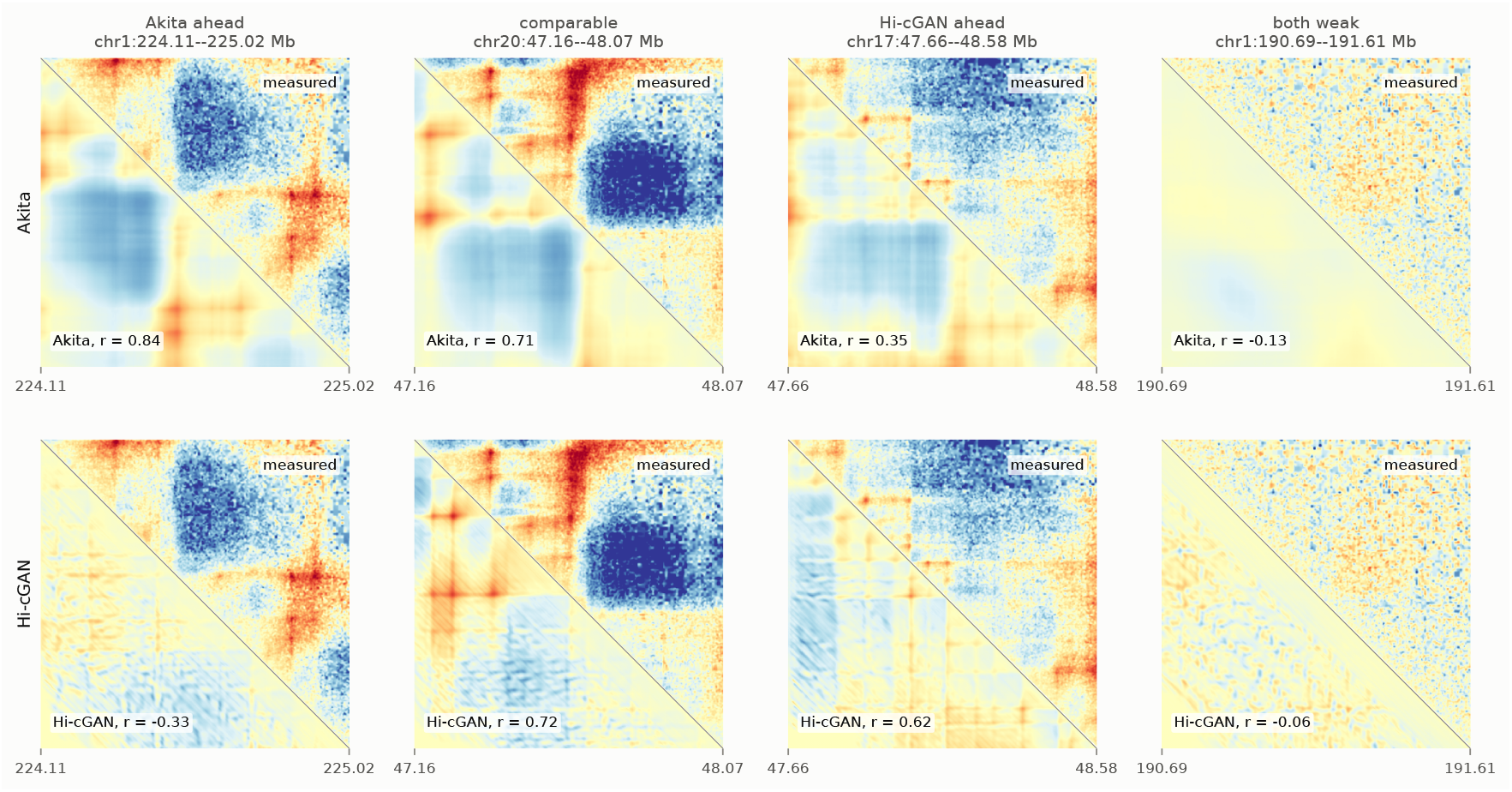
Held-out windows, one per outcome. Measured map upper-right, prediction lower-left, shared scale. Windows were selected by rule: the largest lead for each method restricted to windows that method predicts above its own median, the smallest difference among windows where both exceed their own median, and the window at which the better of the two methods is weakest. Both rows show GM12878 scored against the target carried in Akita’s published test records.

### S6.5 Decomposition by genomic separation

Splitting the per-window correlation by genomic separation shows that the difference between the two methods is not uniform across scales. Table S8 gives the mean within each band. The difference is largest at the scale of topologically associating domains and smallest at the shortest separations and beyond 717 kb, where both methods approach zero.

**Table S8:**
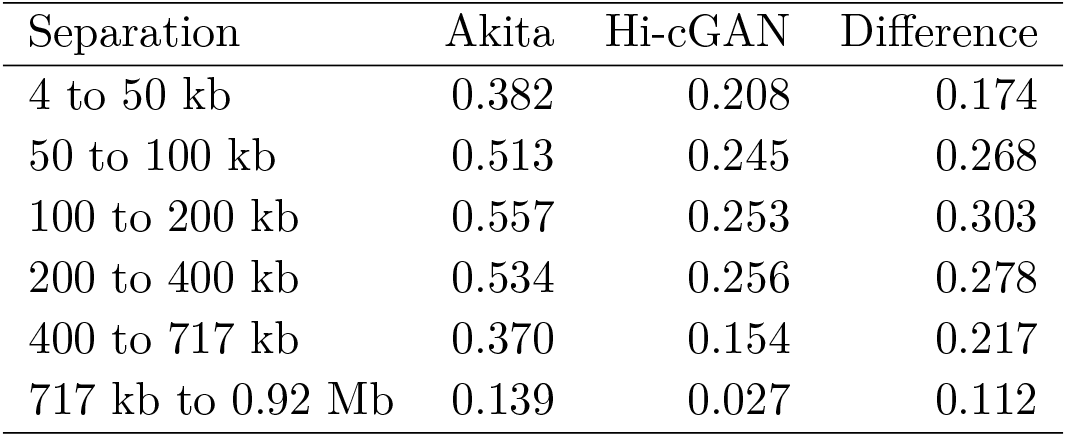
Per-window Pearson correlation split by genomic separation, averaged over the 411 held-out windows. Both methods are scored on identical pixels within every band. Akita’s advantage is largest at the scale of topologically associating domains and smallest at the shortest separations and beyond 717 kb, where both approach zero.

### S6.6 Calibration

Contrast here is the standard deviation of a window’s values over the 99,681 scored pixels, of the measured map or of a prediction as stated. Restricting to the 123 windows whose measured contrast falls in the middle third of its range holds the difficulty of the target roughly fixed and isolates each method’s residual failure mode. Within that band, Akita’s twenty weakest windows average a correlation of 0.239 and its twenty strongest 0.732, while the standard deviation of its output varies only from 0.199 to 0.257, a factor of 1.29. The corresponding figures for Hi-cGAN are *−*0.111 and 0.523, with output contrast from 0.196 to 0.225, a factor of 1.15. Neither model therefore fails by producing an output that lacks contrast; both fail by producing structure in the wrong place, and the average correlation of Akita’s twenty weakest windows remains positive while Hi-cGAN’s does not.

Across all windows and without that restriction, the relationship between measured contrast and accuracy differs, and it underlies the calibration statement in the main text. The rank correlation between the contrast of the measured map and the model’s accuracy is 0.789 for Akita and 0.524 for Hi-cGAN, and the standard deviation of Akita’s output falls to 0.083 on its forty weakest windows from 0.345 on its forty strongest, against 0.182 and 0.246 for Hi-cGAN, on a scale where the measured map averages 0.382.

## S7 Contact maps in contact space

The three regions of the main text’s map figure, drawn in the ordinary contact-map view with the distance decay left in. Every prediction is returned to a contact-count scale through the expected profile of the measured map. Akita is absent: its normalization carries no count scale.

**Figure S11:**
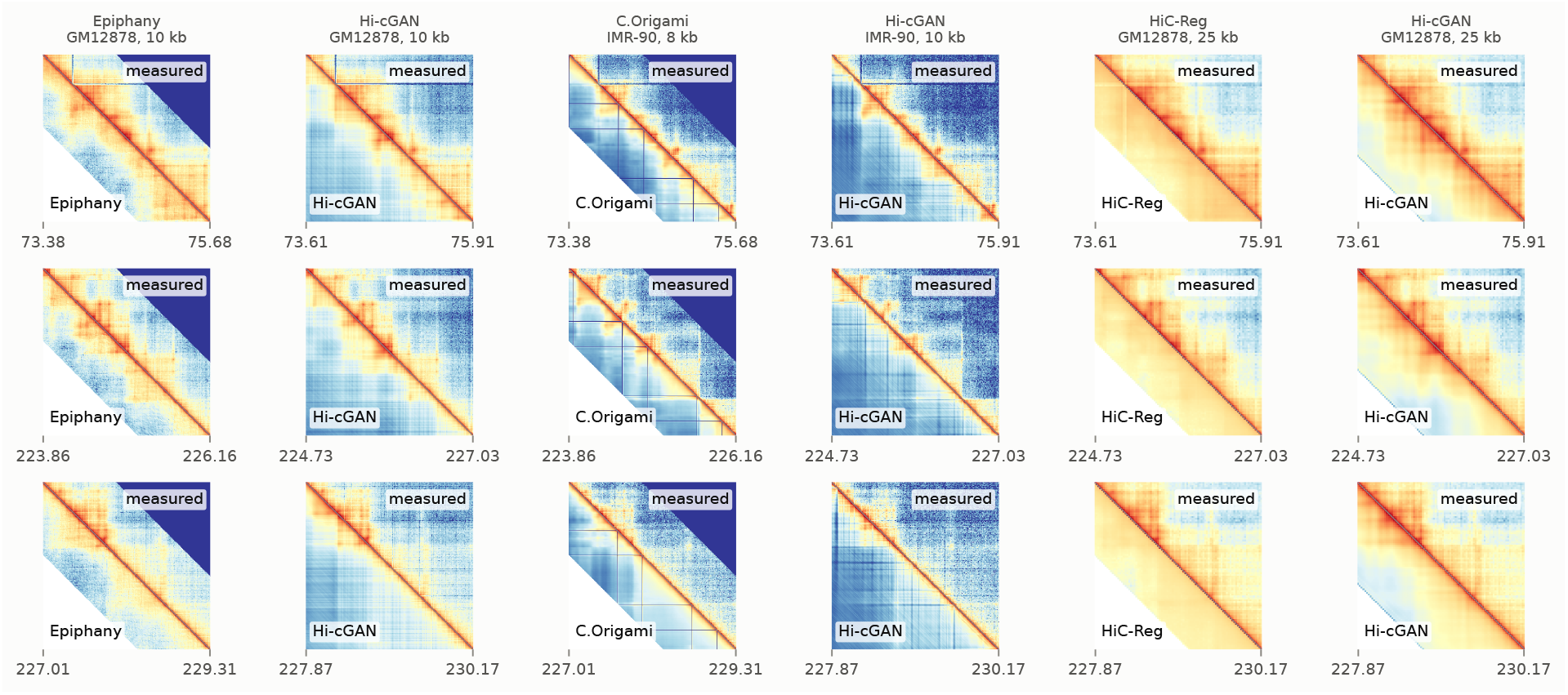
Contact maps in contact space. The same three regions of chromosome 2 as in the main text, one row per region, the competing method and Hi-cGAN at the same bin size in adjacent columns, drawn as log10(count+1) on the measured map’s scale. Bins a method cannot reach are white. Akita is absent: its normalization carries no count scale.

## S8 Called features: TAD boundaries and loops

The measures of the main text compare matrices. Here the features called from a predicted matrix are compared with those called from the measured matrix. Boundaries and loops were called with hicFindTADs and hicDetectLoops Wolff et al. (2018) on the measured and the predicted matrix under identical parameters, and matched within a tolerance of *t* bins. Hi-cGAN is last on boundaries in every block.

Loop recovery is poor for every method except C.Origami: Hi-cGAN reaches an *F*_1_ of 0.052 to 0.077, Epiphany 0.130, and HiC-Reg 0.068 only because it calls almost nothing, an average of 6.9 loops where the measured matrix holds 134, so its precision is high and its recall near zero. Only C.Origami’s loop *F*_1_ of 0.343 would support a loop analysis.

Boundaries separate the methods, and Hi-cGAN is last in all three blocks. Since a boundary is a local minimum of the insulation profile and the depth of that minimum scales with how far the map departs from its local average, a profile correlating at 0.851 whose boundaries match at an *F*_1_ of 0.355 would be explained if the profile had the right shape and the wrong amplitude. We tested that explanation directly.

**Figure S12:**
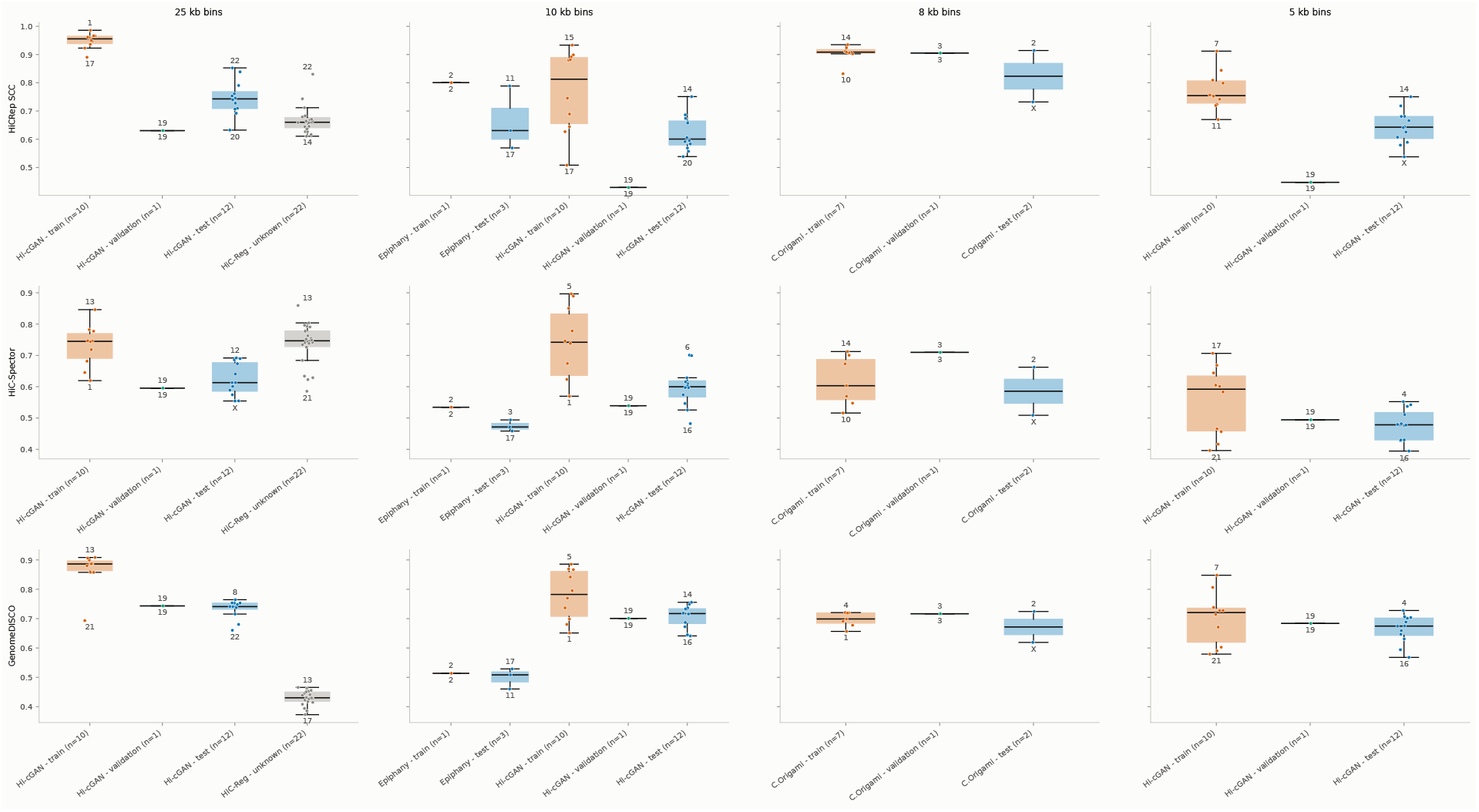
Per-chromosome scores for every method, all three measures. HiCRep SCC, HiC-Spector and GenomeDISCO per chromosome for every method at every resolution it exists at, split into the chromosomes each method trained on and those it did not.

**Table S9:**
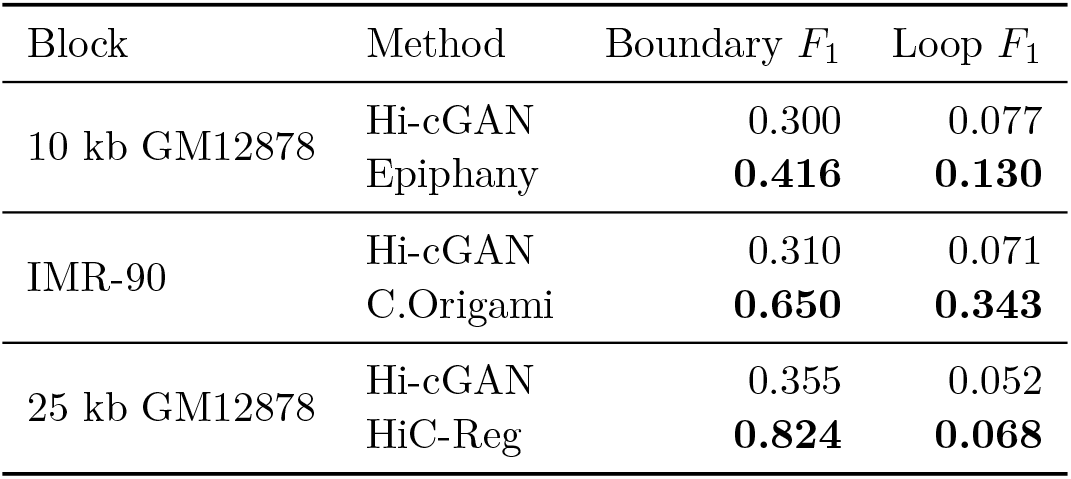
Boundary and loop recovery, tolerance two bins. *F*_1_ against the features called on the measured matrix with the same tool and parameters. A tolerance of two bins is 20 kb in the 10 kb blocks and 50 kb in the 25 kb block, so the physical strictness of the match differs between them and the blocks are not comparable with one another for that reason as well. The 25 kb row is the mean over the twelve held-out chromosomes; the 10 kb rows are chromosome 2. The split used for the released HiC-Reg model is not documented, so that row may be optimistic. Unlike the method battery of the main text, which scores each method at its own bin size, boundaries can only be matched between two call sets in shared coordinates, so C.Origami’s 8,192 bp output and the 8 kb measured map are both regridded onto a common 10 kb grid here.

| Block | Method | Boundary $F_1$ | Loop $F_1$ |
| --- | --- | --- | --- |
| 10 kb GM12878 | Hi-cGAN | 0.300 | 0.077 |
|  | Epiphany | **0.416** | **0.130** |
| IMR-90 | Hi-cGAN | 0.310 | 0.071 |
|  | C.Origami | **0.650** | **0.343** |
| 25 kb GM12878 | Hi-cGAN | 0.355 | 0.052 |
|  | HiC-Reg | **0.824** | **0.068** |

## S9 Post-hoc contrast rescaling

We define contrast as the standard deviation of the map on the clipped log observed/expected scale, as in the comparison against Akita. Writing *E*(*d*) for the mean of diagonal *d*, we rescaled each predicted map as

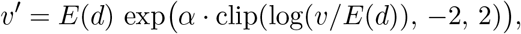

which expands (*α >* 1) or compresses (*α <* 1) the departure from the distance decay and leaves the decay itself in place, and called the boundaries again with hicFindTADs under the parameters of Table S9. The measured calls are unchanged between arms, so the only difference is the contrast of the prediction.

We used two choices of *α*. The first is a single number, fixed on chromosome 19 as the ratio of the measured to the predicted contrast there and applied unchanged to all twelve held-out chromosomes; it uses no held-out data, and a user could apply it to their own prediction. The second sets *α* per window from the measured contrast of the chromosome being predicted. This uses the test data and is not a usable method; we report it as the ceiling of any correction acting on contrast alone.

**Table S10:** Correcting the contrast does not recover the boundaries. Mean over the twelve held-out chromosomes at 25 kb; the measured matrices carry 259 boundaries per chromosome. The global exponent *α* = 0.88 is fixed on the validation chromosome and applied unchanged to every held-out chromosome. The oracle sets *α* per window from the measured contrast of the chromosome being predicted, which no user can do; it is the ceiling of any correction acting on contrast alone. Every difference is in the third decimal and the ordering changes with the tolerance.

|  | As trained | Global rescaling | Per-window oracle |
| --- | --- | --- | --- |
| Boundaries called per chromosome | 275 | 278 | 279 |
| Boundary $F_1$, tolerance 0 bins | 0.082 | 0.083 | **0.084** |
| Boundary $F_1$, tolerance 1 bin | **0.230** | **0.230** | 0.229 |
| Boundary $F_1$, tolerance 2 bins | **0.355** | 0.354 | 0.352 |
| Boundary $F_1$, tolerance 3 bins | **0.472** | 0.470 | 0.467 |
| Boundary $F_1$, tolerance 4 bins | 0.567 | **0.568** | 0.566 |
| Insulation profile Pearson | 0.800 | **0.803** | 0.773 |

Neither arm moves the boundary calls. The *F*_1_ at a tolerance of two bins is 0.355 as trained, 0.354 under the validation-fitted correction and 0.352 under the oracle, and the ordering is the same at every tolerance from zero to four bins. The oracle lowers the insulation correlation, from 0.800 to 0.773, because scaling each window by its own factor perturbs the profile across window seams without improving where its minima fall.

The boundary counts indicate why. Hi-cGAN calls 275 boundaries per chromosome against 259 in the measured map, so it places boundaries wrongly rather than missing them, and an amplitude correction cannot move a boundary. The contrast statistic measures a response, not a deficit: over the 1,939 windows of the twelve held-out chromosomes at 25 kb, predicted contrast regresses on measured contrast with a slope of 0.250 at a correlation of 0.300, close to the 0.22 we obtain at 2 kb, while the mean predicted contrast of 0.741 *exceeds* the measured 0.543. The model emits structure at a roughly constant amplitude whatever the region contains, which is a failure to locate structure and not to scale it.

We therefore expect no gain from a variance-matching output transform, a loss term penalising the difference of standard deviations, or any other device acting on the amplitude of the prediction: the ceiling for that family is an *F*_1_ of 0.352, below the 0.355 the model already reaches. Improving the boundary calls requires the structure to be placed correctly, which bears on the generator’s inputs and receptive field rather than on its calibration.

## S10 Training set size

The sampler slides by one bin, so two consecutive training positions differ by a single bin of target. Counting positions therefore measures how often a training example is presented rather than the amount of distinct data, so both counts are reported alongside the generator’s parameter count, which is in the tens of millions.

The 10 kb configuration has the fewest non-overlapping windows (Table S11). A window of 512 bins spans 5.12 Mb, so only 251 of them tile the ten training chromosomes, and the model fitted to them has 58.7 million parameters. At 25 kb the window of 64 bins spans 1.6 Mb and tiles 857 times, and the network fitted to it is smaller; the 5 kb configuration tiles 526 times, between the two.

**Table S11:** Training data available in each configuration. On the ten training chromosomes of GM12878, for the configurations used in the main text. A chromosome of *n* bins offers *n −* 3*w* sliding positions and holds (*n −* 2*w*)*/w* non-overlapping target windows. No coverage filter was applied to these sweeps, so the counts are geometric. Neither column is an exact count of independent samples: the first greatly overstates it, and the second still counts as independent two windows that sit inside one compartment.

| Bin size | Window | Sliding positions | Non-overlapping windows | Unique target bins | Generator parameters |
| --- | --- | --- | --- | --- | --- |
| 25 kb | 64 | 54,508 | 857 | 55,148 | 33,428,609 |
| 10 kb | 512 | 125,702 | 251 | 130,822 | 58,713,665 |
| 5 kb | 512 | 266,760 | 526 | 271,880 | 58,713,665 |

The non-overlapping count is a counting convention that removes direct overlap; it is not a statistical effective sample size, which would have to account for the correlation that remains between windows sharing a compartment or a replication-timing domain, and which would be smaller again. The ratio of parameters to samples is not a measure of overfitting risk in a convolutional network, where weights are shared across positions, the objective is regularized and the target has strong spatial structure; a network with more parameters than samples in this sense is ordinary rather than pathological. The two columns bracket how training data can be counted, and we draw no formal conclusion from either endpoint. They do support a qualitative statement that the rest of the article bears out: the gap between a model’s accuracy on its training chromosomes and on the held-out ones is wide throughout, and differences of 0.01 to 0.03 between input factors rest on little independent data.

## S11 Contribution of individual chromatin factors

### S11.1 Single factors

Fourteen models were trained, one per chromatin factor, at three bin sizes and up to four window sizes, giving 125 configurations. Table S13 gives the held-out SCC of each factor at 25 kb with a window size of 64, averaged over the twelve held-out chromosomes. Figures S13, S14 and S15 give the per-chromosome distributions behind these means for every factor at every bin and window size, separated into training and held-out chromosomes.

**Figure S13:**
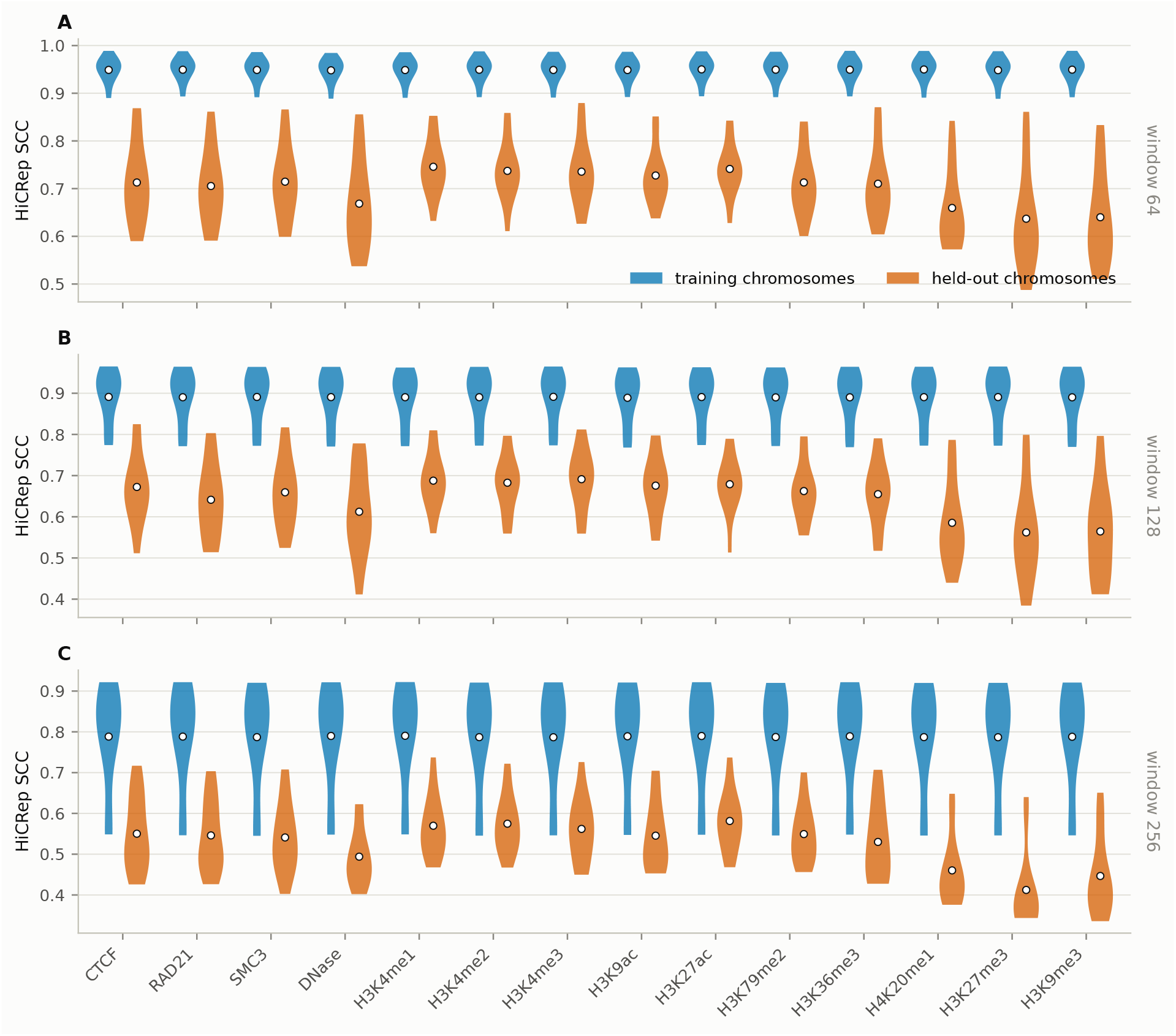
Per-chromosome accuracy of each chromatin factor used alone, 25 kb. HiCRep SCC per chromosome at window sizes 64, 128 and 256, one violin pair per factor: the training chromosomes and the held-out chromosomes, the white dot marking the mean of each. The validation chromosome 19 belongs to neither group and is not drawn.

**Figure S14:**
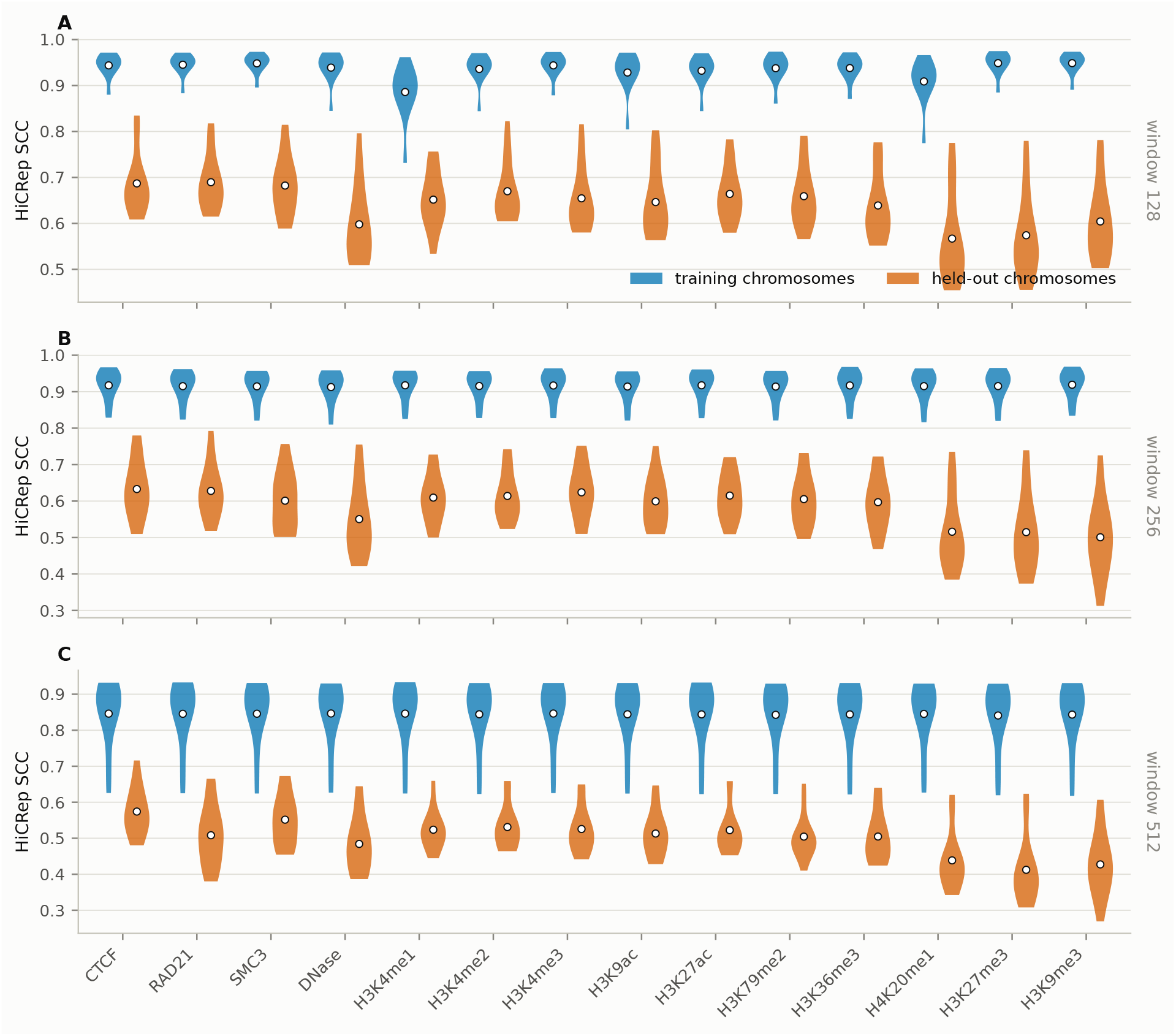
Per-chromosome accuracy of each chromatin factor used alone, 10 kb. As in Figure S13, at window sizes 128, 256 and 512.

**Figure S15:**
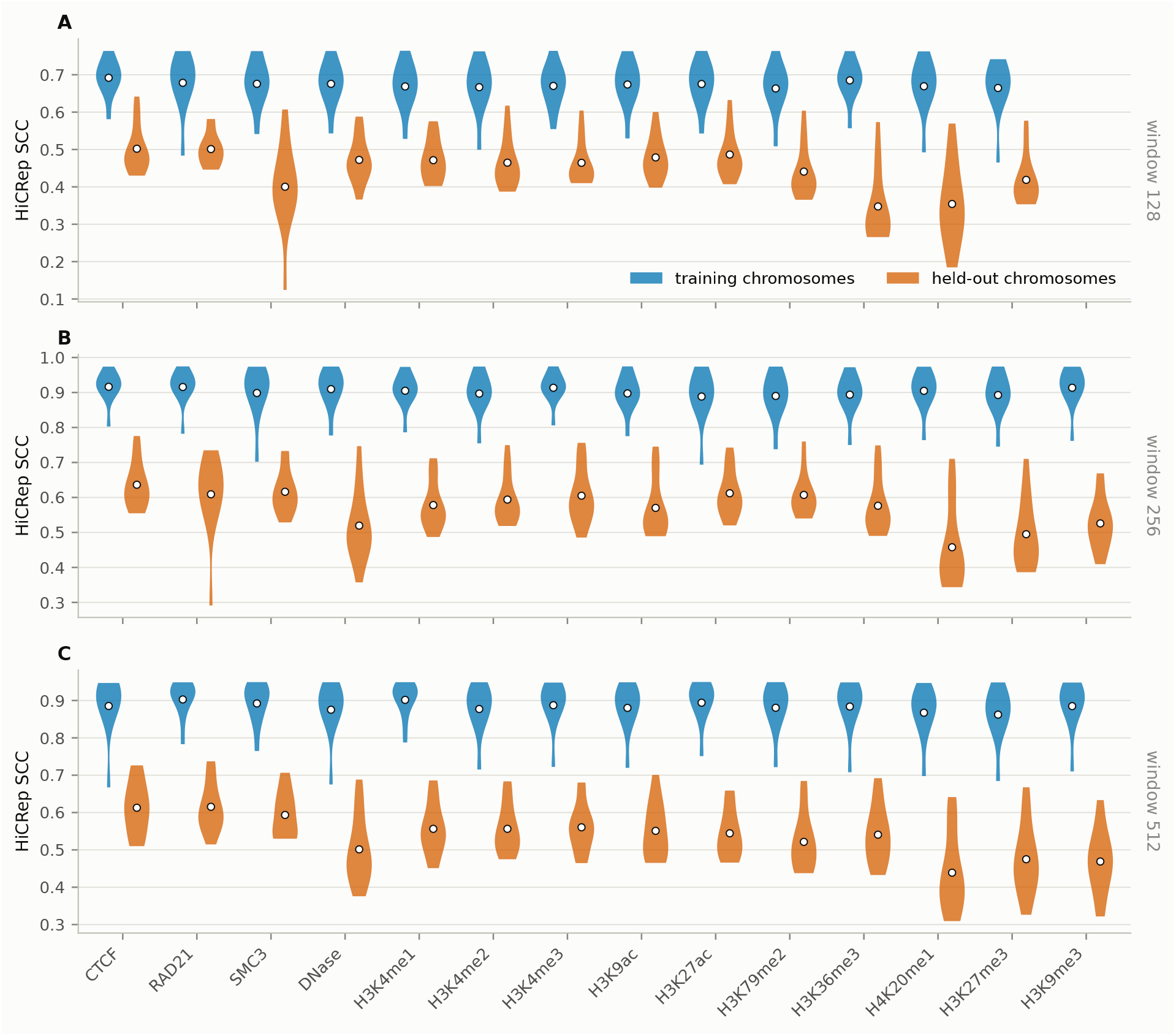
Per-chromosome accuracy of each chromatin factor used alone, 5 kb. As in Figure S13, at window sizes 128, 256 and 512. RAD21 was not trained at a window of 128 at this bin size.

The spread across factors is small, 0.637 for the weakest against 0.746 for the strongest, so every one of the fourteen carries a usable amount of the signal. Which factor is best depends on the resolution being predicted, and the way it changes has a straightforward biological reading.

### S11.2 Dependence of the best factor on the resolution

Table S12 gives the leading factor for each of the nine configurations in the sweep. Ranks rather than raw scores are compared across configurations, because SCC is not on the same scale at different bin sizes.

**Table S12:** Leading chromatin factor per resolution. Leading and second factor for each configuration in the single-factor sweep, with the rank of CTCF among the fourteen. At 5 kb and 10 kb the architectural proteins lead and CTCF is first or second in every case; at 25 kb the active histone marks lead and CTCF falls to fifth or eighth.

| Configuration | Best factor | SCC | Runner-up | SCC | CTCF rank |
| --- | --- | --- | --- | --- | --- |
| 5 kb, w128 | CTCF | 0.502 | SMC3 | 0.501 | 1 |
| 5 kb, w256 | CTCF | 0.636 | SMC3 | 0.616 | 1 |
| 5 kb, w512 | RAD21 | 0.615 | CTCF | 0.613 | 2 |
| 10 kb, w128 | RAD21 | 0.690 | CTCF | 0.687 | 2 |
| 10 kb, w256 | CTCF | 0.633 | RAD21 | 0.628 | 1 |
| 10 kb, w512 | CTCF | 0.574 | SMC3 | 0.552 | 1 |
| 25 kb, w64 | H3K4me1 | 0.746 | H3K27ac | 0.741 | 8 |
| 25 kb, w128 | H3K4me3 | 0.691 | H3K4me1 | 0.688 | 6 |
| 25 kb, w256 | H3K27ac | 0.581 | H3K4me2 | 0.575 | 5 |

The separation by resolution is consistent across all nine configurations. At 5 kb and 10 kb the top two places are taken by CTCF and the cohesin subunits RAD21 and SMC3 in all six configurations, and CTCF is first or second in every one of them. At 25 kb the leaders are H3K4me1, H3K4me3 and H3K27ac, and CTCF drops to rank five to eight. The repressive marks H3K9me3 and H3K27me3 are last almost everywhere, at ranks 12 to 14, and DNase accessibility is consistently poor at rank 11 or 12 regardless of resolution.

### S11.3 Interpretation

The crossover follows the biology of what a Hi-C matrix contains at each scale.

At 5 kb and 10 kb the resolvable features are individual loop anchors and TAD boundaries. These are physically created by CTCF-bound sites and the cohesin ring, and the current model of their formation is loop extrusion, in which cohesin extrudes chromatin until it is halted at convergently oriented CTCF sites Rao et al. (2014, 2017). At this resolution a CTCF peak corresponds nearly one to one with the feature to be predicted, and the network effectively receives the anchor positions as input. That RAD21 and SMC3, the two cohesin subunits in the panel, occupy the remaining top places is consistent with the same mechanism rather than being an independent finding: cohesin is retained where CTCF halts it, so the three tracks mark the same sites.

At 25 kb an individual anchor spans a fraction of one bin and the loops largely disappear into the diagonal. What survives at this scale is compartment organisation, the segregation of the genome into an active A and an inactive B compartment Lieberman-Aiden et al. (2009); Schwarzer et al. (2017). Compartment identity is not set by CTCF; it tracks transcriptional activity, and the marks that report activity are exactly the ones that come first at 25 kb: H3K4me1 at enhancers, H3K27ac at active enhancers and promoters, H3K4me3 and H3K9ac at active promoters. The network is no longer locating anchors but separating active from inactive chromatin, for which a CTCF track is close to uninformative.

The repressive marks H3K9me3 and H3K27me3 are last at nearly every configuration, which at first sight is surprising, since the B compartment is heterochromatic and might be expected to be predictable from them. It is consistent, however, with the compartments being close to a binary partition: once the A compartment is located from active marks, B is its complement, and a separate repressive track adds little. DNase accessibility performs poorly at every resolution, which we attribute to it being a narrow-peak signal reporting individual regulatory elements rather than either anchors or domain-scale state. The qualification is that the differences between neighbouring factors are small, frequently 0.005 or less, so the identity of the single best factor within a resolution should not be over-read; the pattern that matters is the group, architectural against active, not the ordering inside it.

Input tracks should be chosen for the resolution being predicted. If loops and TAD boundaries at 5 to 10 kb are the target, CTCF or a cohesin subunit is the single most useful track. If the aim is compartment-scale structure at 25 kb or coarser, an active mark such as H3K27ac or H3K4me1 is the better choice, and CTCF is not. CTCF with DNase is a reasonable two-track choice across resolutions, but it is not the best choice at any single one.

**Table S13:**
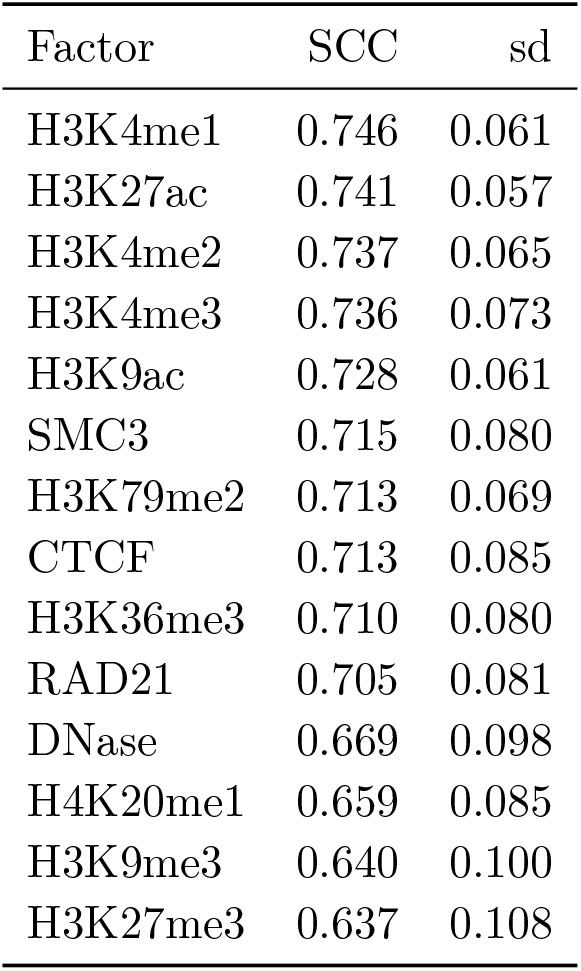
Held-out SCC of each chromatin factor used alone. GM12878 at 25 kb with a window size of 64, mean and standard deviation over the twelve held-out chromosomes.

| Factor | SCC | sd |
| --- | --- | --- |
| H3K4me1 | 0.746 | 0.061 |
| H3K27ac | 0.741 | 0.057 |
| H3K4me2 | 0.737 | 0.065 |
| H3K4me3 | 0.736 | 0.073 |
| H3K9ac | 0.728 | 0.061 |
| SMC3 | 0.715 | 0.080 |
| H3K79me2 | 0.713 | 0.069 |
| CTCF | 0.713 | 0.085 |
| H3K36me3 | 0.710 | 0.080 |
| RAD21 | 0.705 | 0.081 |
| DNase | 0.669 | 0.098 |
| H4K20me1 | 0.659 | 0.085 |
| H3K9me3 | 0.640 | 0.100 |
| H3K27me3 | 0.637 | 0.108 |

### S11.4 Combinations

The forward selection extended the best single factor of each bin size by candidate second, third and fourth factors drawn from the top of the single-factor ranking. Table S14 gives the best set at each number of inputs and Figure S16 every set trained; the outcome is the reason the main text uses one or two tracks and not fourteen.

**Table S14:**
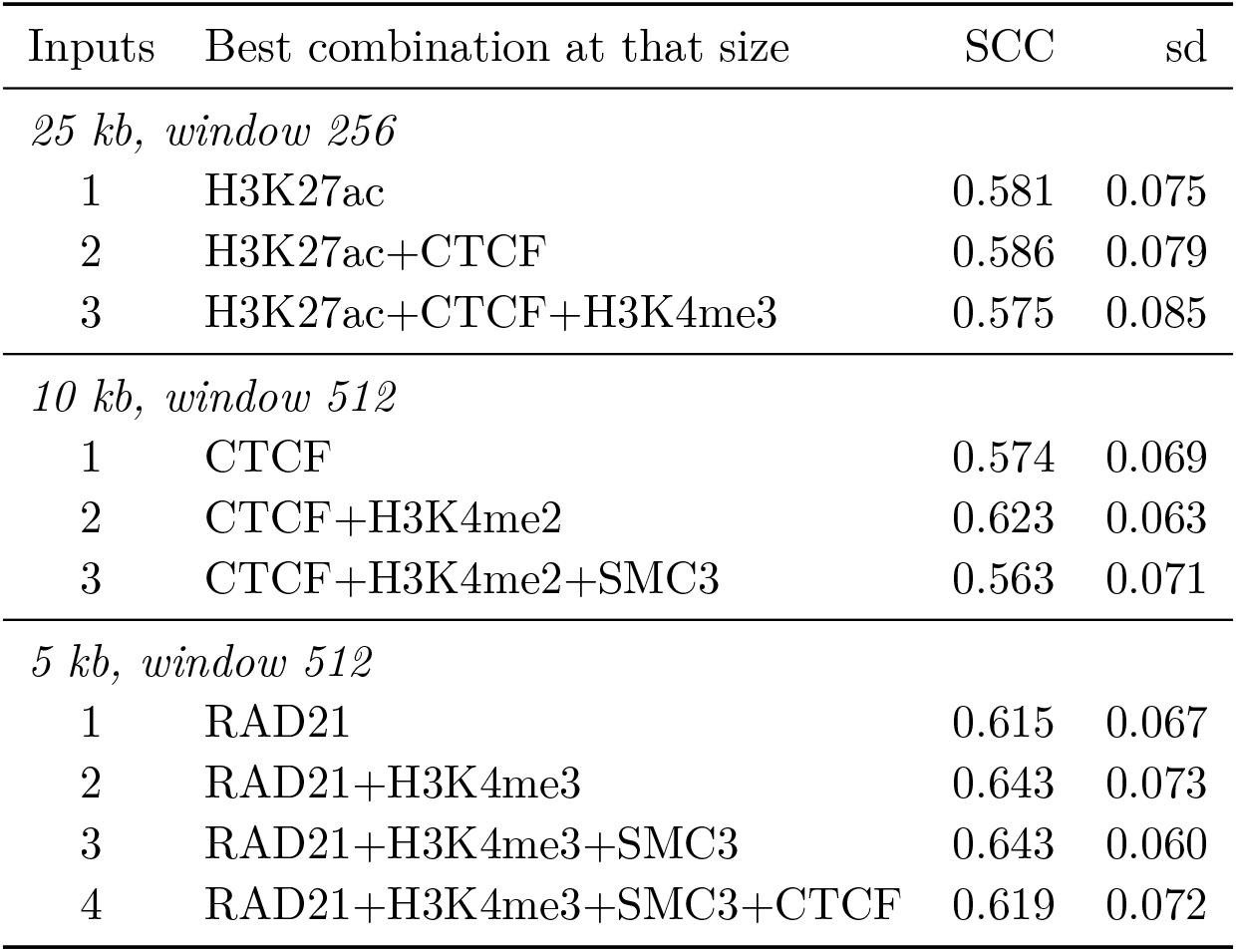
Accuracy against the number of input factors. The best combination at each number of inputs for each bin size, held-out SCC with its standard deviation over the held-out chromosomes. Combinations were trained at one window per bin size, so rows within a block share one configuration and the comparison is not confounded by resolution or window size.

**Figure S16:**
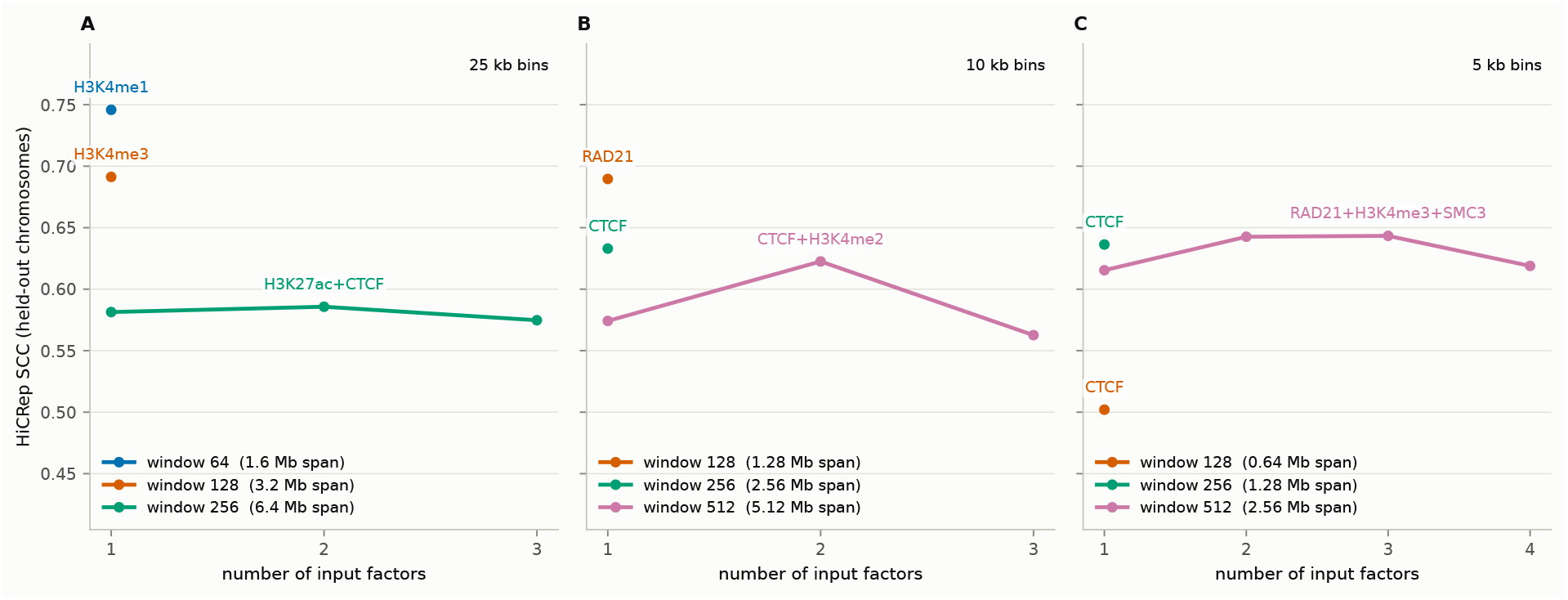
Accuracy against the number of input factors. Held-out SCC of the best set at each number of chromatin factors, one panel per bin size, one line per window size. The best point of each line is labelled with its factor set.

Accuracy does not increase with the number of inputs. The best pair beats the best single factor at every bin size, a third factor helps only at 5 kb, and there by 0.001, and the one extension to four factors, made at 5 kb, lowers the score.

This runs against the intuition that more information should help. The chromatin tracks are strongly correlated with one another, so an additional track adds little that is genuinely new while adding parameters to the embedding network; the number of training samples is also fixed by the genome, so a wider input is fitted from the same data. The two effects cannot be separated here. Either way, one or two tracks are enough, which also makes the method applicable to cell types for which few ChIP-seq experiments exist.

## S12 Which data selected each choice

The factor ranking uses held-out scores. This section reports which data settled every other choice in the study (Table S15).

**Table S15:**
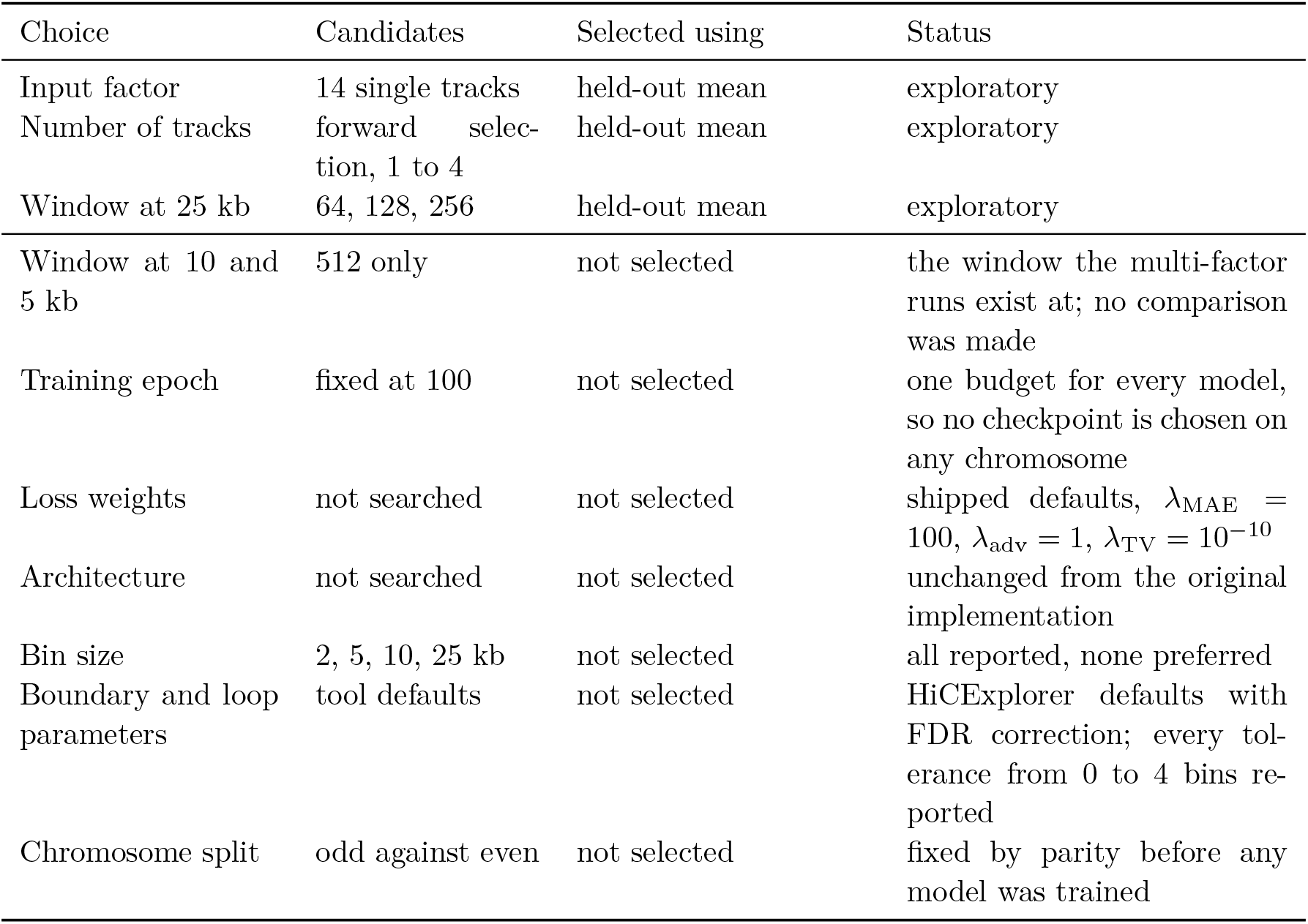
Provenance of each choice. *Held-out mean* is the HiCRep SCC averaged over the twelve held-out chromosomes, the even ones and X, at the fixed checkpoint. The three entries above the rule used those chromosomes and are therefore exploratory; their reported accuracies are optimistic and Table S16 measures by how much. Everything below the rule was fixed without reference to the held-out chromosomes, either because it was pre-specified or because only one candidate existed.

A configuration may be *fixed in advance* and then evaluated on held-out data, which is a clean estimate and is what the comparison against Akita is. It may be *selected on validation data* and then evaluated on held-out data, which is also clean, and is what the validation column of Table S16 reconstructs. Or it may be *selected and evaluated on the same held-out data*, which is not a clean estimate at all, and is what the factor sweep of the main text does. Only the first two are confirmatory. The distinction is not whether a number was computed on held-out chromosomes, since all three are, but whether those chromosomes also chose the model being reported.

The three exploratory entries are related: all concern which inputs and window the model is given, and all three were ranked on held-out score. The window at 25 kb was chosen the same way as the factor, on held-out SCC, which is monotone in the window there and favours 64 outright. The window at 10 and 5 kb was not selected; 512 is the only window at which the multi-factor runs exist, so singles and combinations are compared at one window rather than across windows.

The comparison against Akita is not affected by these selection choices. Its configuration, its held-out intervals and its target were all fixed by Akita’s own published split before we trained anything, and there is no second collection of Akita test windows on which a post-hoc choice could have been fitted. The same holds for the loss ablation, whose three conditions differ only in a loss weight set in advance.

## S13 Cost of selecting the input factor on the test chromosomes

The factor rankings of the main text order the candidates by their held-out score, so the winner is chosen on the chromosomes it is then reported on. This section measures the consequence by replaying both selection rules on every configuration. No model is retrained: each candidate was already scored on every chromosome at the fixed checkpoint, so the validation rule, which ranks by the score on chromosome 19 alone, and the held-out rule, which ranks by the mean over the twelve held-out chromosomes, can both be applied to the same set of models. In each case we report the *held-out* mean of whichever model the rule picks, so the two columns are on the same scale and differ only in how the model was chosen.

**Table S16:** Both selection rules, every configuration. Each cell names the factor or combination the rule picks and, in brackets, the held-out HiCRep SCC of that model, averaged over the twelve held-out chromosomes at the fixed checkpoint. Both columns are held-out numbers; only the selection differs. The validation column is the unbiased one, chromosome 19 alone having chosen the model. *r_h_* is the rank of the validation-selected model in the held-out ordering and *r_v_* the rank of the held-out-selected model in the validation ordering, so a configuration in which both are small is one where the two rules nearly agreed. The rules pick the same model in two of the nine configurations, and where they differ the gap runs from 0.009 to 0.060, median 0.029 across all nine. No configuration had a tie at the top of either ranking; ties would be broken by candidate name. The held-out-selected configurations at 25 kb window 64, 10 kb window 512 and 5 kb window 512 are the ones tabulated in the method battery of the main text. Table S17 rescores the two GM12878 blocks with the validation-selected configurations instead, and it is those figures that the main text quotes when it compares against Epiphany and that the abstract reports. The difference column is the observed consequence of the two rules in that configuration, not a general estimate of selection bias.

| Bin | Win. | Cand. | Validation-selected | Held-out-selected | Diff. | $r_h$ | $r_v$ |
| --- | --- | --- | --- | --- | --- | --- | --- |
| 5 kb | 128 | 13 | CTCF (0.502) | CTCF (0.502) | 0.000 | 1 | 1 |
| 5 kb | 256 | 14 | H3K79me2 (0.607) | CTCF (0.636) | 0.029 | 5 | 2 |
| 5 kb | 512 | 24 | RAD21+CTCF (0.601) | RAD21+H3K4me3+SMC3 (0.643) | 0.042 | 12 | 15 |
| 10 kb | 128 | 14 | RAD21 (0.690) | RAD21 (0.690) | 0.000 | 1 | 1 |
| 10 kb | 256 | 14 | H3K4me2 (0.614) | CTCF (0.633) | 0.019 | 5 | 2 |
| 10 kb | 512 | 21 | CTCF+H3K4me2+SMC3 (0.563) | CTCF+H3K4me2 (0.623) | 0.060 | 5 | 4 |
| 25 kb | 64 | 14 | SMC3 (0.715) | H3K4me1 (0.746) | 0.031 | 6 | 4 |
| 25 kb | 128 | 14 | H3K4me2 (0.683) | H3K4me3 (0.691) | 0.009 | 3 | 8 |
| 25 kb | 256 | 25 | SMC3 (0.541) | H3K27ac+CTCF (0.586) | 0.045 | 20 | 3 |

The difference is not a constant and should not be treated as one. It is zero where the rules happen to agree and 0.060 at 10 kb with a window of 512, which is the configuration behind the comparisons against Epiphany and C.Origami. It is also not, strictly, an estimate of selection bias: it is the gap between the two models the two rules pick in that configuration; a proper bias estimate would require repeated splits we do not have.

The two ranks quantify the disagreement between the rules. In six of the nine configurations the held-out-selected model sits in the top four of the validation ordering, so the two rules were choosing between near neighbours. In the remaining three they were not: at 5 kb with a window of 512 each rule’s winner sits twelfth or fifteenth in the other’s ordering, and at 25 kb with a window of 256 the validation-selected model ranks twentieth of twenty-five on held-out score. The spread among candidates is small at every configuration, which is why so many of these choices are close and why the rules nonetheless disagree in seven of nine.

The number of tracks was itself settled by forward selection on the held-out mean, a sequence of adaptive decisions rather than one choice among fourteen, and is therefore the more strongly selected of the exploratory choices; each step of that search, with its score, is in the released results.

## S14 The GM12878 blocks under validation-only selection

The substitution is applied here to the blocks of the main text, on the same chromosomes, against the same measured matrices and through the same three scoring paths, with only the input configuration changed to the one chromosome 19 alone would have picked. Table S16 compares the two rules on the sweep-wide held-out mean instead.

**Table S17:** The GM12878 blocks scored with both input configurations. Held-out-selected rows use the configuration reported in the method battery of the main text; validation-selected rows use the configuration chromosome 19 would have chosen. Only the configuration differs; chromosomes, measured matrices and scoring code are identical, and both rows come from one run so that they are comparable with each other. Two entries differ from the main table in the third decimal, HiC-Spector 0.622 against 0.623 and GenomeDISCO 0.737 against 0.732 at 25 kb, the block means there having been assembled from separate runs of the three scoring paths. HiCRep is the measure the selection was made on, and it is the only one that falls consistently under the substitution.

| Block | Configuration | HiCRep | Spector | DISCO | Insul. | Bound. |
| --- | --- | --- | --- | --- | --- | --- |
| 25 kb, 12 chr | H3K4me1, held-out-selected | 0.746 | 0.622 | 0.733 | 0.851 | 0.055 |
|  | SMC3, validation-selected | 0.715 | 0.622 | 0.737 | 0.857 | 0.062 |
| 10 kb, chr2 | CTCF+H3K4me2, held-out-selected | 0.662 | 0.611 | 0.716 | 0.761 | 0.076 |
|  | CTCF+H3K4me2+SMC3, validation-selected | 0.640 | 0.638 | 0.748 | 0.752 | 0.059 |

At 25 kb, selecting on HiCRep raises HiCRep by 0.031 and improves no other measure: HiC-Spector is identical to three decimals, and GenomeDISCO, insulation and boundary agreement all favour the model that HiCRep ranks lower. A single statistic used for selection improves that statistic, and the four measures disagree about the ordering of two configurations of the same method in the same way they disagree about the ordering of different methods.

## S15 Maps produced by the linear baseline

The ridge baseline of the main text reaches a HiCRep SCC of 0.691 against Hi-cGAN’s 0.715, a difference small enough that the maps must be inspected rather than only scored.

The comparison is not close in the way the SCC suggests. The ridge output is striped: a bin carrying high signal raises every pixel that involves it, horizontally and vertically, which in observed/expected space appears as a lattice. Hi-cGAN produces blocks with interiors elevated relative to their borders, the defining feature of a domain, at approximately the measured positions.

This is a property of the measure as much as of the baseline: a map with the right marginal structure and no domains scores 0.691 on a statistic the field routinely reports as its primary measure of quality. The HiCRep score at this bin size is therefore not driven by the presence of domains.

**Figure S17:**
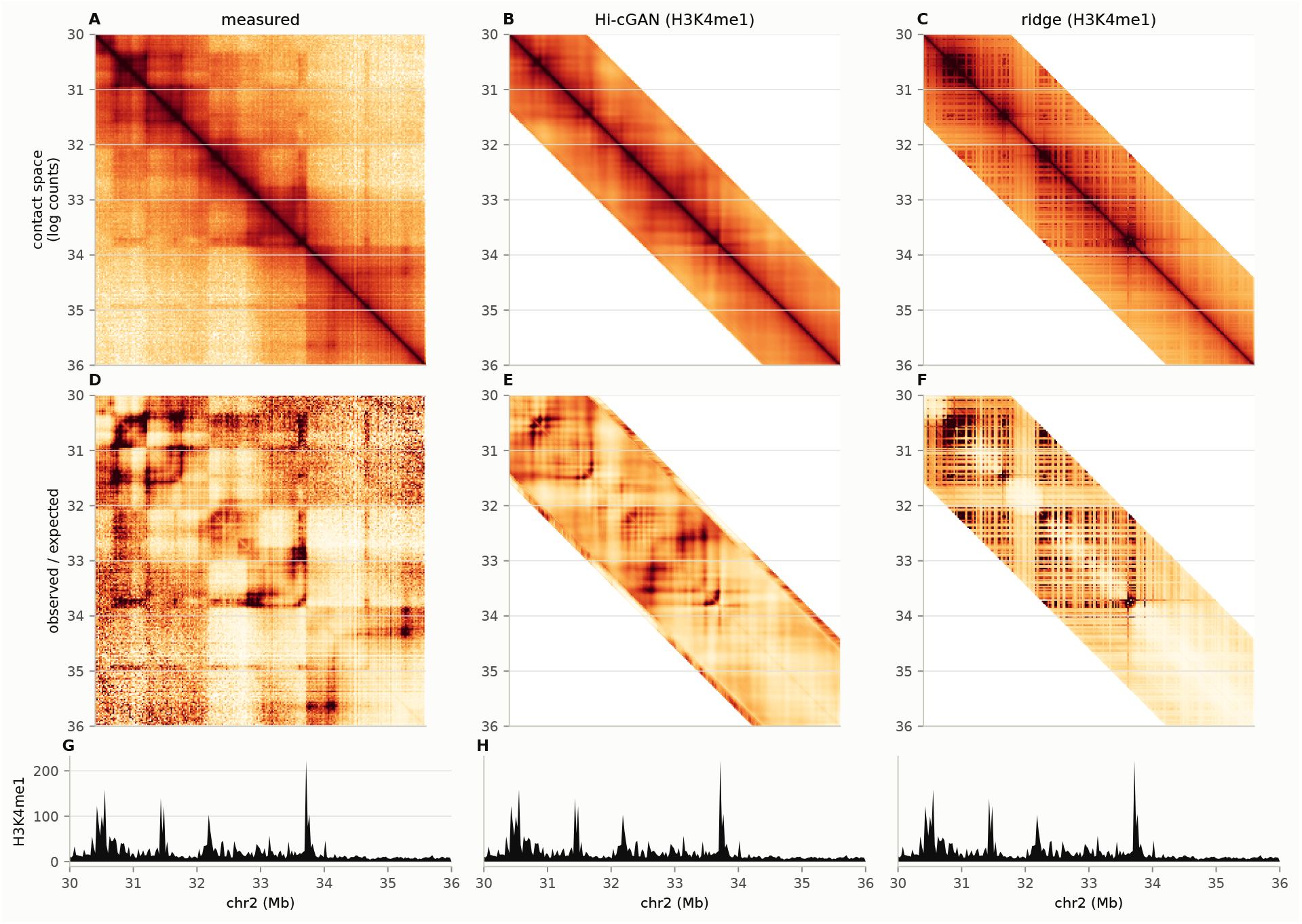
Measured, Hi-cGAN and the linear baseline over the same region. Chromosome 2, 30 to 36 Mb, at 25 kb. Top row in contact space on a shared logarithmic colour scale taken from the measured map; middle row after dividing each diagonal by its mean, which removes the distance decay and leaves local structure; bottom row the H3K4me1 track both predictors read. The predictions fill the 1.6 Mb band their window reaches and are blank beyond it. The lattice in panel F is the signature of a model whose prediction at a pixel depends only on the track at the two bins it joins.

## S16 Contribution of the adversarial term

The contribution of each of the three terms of the objective was measured by ablation. Three losses were trained under conditions identical in every other respect: the full objective, the same without the adversarial term, and the same again without the total variation term as well. Each was run with three random seeds, giving nine models. Training is on chromosome 1, validation on 19 and scoring on the held-out chromosome 4, at 25 kb with a window of 64 and H3K4me1 as the input, for 100 epochs. One chromosome per role keeps nine runs affordable; the resulting scores are lower than the whole-genome ones of the method battery (Table S3) and are not comparable with them, since only the differences between the three losses are read from them.

No measure separates the three (Table S18; the nine individual models are in Table S19). The largest spread between conditions is 0.012 on insulation correlation against a seed-to-seed standard deviation of 0.010, and on HiCRep the conditions span 0.003 against a spread of 0.009. Removing the total variation term also changes nothing, as expected at *λ*_TV_ = 10*^−^*^10^. The conditions come closest to separating on the contrast slope, and there the difference runs against the usual argument for adversarial training: the slope is lowest with the discriminator, 0.0485 against 0.0502 without it.

**Figure S18:**
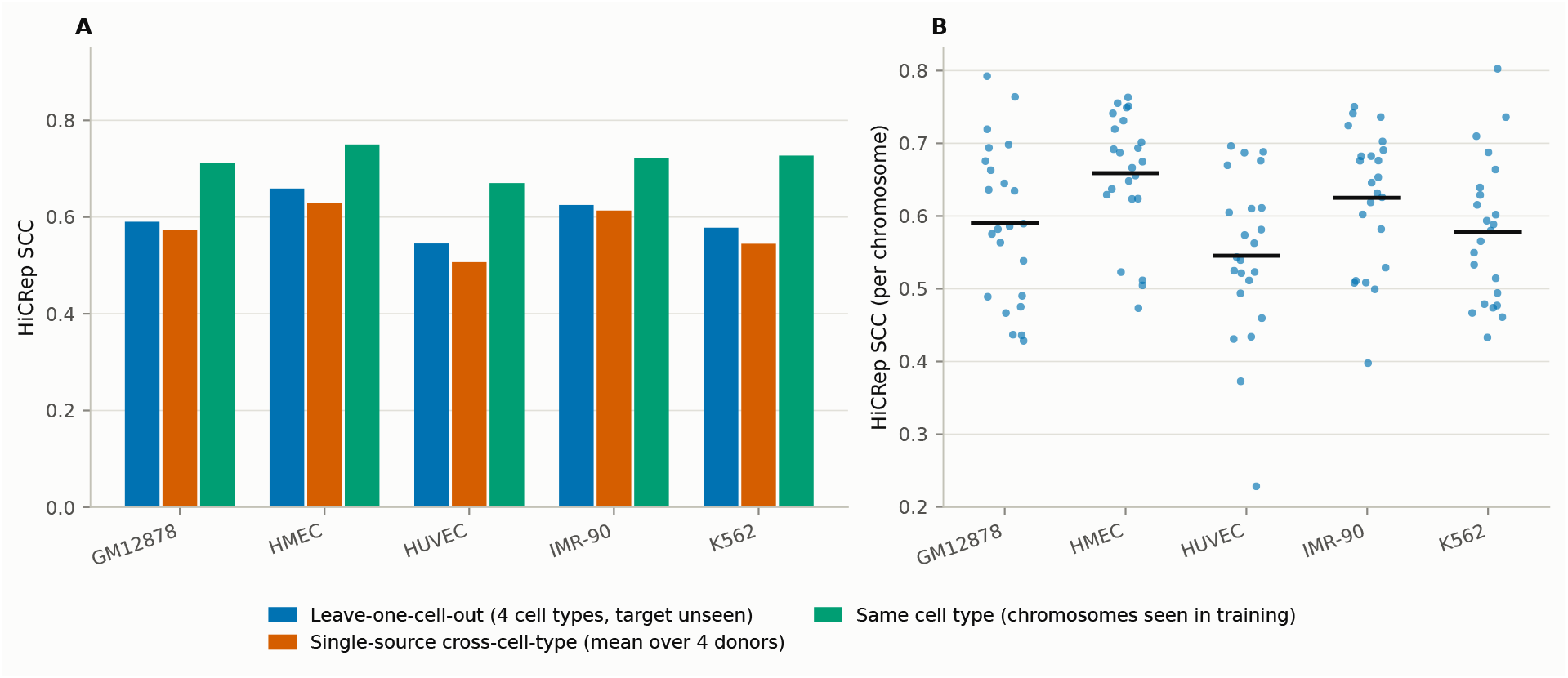
Leave-one-cell-out generalisation at 25 kb. (A) A model trained on four cell types and applied to the fifth, against transfer from a single source and against the same cell type the model was trained on. (B) the per-chromosome distribution behind the leave-one-cell-out means, 23 chromosomes per cell type. H3K27ac and CTCF as input, window 256, 100 epochs.

The result is bounded to the tested configuration. At a window of 64 a PatchGAN output unit spans 34 of the 64 bins against 70 of 512 at the largest window (Table S2), so what is tested is a discriminator judging half the submatrix at a time, not the far more local critic it becomes at 512, where the comparison was not run. Within this configuration, which is the one behind the GM12878 results at 25 kb, a user gains nothing measurable from the discriminator and can train the generator alone, which also removes the discriminator’s share of the cost per epoch. Epiphany reports the same comparison for its own architecture, where the adversarial loss lowered correlation and was retained for the sharpness it gave Yang et al. (2023).

## S17 The adversarial ablation, model by model

The main text reports the mean over three seeds per loss and the spread across them. We give the nine underlying models individually here, so that the overlap between conditions can be read directly rather than inferred from a mean of three.

**Figure S19:**
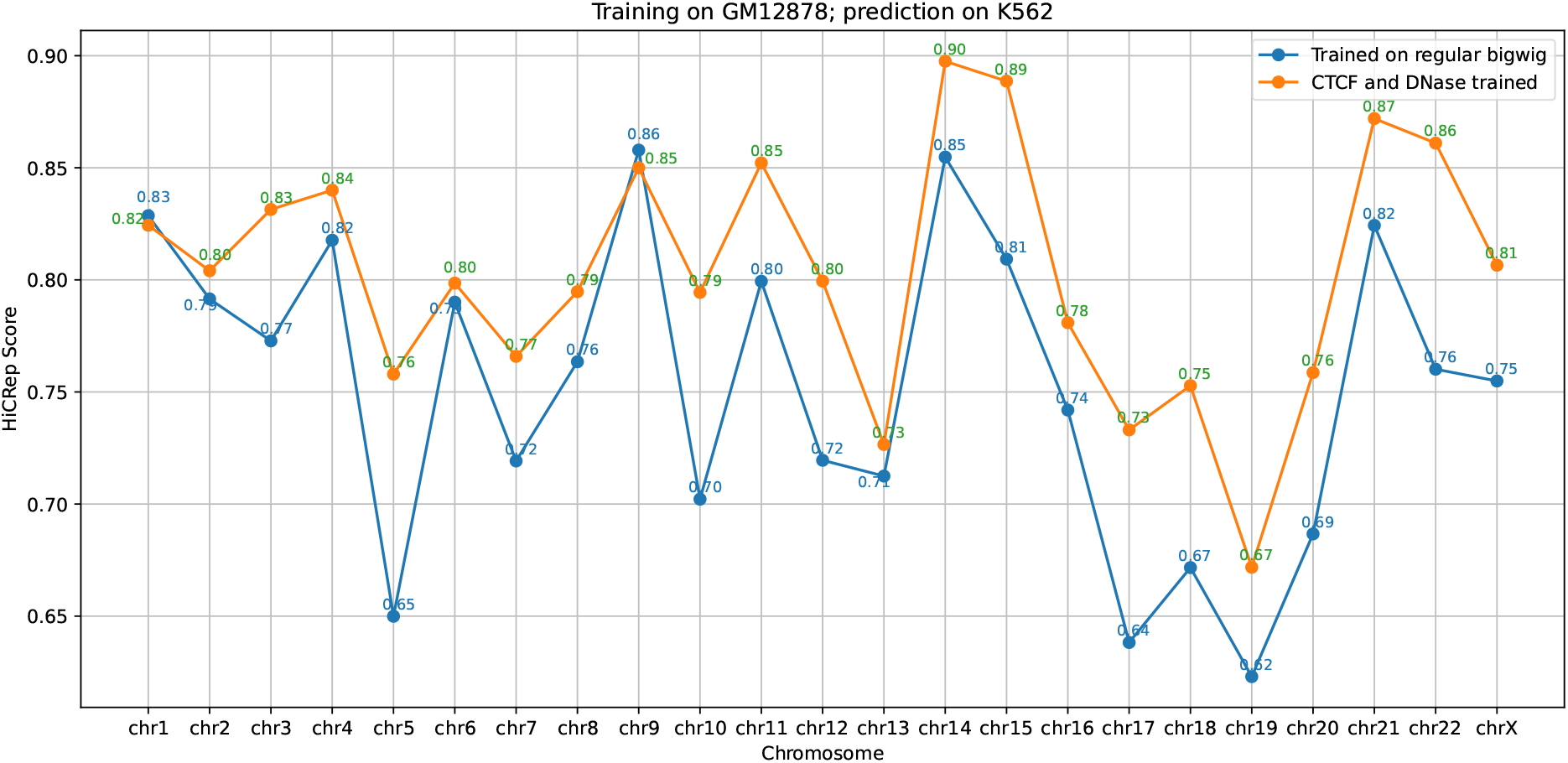
Predicting K562 from a GM12878 model. Training on all GM12878 chromosomes except chromosome 19, which is used for validation; prediction of every K562 chromosome. Two input configurations are shown: the full set of 14 chromatin binding proteins, and CTCF with DNase alone. Window size 64, 10 epochs, validated against the data of Rao et al. (2014).

We draw this conclusion for one regime only: one bin size, one input track, one chromosome in each role and a window of 64, at which a PatchGAN output unit spans 34 of the 64 bins (Table S2). At the window of 512 used at 10 kb and below each output unit spans 70 bins, about an eighth of the window, so the discriminator covers a far smaller share of the submatrix at a time. We did not repeat the experiment there, an epoch costing some forty times as much.

## S18 Prediction across cell types

The five-by-five transfer matrix and the leave-one-cell-out result are in the main text. This section records the single-pair experiment behind them.

A model was trained on all GM12878 chromosomes except chromosome 19, which was used for validation, and every chromosome of K562 was then predicted from K562 input tracks (Figure S19). With CTCF and DNase alone, 15 of 22 chromosomes reach a HiCRep score of at least 0.8; with the full set of 14 chromatin binding proteins, 7 do. No K562 data entered training, so the overfitting seen when the training cell type is predicted is absent here.

Recovering cell-type-specific structure, rather than the conserved part of the conformation, would require a system in which the conformation genuinely differs from the source, such as cohesin depletion Rao et al. (2017).

## S19 Accuracy per chromosome and per epoch

For the GM12878 test chromosomes at 25 kb, read at every tenth training epoch, which chromosome is predicted matters more than how long the model is trained, and the two principal quality measures agree on this (Figures S20, S21a and S21b). Chromosome 22 reaches roughly 0.65 at every epoch and no other chromosome comes close, while chromosome 20 stays between 0.37 and 0.45. The average oscillates between 0.50 and 0.55 and peaks near epoch 40. The Pearson correlation of the HiCExplorer TAD separation score, computed on the predicted against the measured matrix, follows the same pattern, which is evidence that the effect is a property of the chromosome rather than of the measure (Figure S21).

**Figure S20:**
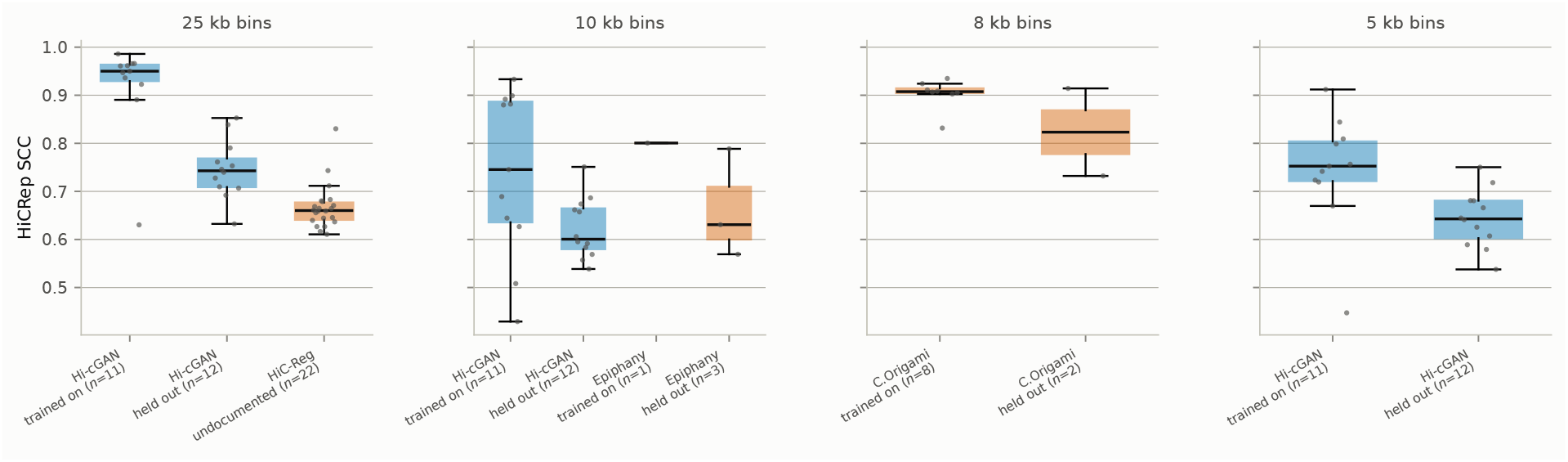
Per-chromosome HiCRep for every method at every resolution it exists at. One point per chromosome, split into the chromosomes each method trained on and those it did not, taken from each method’s own source. The split for the released HiC-Reg model is undocumented, so its 22 chromosomes are shown together. The gap between the groups is the memorisation a chromosome-level split leaves visible: 0.920 against 0.746 for Hi-cGAN at 25 kb. HiC-Spector and GenomeDISCO are Supplementary Figure S12.

**Figure S21:**
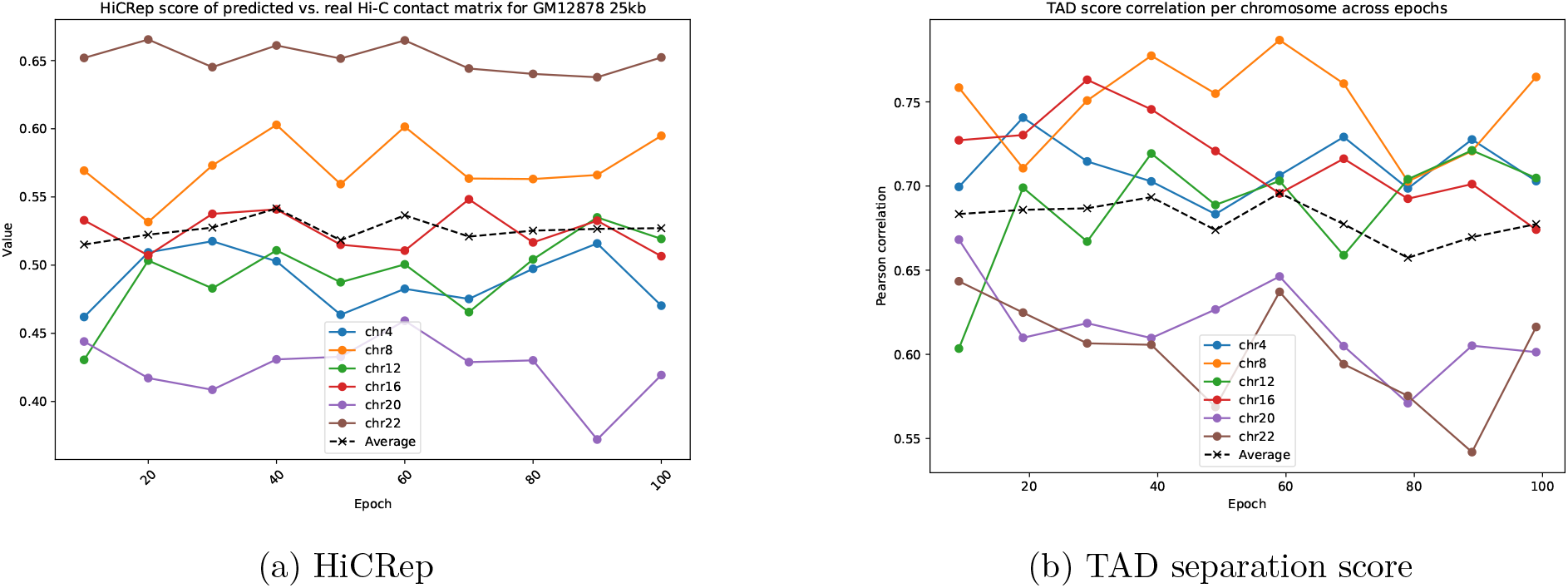
Accuracy per chromosome and training epoch. (a) HiCRep score for GM12878 on the test chromosomes against Rao 2014 data, using full bigwig tracks. (b) Pearson correlation of HiCExplorer TAD separation score on predicted against measured.

Table S20 quantifies the same ordering for the models trained on single chromosomes. Each chromosome is ranked among the 23 for every model, and the table gives the distribution of that rank. Chromosome 1 is the easiest to predict, with a mean rank of 4.9, and chromosome 19 the hardest, at 21.2 with a standard deviation of only 1.1, meaning it is close to last irrespective of what the model was trained on. Chromosome 19 has the highest gene density in the human genome, which is a plausible reason for it behaving differently from the rest.

**Table S18:** The adversarial term makes no measurable difference here. Mean over three seeds per loss, on held-out chromosome 4. The last row is the mean within-condition standard deviation across seeds. Every difference between the three losses is at or below it.

| Loss | HiCRep | Insul. | Bound. | Slope |
| --- | --- | --- | --- | --- |
| MAE only | 0.602 | 0.690 | 0.041 | 0.049 |
| MAE + TV | 0.605 | 0.690 | 0.035 | 0.050 |
| MAE + TV + adv. | 0.603 | 0.702 | 0.035 | 0.049 |
| seed spread (sd) | 0.009 | 0.010 | 0.006 | 0.001 |

**Table S19:** Every model of the adversarial ablation. Held-out chromosome 4, 25 kb, window 64, H3K4me1, 100 epochs, training on chromosome 1 and validation on 19. Columns are the HiCRep SCC, the Pearson correlation of the insulation profile, the Jaccard index of the called boundaries and the contrast slope. Every column’s between-condition range is covered by the within-condition spread, with the partial exception of the contrast slope discussed in the main text.

| Loss | Seed | HiCRep | Insul. | Bound. | Slope |
| --- | --- | --- | --- | --- | --- |
| MAE only | 1 | 0.5874 | 0.6811 | 0.0323 | 0.0488 |
|  | 2 | 0.6133 | 0.7024 | 0.0521 | 0.0499 |
|  | 3 | 0.6063 | 0.6872 | 0.0380 | 0.0491 |
|  | sd | 0.0134 | 0.0110 | 0.0102 | 0.0006 |
| MAE + TV | 1 | 0.6071 | 0.6978 | 0.0413 | 0.0515 |
|  | 2 | 0.6068 | 0.6860 | 0.0351 | 0.0501 |
|  | 3 | 0.6020 | 0.6860 | 0.0298 | 0.0488 |
|  | sd | 0.0029 | 0.0068 | 0.0057 | 0.0013 |
| MAE + TV + adv. | 1 | 0.5926 | 0.6917 | 0.0380 | 0.0479 |
|  | 2 | 0.6138 | 0.6978 | 0.0344 | 0.0494 |
|  | 3 | 0.6014 | 0.7175 | 0.0336 | 0.0484 |
|  | sd | 0.0107 | 0.0135 | 0.0023 | 0.0008 |

**Table S20:** Rank of each chromosome across the individual-chromosome models. Mean, standard deviation, minimum and maximum rank, sorted by mean rank.

|  | mean rank | std rank | min rank | max rank |
| --- | --- | --- | --- | --- |
| chr1 | 4.9 | 3.5 | 1 | 15 |
| chr5 | 6.6 | 4.7 | 1 | 18 |
| chr9 | 7.4 | 4.7 | 1 | 18 |
| chr8 | 8.4 | 4.5 | 1 | 15 |
| chr3 | 8.1 | 4.3 | 1 | 17 |
| chr11 | 8.1 | 4.5 | 1 | 14 |
| chr14 | 8.2 | 4.5 | 1 | 16 |
| chr7 | 8.8 | 4.2 | 1 | 20 |
| chr2 | 9.3 | 5.2 | 1 | 17 |
| chr13 | 9.6 | 6.1 | 1 | 19 |
| chr18 | 10.5 | 5.1 | 1 | 18 |
| chr15 | 11.3 | 6.7 | 1 | 21 |
| chr12 | 11.5 | 5.7 | 1 | 17 |
| chr4 | 12.2 | 4.8 | 1 | 19 |
| chr6 | 12.3 | 4.4 | 3 | 19 |
| chr16 | 12.8 | 5.4 | 1 | 20 |
| chr10 | 15.4 | 4.6 | 4 | 21 |
| chr22 | 16.4 | 7.7 | 1 | 23 |
| chr21 | 16.7 | 7.1 | 1 | 23 |
| chrX | 17.6 | 4.4 | 1 | 23 |
| chr17 | 17.7 | 4.7 | 1 | 22 |
| chr20 | 20.4 | 3.5 | 6 | 23 |
| chr19 | 21.2 | 1.1 | 19 | 23 |

## S20 Parameter sweeps

Each figure in this section is a sweep over one parameter of the training setup, with the per-chromosome detail: the choice of training chromosomes (Figure S22) and the form of the input, all fourteen tracks, their peak calls, or CTCF with DNase alone (Figure S25).

**Figure S22:**
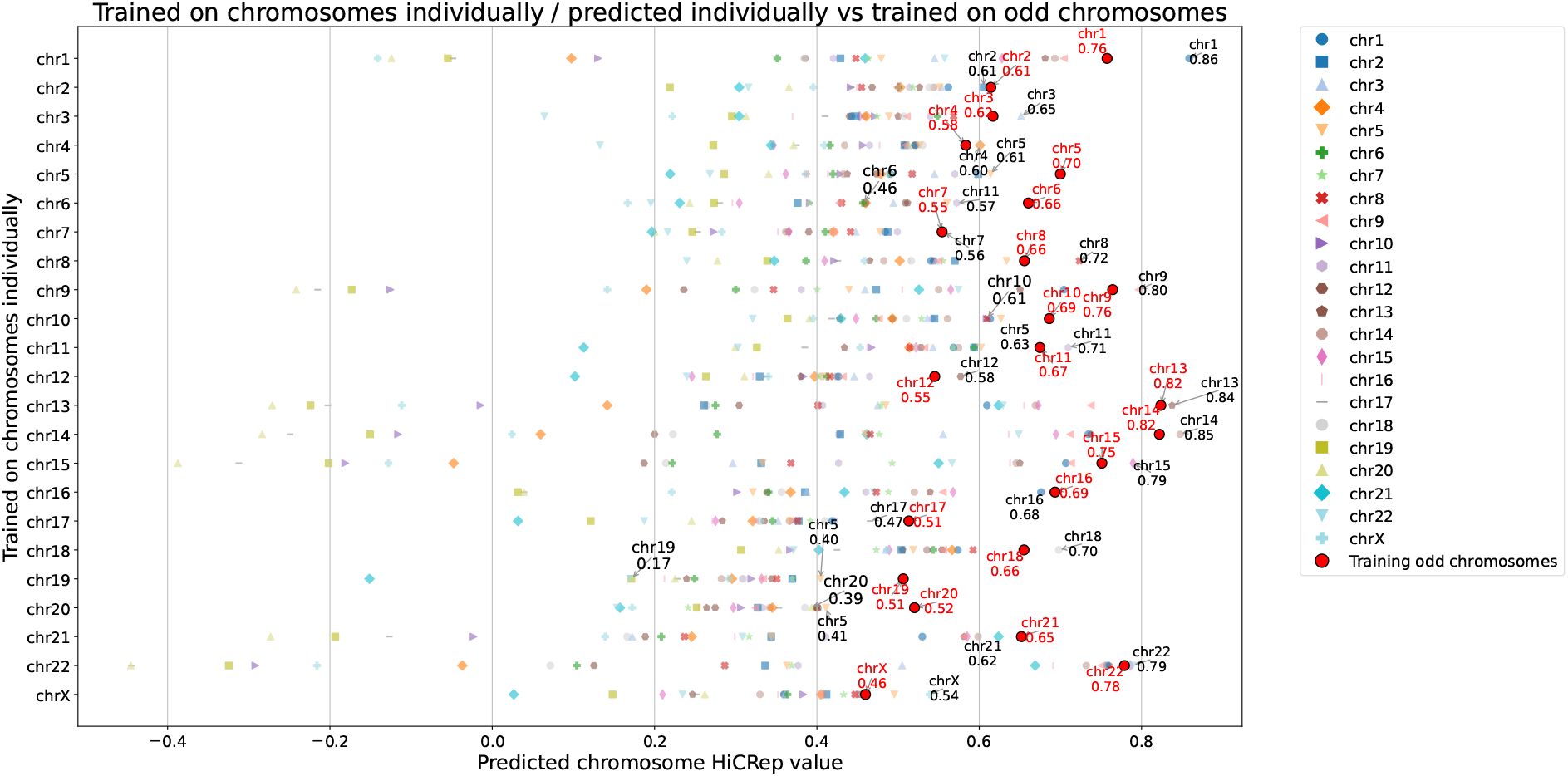
HiCRep scores of different chromosomes as training data. Training for individual chromosomes and validation on chromosome 2, except for training on chromosome 2 with validation on chromosome 1. Per individual trained chromosome, all chromosomes are predicted and the HiCRep score for 1 MB genomic distance is computed. The in red highlighted HiCRep scores are from a training on all odd chromosomes, and validation on chromosomes 2, 6, 10, 14 and 18. Window size of 128 used for all trainings and the full set of bigwigs.

**Figure S23:**
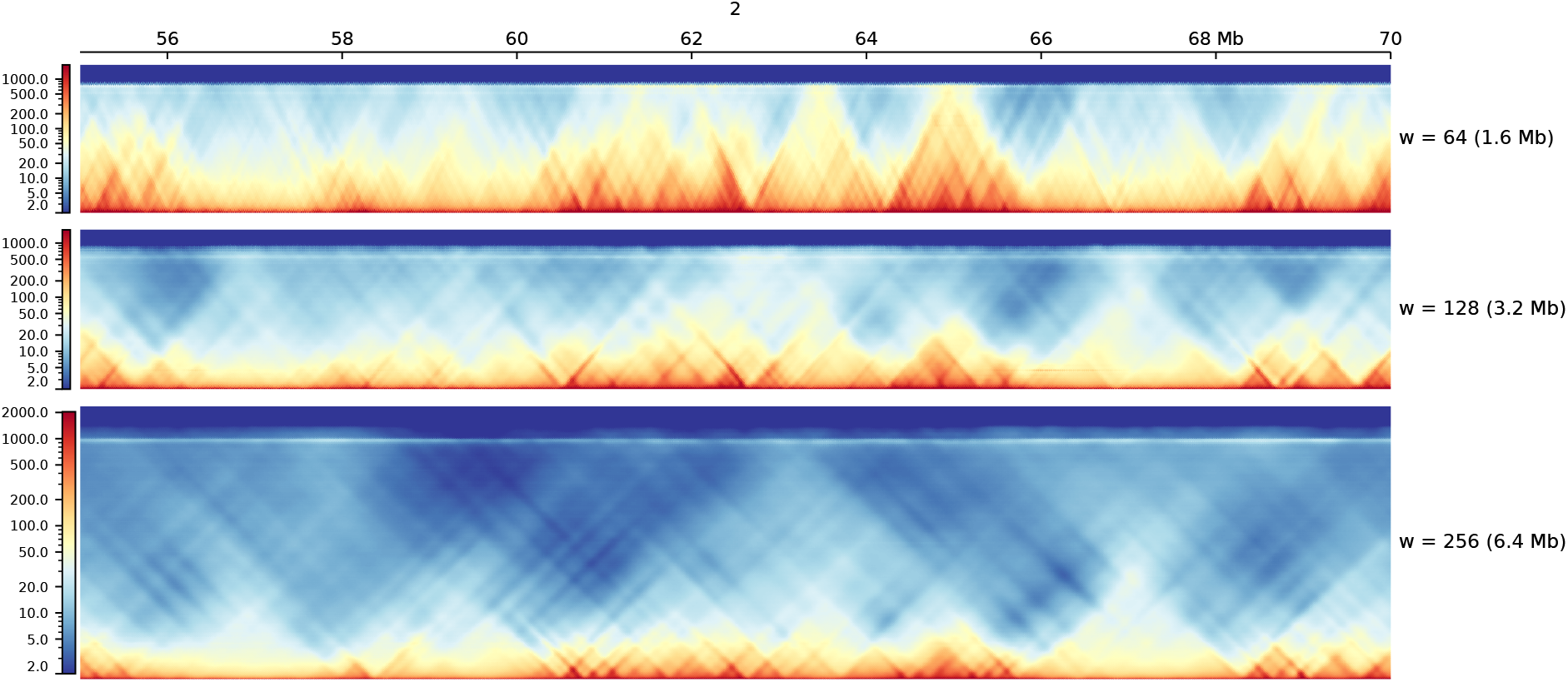
Prediction depth of the three windows at 25 kb. Chromosome 2, 55–70 Mb, predicted from H3K4me1 at windows of 64, 128 and 256 bins and drawn by pyGenomeTracks at a shared scale per megabase. The height of each track is the maximum genomic distance that window can predict, 1.6, 3.2 and 6.4 Mb.

**Figure S24:**
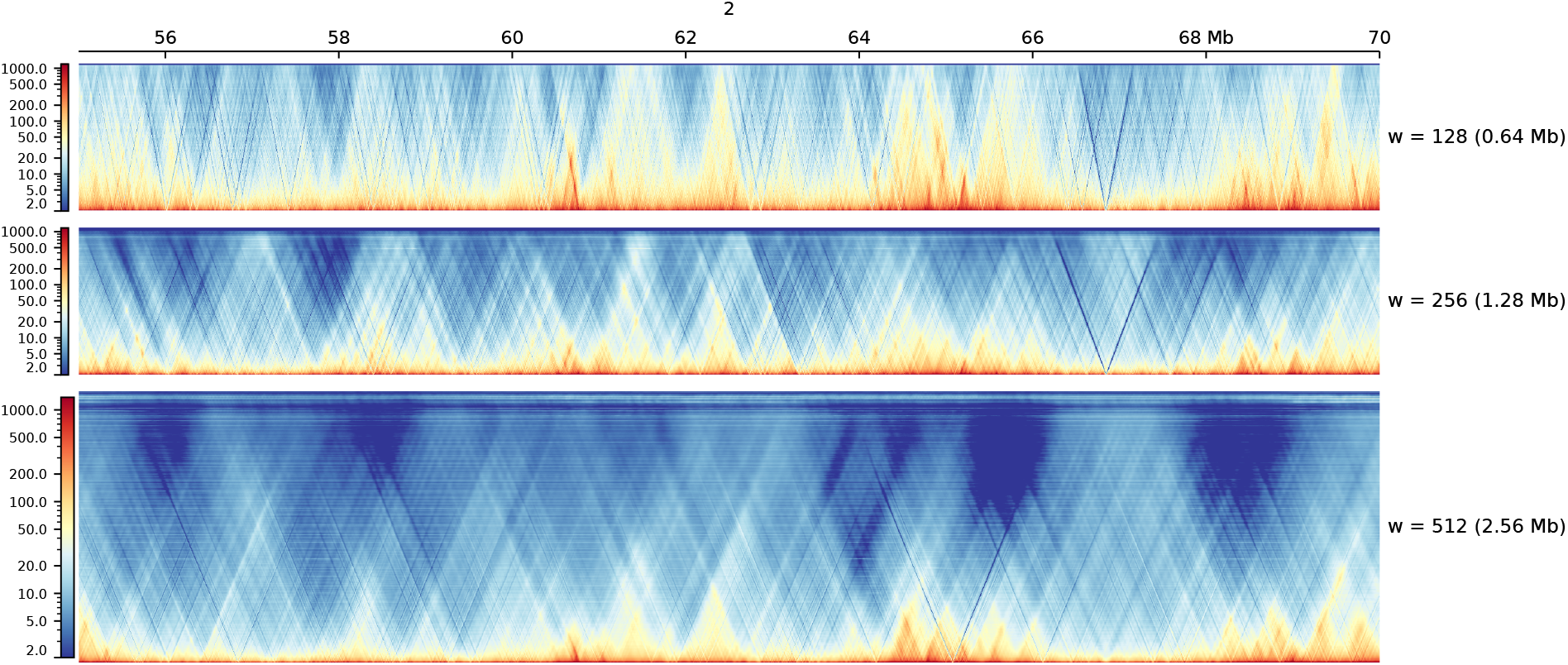
Prediction depth of the three windows at 5 kb. Chromosome 2, 55–70 Mb, predicted from CTCF at windows of 128, 256 and 512 bins, drawn as in Figure S23. The reaches are 0.64, 1.28 and 2.56 Mb.

## S21 Software environment and archived material

Every number in this article was produced under Python 3.11.9 with TensorFlow 2.15.0, cooler 0.10.3, HiCExplorer 3.7.3, NumPy 1.26.4, SciPy 1.14.0, pandas 2.2.2, scikit-learn 1.6.1 and Matplotlib 3.8.4. A specification file for the complete environment is archived with the data.

The archive also holds the scripts that produce every figure and table in this article, the held-out interval file, the per-window implementations of SCC, insulation agreement, GenomeDISCO and HiC-Spector, the script that validates the SCC against the hicrep reference implementation, and the random seeds used by the ablation.

## S22 Commands

The commands below are the ones used here, with the option names of the released software. hicTraining and hicPredict are installed on the path by the package; they are not scripts to be called with python. Both are also given as plain text with the archived data.

Training takes bigwig files and Hi-C matrices in cool format. The split is the one of the main text: the odd chromosomes except 19 for training, 19 for validation.

`hicTraining -tm gm12878_25kb.cool -tchroms 1 3 5 7 9 11 13 15 17 21 -tcp gm12878_bigwig/-vm gm12878_25kb.cool -vchroms 19 -vcp gm12878_bigwig/ -ws 64 -o trained_model_w64 - ep 100 --seed 1`

Several cool files may be given to -tm, in which case a bigwig folder is given to -tcp for each of them. Checkpoints are written every ten epochs as generator_000NN.keras.

Prediction reads a checkpoint and writes a cool file: `hicPredict --trainedModel trained_model_w64/generator_00099.keras -tcp gm12878_bigwig/ - pc 2 4 6 8 10 12 14 16 18 20 22 X -o gm12878_w64/ -mn predicted.cool –multiplier 1000 -ws 64 -b 25000`

**Figure S25:**
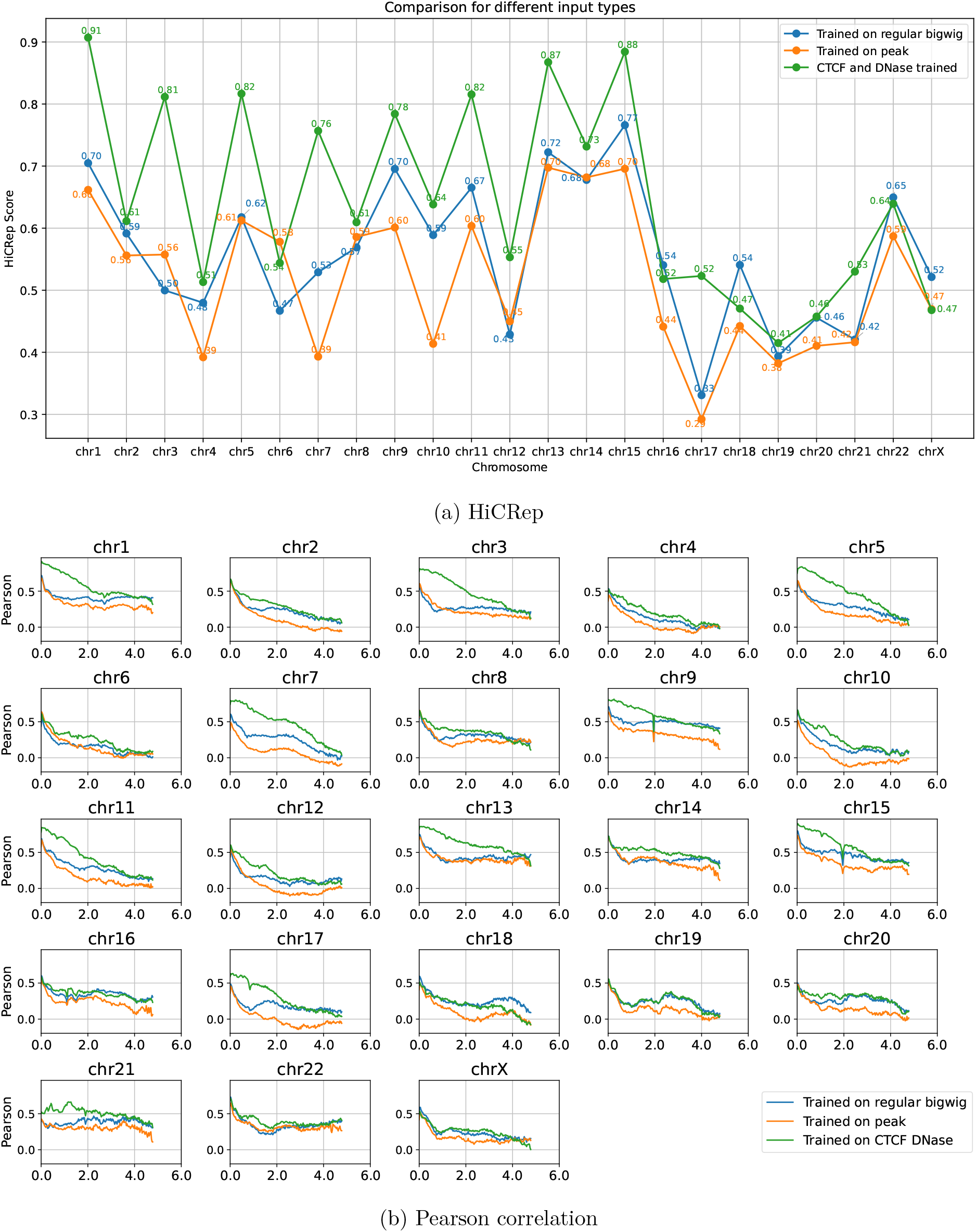
Predicted GM12878 with three input sets: the fourteen chromatin tracks, their peak calls, and CTCF with DNase alone. Window size 256, epoch 20, validated against the Rao 2014 data. (a) HiCRep score per chromosome, (b) Pearson correlation per chromosome and genomic distance.

